# Global and local adaptation to aridity in a desert plant *Gymnocarpos przewalskii*

**DOI:** 10.1101/2023.08.13.553124

**Authors:** Ruirui Fu, Yuxiang Zhu, Ying Liu, Zhaoping Yang, Ruisen Lu, Yingxiong Qiu, Martin Lascoux, Pan Li, Jun Chen

## Abstract

In order to thrive and survive plant species need to combine stability in the long term and rapid response to environmental challenges in the short term. The former would be reflected by global adaptation across species and the latter by pronounced local adaptation among populations of the same species. It remains unclear how much overlap is to be expected between the parts of the genome associated to these two contrasted adaptation processes. In the present study, we generated a high-quality genome and re-sequenced 177 individuals for *Gymnocarpos przewalskii*, an important desert plant species from North-West China, to detect local adaptation. To test for global adaptation to aridity at the molecular level we compared genomic data of 15 species that vary in their ability to withstand drought. A total of 118 genes were involved in global adaptation to aridity. Sixty-five *G. przewalskii* genes were shared across all xerophytic species, of which sixty-three were under stabilizing selection and two under directional selection. While 20% of *G. przewalskii* genome showed signatures of local adaptation to aridity during population divergence, only 13 of those genes were also under global adaptation. Hence, our results suggest that long-term stability is crucial for adaptation to extreme environmental stress but is only maintained in a small group of highly pleiotropic genes while a rapid response to recent changes elicits a genome-wide response, including gene family expansion. The overlap between the two evolutionary mechanisms appears limited.

## Introduction

Adaptation is intrinsically a highly dynamic and multi-layered process as it tracks the response of a complex system, the organism, to the environment, a no less complex and structured set of biotic and abiotic factors. The rate of change of these environmental factors will vary in both time and space and one would therefore expect to observe varying adaptive responses on both scales. In some extreme and constraining environments, adaptive traits can develop highly conserved patterns across distantly related organisms leading to ‘convergent or parallel evolution’ across species and global adaptation within species ^1^. Global adaptation within species simply means that there is a single optimum across the species range. Meanwhile, as most environmental factors are heterogenous in space, adaptive traits will also vary across the geographic distribution range of a species resulting in a pattern of local adaptation ^2^. We should therefore observe signatures of both global and local adaptation at the genomic level. Since a given environmental factor (e.g. drought or salinity) can lead to both global and local adaptation and since most adaptive traits are complex, i.e. controlled by a large number of loci, one would also expect to see some overlap. It is, however, unclear how different the genomic architectures of global and local adaptation are and how much overlap is to be expected between loci involved in global and local adaptation. In the present study, we focus on these two types of evolutionary responses to aridity. Aridity is a key environmental factor and, considering the ongoing global warming, understanding the genetic basis of adaptation to drought appears timely.

Arid zones are defined as regions where evaporation is greater than precipitation ^3^. Arid zones can also be characterized by a series of abiotic factors, including low and unpredictable precipitation, desiccating winds, extremely hot summer and cold winter, intensive UV-radiation and high soil salinity, that hinder or prevent the growth of living organisms ^4,5^. Today up to 40% of the land surface is arid but this percentage has fluctuated through Earth history. Following the steep increase of temperature under the last century, this percentage will certainly increase, affecting the life and survival of a large number of species ^3^. Aridity imposes a strong selective pressure on plant species and leads to an adaptive syndrome that comprises traits such as dwarf and succulent stems, waxy and thorny leaves, deep-spreading roots, and unique stomatal aperture regulation for balancing water and gas exchange ^6,7^. In the present study, we leverage newly developed genomic resources to study genomic adaptation to aridity in *G. przewalskii* and across a larger group of both xerophytic and non-xerophytic species. *G. przewalskii* belongs to Caryophyllaceae family, and is a keystone species found across stony deserts of the arid and semi-arid regions of northwest China and Mongolia. Its physiological characters, such as mucronate leaves, sunken stomata, thick and waxy cuticle, well developed palisade tissue, high ratio of root to stem, indicate high adaptation to life in a dry desert ^8^. Together with its distribution across a wide range of longitude and latitude, these characteristics should favor local adaptation and make *G. przewalskii* ideal for studying the molecular mechanisms of adaptation to aridity at local scale.

To study global and local adaptation to aridity, we first established a high-quality reference genome of *G. przewalskii*. We then included this genome in a comparison of genomes of multiple xerophytic and non-xerophytic plant species to identify genes associated to adaptation to aridity at a global scale. The set of genes thus identified will be conservative and will include genes that are under convergent or parallel evolution but it will miss genes specific to *G. przewalskii* alone. For local adaptation, we used genome scans to identify molecular polymorphisms in *G. przewalskii* that are outliers for genetic differentiation among the main ancestral clusters of the species and are associated to variation in environmental factors. Finally, we used inferences of selection at global and local scales to estimate the relative parts played by directional, stabilizing and diversifying selection in shaping the genomic evolution of xerophytic plants.

## Results

### Genome sequences of xerophytic and non-xerophytic species

Global adaptation was studied using high-quality genome sequences of fifteen xerophytic and non-xerophytic plant species; fourteen genomes were collected from public databases (see **Table 1**) and the reference genome of *Gymnocarpos przewalskii* (2n=2×=40) was sequenced and *de novo* assembled with 385 Gb (∼166×) PacBio long reads and 208 Gb (∼90×) Illumina short reads. The final assembled *G. przewalskii* genome size was 2.095 Gb with contig N50 of 705.53 kb and scaffold N50 of 91.85 Mb. Approximately 94.4% (1.978 Gb) of the whole-genome assembly was anchored to 20 pseudo-chromosomes with length varying from 64.2 Mb to 178.5 Mb. Completeness was high as 88.4% of BUSCO ^9^ and 95.16% of CEGMA ^10^ were retrieved. In total, 51,990 protein-coding genes were predicted and 93.60% were functionally annotated based on sequence similarity search against known databases (see details in **Supplementary Methods S1**).

**Table 1.** Statistics of genome assembly and annotation of fifteen xerophytic and non-xerophytic species.

| Categories | Species | Genome sizes (Gb) | Number of protein-coding genes | Assembly quality | Contig N50 (kb) | Species description | Refs |
| --- | --- | --- | --- | --- | --- | --- | --- |
| OX | *Populus euphratica* | 0.57 | 36,606 | Chr. | 900 | Specifically adapted to extreme desert environments. | 74 |
|  | *Pugionium cornutum* | 0.55 | 31,412 | Chr. | 312 | Endemic to the Kubuqi Desert and Mu Us Desert in northwestern China. | 75 |
|  | *Ricinus communis* | 0.32 | 30,066 | Chr. | 8,960 | Grows on high-altitude tropical deserts of the African Plateau. | 76 |
|  | *Phaseolus acutifolius* | 0.66 | 27,095 | Chr. | 47 | Native to the Sonoran Desert, highly adapted to heat and drought. | 77 |
|  | *Cicer arietinum* | 0.53 | 28,269 | Chr. | 24 | Mainly cultivated in semi-arid environments. | 78 |
|  | *Lycium ruthenicum* | 2.27 | 32,711 | Scaff. | 16 | Distributed in northwest China, important to local plant communities in desertic regions. | 79 |
|  | *Pistacia vera* | 0.67 | 31,784 | Scaff. | 76 | Desert plant that is highly tolerant to drought and salt stresses. | 80 |
|  | *Ammopiptanthus nanus* | 0.82 | 37,188 | Contig | 2,760 | A rare broad-leaved shrub found in deserts and arid regions of Central Asia. | 81 |
| CX | *Hylocereus undatus* | 1.39 | 27,753 | Chr. | 580 | Originated from semi-desert, hot tropical regions of Latin America. | 82 |
|  | *Gymnocarpus przewalskii* | 2.10 | 51,990 | Chr. | 706 | Vastly distributed keystone species of stony deserts from arid and semi-arid regions of northwest China and Mongolia. | this study |
|  | *Simmondsia chinensis* | 0.89 | 23,490 | Chr. | 5,210 | Dioecious desert shrubs that are native to the Sonoran Desert and Baja California regions of North America. | 83 |
| CNX | *Beta vulgaris* | 0.57 | 27,421 | Chr. | 27 | An important crop of temperate climates, distributed throughout Asia Minor, the Mediterranean, and Europe. | 84 |
|  | *Fagopyrum tataricum* | 0.49 | 33,366 | Chr. | 551 | Originates from southwest China and currently grown in western China and the Himalayas, and also found in Japan, South Korea, Canada, and Europe. | 85 |
|  | *Spinacia oleracea* | 0.91 | 34,877 | Chr. | 1,800 | Native to central Asia, a temperate areas vegetable cultivated throughout the world. | 86 |
|  | *Dianthus caryophyllus* | 0.57 | 56,137 | Scaff. | 17 | One of the major floricultural crops in Japan and worldwide. | 87 |

note: OX refers to eight xerophytic species outside of the Caryophyllales order; CX refers to three xerophytic species in the Caryophyllales order; CNX refers to four non-xerophytic species in the Caryophyllales order, growing under less challenging water and heat conditions.

### The number of shared genes reflects global adaptation to aridity

To identify genes associated to global adaptation to aridity at the genome level and to characterize the way they evolved, we selected fifteen plant species with varying phylogenetic distances to *Gymnocarpos* and tolerance to aridity. Depending on both factors the fifteen species can be divided into three classes, CX (Caryophyllaceae, xerophytic), OX (non-Caryophyllaceae, xerophytic), and CNX (Caryophyllaceae, non-xerophytic) (see **M & M**, **Fig. 1a**). We then proceeded in three steps.

**Fig. 1.**
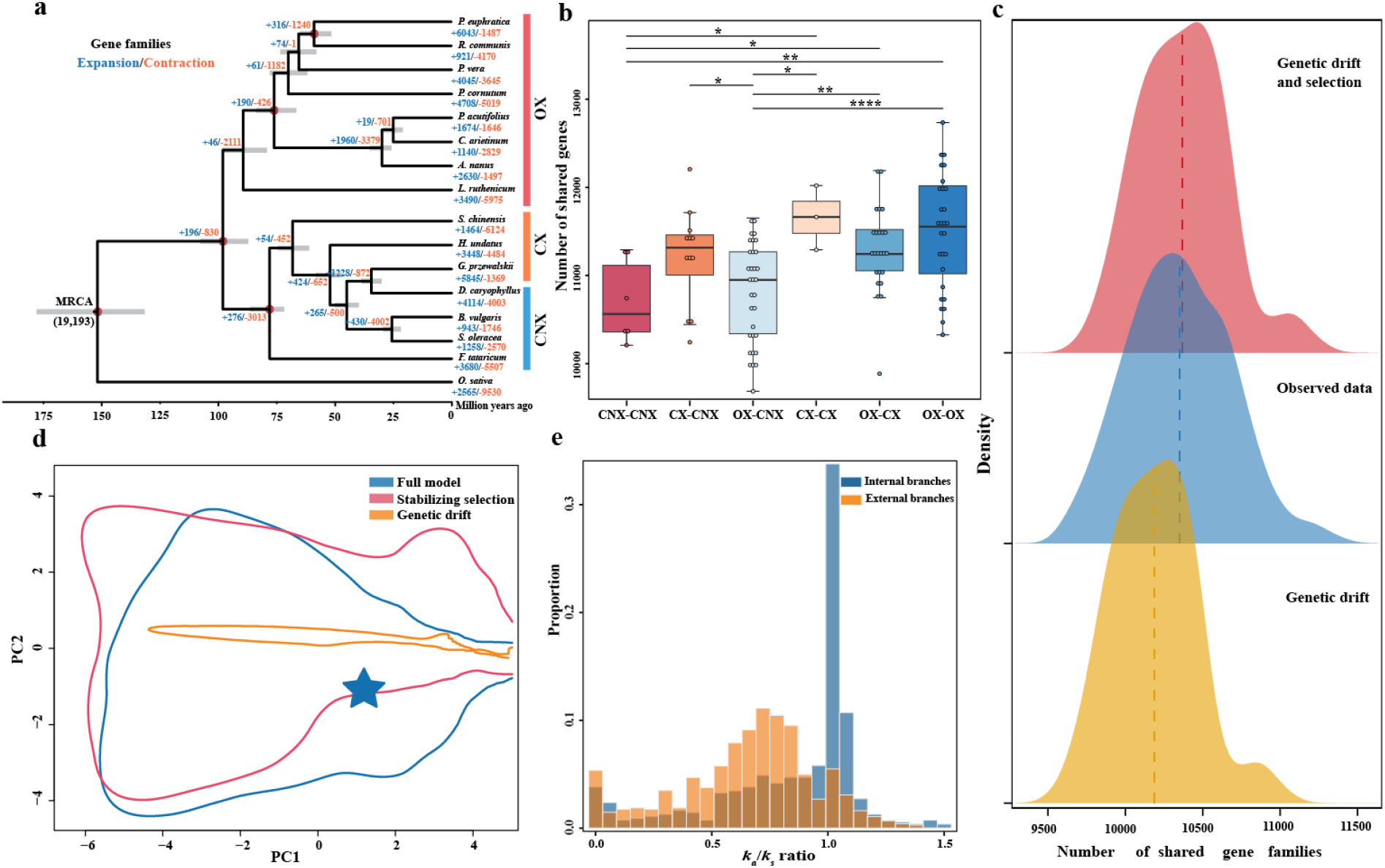
Tests for global adaptation in shared genes of xerophytic plant species. a) The phylogeny of non-xerophytic in Caryophyllales (CNX) and xerophytic species in and out of Caryophyllales (CX and OX, respectively). There are in total 19,193 gene families predicted in the most recent common ancestor (MRCA). The gray bars represent the 95% confidence interval of the divergence time of each node. The red dots represent calibration times. Numbers of expanded and contracted gene families are highlighted in blue and orange, respectively. b) The number of shared genes between pairs of species divided in six classes. Statistical significance was determined by a two-sided t-test (*p*-values: * ≤ 0.05, **≤ 0.01, ***≤ 0.001, **** ≤ 0.0001). Specifically, *p*-values = 0.019, 0.027, 0.006, 0.035, 0.045, 0.001, 4.6e-05 from top to bottom, respectively. c) The distributions of the observed and predicted number of shared genes without selection (bottom) and with selection (top). The x axis shows the number of shared gene families and is equally divided into 512 bins and the y axis gives the probability for each bin on the x axis. The dashed line represents the average number of shared genes. d) The goodness-of-fit tests for three models compared in OU simulations. The x axis and y axis represent the first two principal components obtained with PCA using *a priori* simulated summary statistics. The curves display envelopes containing 90% of the simulations. The star represents a projection of the observed summary statistics. The stabilizing selection model includes genetic drift and stabilizing selection. The full model includes genetic drift, stabilizing and directional selection. e) The distribution of *k_a_*/*k_s_* for “common xerophytic” genes that were shared by xerophytic species (≥ four species in CX and OX classes) but absent in all CNX species.

First, we tested if some genes are more likely to be maintained in species with similar life history traits and adapted to the same environment. The effects of pairwise phylogenetic branch length (genetic drift) and life history trait contrast (“xerophytic – xerophytic”, “xerophytic–non-xerophytic”, and “non-xerophytic–non-xerophytic”) were evaluated by the number of genes shared by any two species. Phylogenetic branch length was treated either as a fixed effect or a random effect in a mixed model. In both cases, life history trait contrast had a significant contribution (AIC = 1629 and 1605, *p*-value = 1.03e-6) and xerophytic species shared a significantly larger number of genes than non-xerophytic ones (mean = 11417 vs 10703, *p*-value = 4.6e-6). The effect of branch length and the effect of life history trait explained 23.2% and 17% of the total variance, respectively. The predicted number of genes shared by at least four plant species was significantly lower than the observed value (mean = 10189.34 vs. 10349.38, student t-test *p*-value = 5.4e-11) when the effect of life history trait was not accounted for (**Fig. 1c bottom**). However, the difference was not significant when the effect of life history trait was accounted for (mean 10368.74 vs. 10349.38, student t-test *p*-value = 0.43, **Fig. 1c top**).

Second, to further disentangle the role of natural selection from genetic drift on global adaptation to aridity, we used an Ornstein-Uhlenbeck (OU) process to model the evolution of the number of shared genes along the phylogeny. The OU process considers the effects of genetic drift, stabilizing selection, and directional selection ^11-13^. We applied the effects of stabilizing selection, and directional selection on the branches leading to xerophytic species. The mean numbers of shared genes between pairs of species were grouped in six classes (“OX – OX”, “CX – CX”, “OX – CX”, “CX – CNX”, “OX – CNX”, and “CNX – CNX”. **Fig. 1b**) and the standard errors were compared between observed and simulated data of three models: 1) a pure drift model, 2) a stabilizing selection model that includes genetic drift and stabilizing selection and 3) a full model that includes genetic drift, stabilizing and directional selection. Support for both the full model and the stabilizing selection model was significantly higher than for the pure drift model (B.F. = 35.2 and 19.3, respectively). No stronger support was found for the full model over the stabilizing selection one (B.F. = 1.8 and 0.62 using the ‘rejection’ method and ‘mnlogistic’ regression method, respectively) but the full model had the best performance in the goodness-of-fit test (**Fig. 1d**. See details in **Supplementary file 2**).

Third, to identify specific genes associated to global adaptation to aridity, we used *codeml* program in PAML ^14^ to calculate the ratio of the number of nonsynonymous substitutions per non-synonymous site to the number of synonymous substitutions per synonymous site, *k_a_/k_s_*, along different branches of the phylogeny. To be conservative, only 118 single copy orthologous genes were kept that were shared by xerophytic species (≥ four species in CX and OX classes) but absent in all CNX species (hereafter referred as the “common xerophytic” genes). The *k_a_/k_s_* ratios were computed for all 118 genes along all external and internal branches. There were in total 551 estimates of *k_a_/k_s_* along internal branches and 747 along external branches. The distribution of *k_a_/k_s_* estimates was bimodal for both internal and external branches but the two distributions otherwise differed (**Fig. 1e**, see **Fig. S14** for independent *k_a_* and *k_s_* distributions). The *k_a_/k_s_* distribution for the internal branches had a primary *k_a_/k_s_* peak around one, as expected under neutrality. Ninety-nine gene families showed a neutral signal (0.9≤ *k*_*a*_/*k*_*s*_ ≤ 1.1) and accounted for 49.9% (275 out of 551) of the total number of estimates along internal branches. A secondary *k_a_/k_s_* peak was close to zero (≤ 0.1) and included 29 gene families corresponding to 6.9% (38 out of 551) of the *k_a_/k_s_* along internal branches. Compared to internal branches, the *k_a_/k_s_* distribution for the external branches was shifted towards lower *k_a_/k_s_* values and had a primary *k_a_/k_s_* peak around 0.75. This primary peak included 94 gene families and 47.3% (353 out of 747) of the *k_a_/k_s_* along external branches. There was also a secondary peak close to zero that included 31 gene families and accounted for 6.2% (46 out of 747) of the *k_a_/k_s_* along external branches. For signals of directional selection, 15 gene families had *k_a_/k_s_* ≥ 1.2 along external branches and 16 families had *k_a_/k_s_* ≥ 1.2 along internal branches.

In summary, 118 genes were identified that were specific to xerophytic species. The distribution of *k_a_/k_s_* values for common xerophytic genes along internal branches suggests a predominantly neutral evolution, with a smaller fraction indicating stabilization selection. In contrast, stabilizing selection dominates on external branches. Therefore, most xerophytic genes evolved independently and were under convergent evolution. A much smaller fraction of genes, though, was already under stabilizing or directional selection on internal branches suggesting the presence of some parallel evolution.

### Population structure and demographic inference *in* G. przewalskii

Before evaluating local adaptation in *G. przewalskii*, we inferred the influence of neutral processes, i.e., genetic drift and migration, on population genetic divergence. 177 individuals from 26 populations were collected from the natural distribution range of the species and re-sequenced **(Fig. 2a**). *G. przewalskii* consists of four clearly delineated genetic clusters, namely Tarim, Hami, Hexi, and Alxa clusters (**Fig. 2b** and **Fig. S9**). Comparison of isolation by distance (IBD) and Isolation by environment (IBE) tests suggests that both genetic drift and local adaptation to environmental factors contributed to population divergence in *G. przewalskii* (Partial Mantel tests, coef. = 0.37, *p*-value =0.001 for the geographic distance and coef. = 0.27, *p*-value = 0.044 for the environmental distance, respectively. **Fig. 2c, d** and **Table 2**). In particular, geographic distance had a crucial influence on the divergence of the Tarim cluster from other clusters (**Fig. 2e**). This was confirmed by an effective migration surface (EEMS) analysis that revealed a major migration barrier centered around the Lop Nur and separating the Tarim populations from other populations (**Fig. 2f**).

**Fig. 2.**
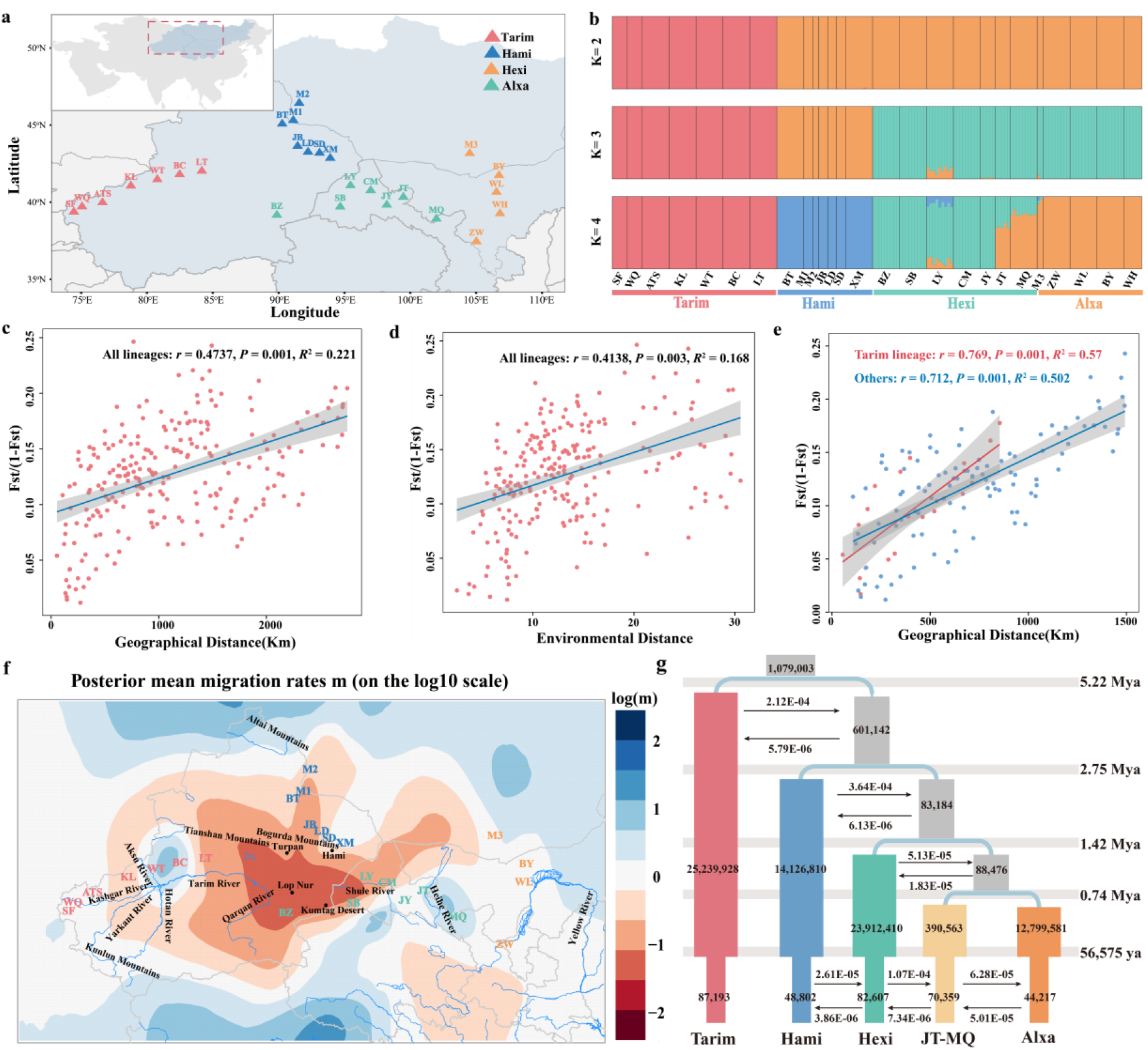
The population clustering and divergence of *G. przewalskii*. a) The distribution range (shaded) and population sampling locations (triangles). b) The results of ADMIXTURE analyses (*K* from 2 to 4). c-e) The Mantel tests of genetic divergence on geographic and environmental distances, respectively. c), d) for all populations and e) for the Tarim lineage and the three other lineages, separately. Shaded areas represent 95% confidence intervals. f) EEMS indicates that the main barrier to gene flow is centered on Lop Nor and separates the Tarim lineage from others. g) The best-supported demographic scenario estimated by fastsimcoal2. The black arrows show migration rates between lineages of *G. przewalskii*. The best point estimates of effective population sizes, divergence times and migration rates are shown.

**Table 2.**
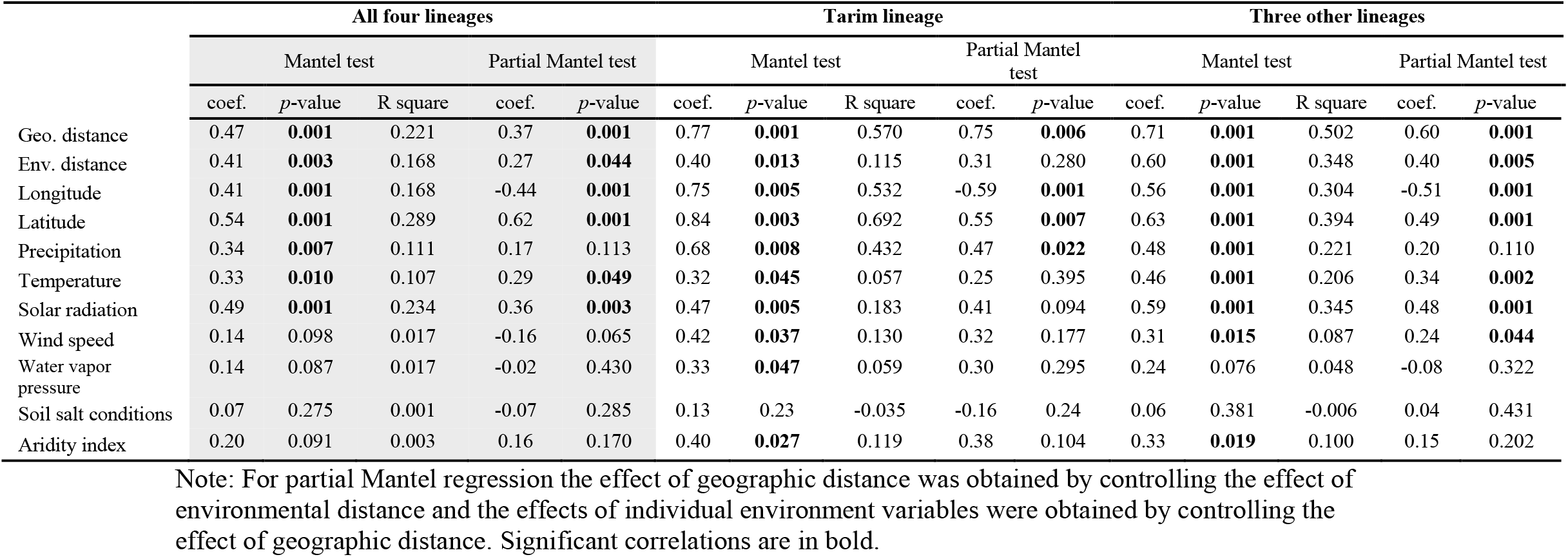
Results IBD and IBE using the Mantel test and partial Mantel test for the correlation between environmental distances, geographical distances and genetic distance from the cluster of Tarim and three other lineages (including Hami, Hexi, and Alxa).

|  | All four lineages |  |  |  |  | Tarim lineage |  |  |  |  | Three other lineages |  |  |  |  |
| --- | --- | --- | --- | --- | --- | --- | --- | --- | --- | --- | --- | --- | --- | --- | --- |
|  | Mantel test |  |  | Partial Mantel test |  | Mantel test |  |  | Partial Mantel test |  | Mantel test |  |  | Partial Mantel test |  |
|  | coef. | p-value | R square | coef. | p-value | coef. | p-value | R square | coef. | p-value | coef. | p-value | R square | coef. | p-value |
| Geo. distance | 0.47 | **0.001** | 0.221 | 0.37 | **0.001** | 0.77 | **0.001** | 0.570 | 0.75 | **0.006** | 0.71 | **0.001** | 0.502 | 0.60 | **0.001** |
| Env. distance | 0.41 | **0.003** | 0.168 | 0.27 | **0.044** | 0.40 | **0.013** | 0.115 | 0.31 | 0.280 | 0.60 | **0.001** | 0.348 | 0.40 | **0.005** |
| Longitude | 0.41 | **0.001** | 0.168 | -0.44 | **0.001** | 0.75 | **0.005** | 0.532 | -0.59 | **0.001** | 0.56 | **0.001** | 0.304 | -0.51 | **0.001** |
| Latitude | 0.54 | **0.001** | 0.289 | 0.62 | **0.001** | 0.84 | **0.003** | 0.692 | 0.55 | **0.007** | 0.63 | **0.001** | 0.394 | 0.49 | **0.001** |
| Precipitation | 0.34 | **0.007** | 0.111 | 0.17 | 0.113 | 0.68 | **0.008** | 0.432 | 0.47 | **0.022** | 0.48 | **0.001** | 0.221 | 0.20 | 0.110 |
| Temperature | 0.33 | **0.010** | 0.107 | 0.29 | **0.049** | 0.32 | **0.045** | 0.057 | 0.25 | 0.395 | 0.46 | **0.001** | 0.206 | 0.34 | **0.002** |
| Solar radiation | 0.49 | **0.001** | 0.234 | 0.36 | **0.003** | 0.47 | **0.005** | 0.183 | 0.41 | 0.094 | 0.59 | **0.001** | 0.345 | 0.48 | **0.001** |
| Wind speed | 0.14 | 0.098 | 0.017 | -0.16 | 0.065 | 0.42 | **0.037** | 0.130 | 0.32 | 0.177 | 0.31 | **0.015** | 0.087 | 0.24 | **0.044** |
| Water vapor pressure | 0.14 | 0.087 | 0.017 | -0.02 | 0.430 | 0.33 | **0.047** | 0.059 | 0.30 | 0.295 | 0.24 | 0.076 | 0.048 | -0.08 | 0.322 |
| Soil salt conditions | 0.07 | 0.275 | 0.001 | -0.07 | 0.285 | 0.13 | 0.23 | -0.035 | -0.16 | 0.24 | 0.06 | 0.381 | -0.006 | 0.04 | 0.431 |
| Aridity index | 0.20 | 0.091 | 0.003 | 0.16 | 0.170 | 0.40 | **0.027** | 0.119 | 0.38 | 0.104 | 0.33 | **0.019** | 0.100 | 0.15 | 0.202 |

Note: For partial Mantel regression the effect of geographic distance was obtained by controlling the effect of environmental distance and the effects of individual environment variables were obtained by controlling the effect of geographic distance. Significant correlations are in bold.

Demographic inference based on the joint site frequency spectrum (SFS) indicated that the Tarim cluster split from others about 5.22 Mya (**Fig. 2g**). After the Hami lineage diverged from others about 2.75 Mya, ancestral migration also ceased between the Tarim and the three other clusters, reinforcing its isolation. The Hexi and Alxa clusters diverged more recently, about 1.42 Mya. Gene flow between the Hami, Hexi, and Alxa clusters persisted at a slightly higher rate from west to east (2.61e-5 ∼ 1.07e-4) than in the opposite direction (3.86e-6 ∼ 5.01e-5). Population size contractions were identified in all four clusters around 56 Kya (See **Table S17** for results of model comparison and **Table S18** for parameter estimates). More detailed results can be found in **Supplementary Methods S3**.

### Local adaptation

To assess local adaptation, we first calculated the X^t^X statistic ^15^, an *F*_ST_-like statistic explicitly accounting for demographic history, that allows the identification of SNPs with higher levels of differentiation than would be expected under neutrality. A total of 174,675 SNPs deviated from neutral expectations (0.62% of the total number of SNPs investigated). Second, among the outlier SNPs identified with the X^t^X statistic, we tested for association between allele frequency and environment variables (**Fig. 3a**). In total, 146,515 SNPs (0.52% of the total number of SNPs) were significantly associated to one or more environment variables. The largest proportion of SNPs was associated with water vapor pressure (68.5%, 100,297 out of 146,515). In addition, 36,731 SNPs were associated to the aridity index, 31,153 SNPs to solar radiation, 33,439 SNPs to latitude, 30,242 SNPs to longitude, 18,972 SNPs to wind speed, 17,875 SNPs to precipitation, 11,282 SNPs to temperature, and 8,069 SNPs to salinity. For all climate variables, climate-related SNPs were significantly enriched in the 5% upper tail of the X^t^X distribution (*P* < 2.2e-16), with a fold-enrichment ranging from 1.29 for longitude to 2.43 for salinity.

**Fig. 3.**
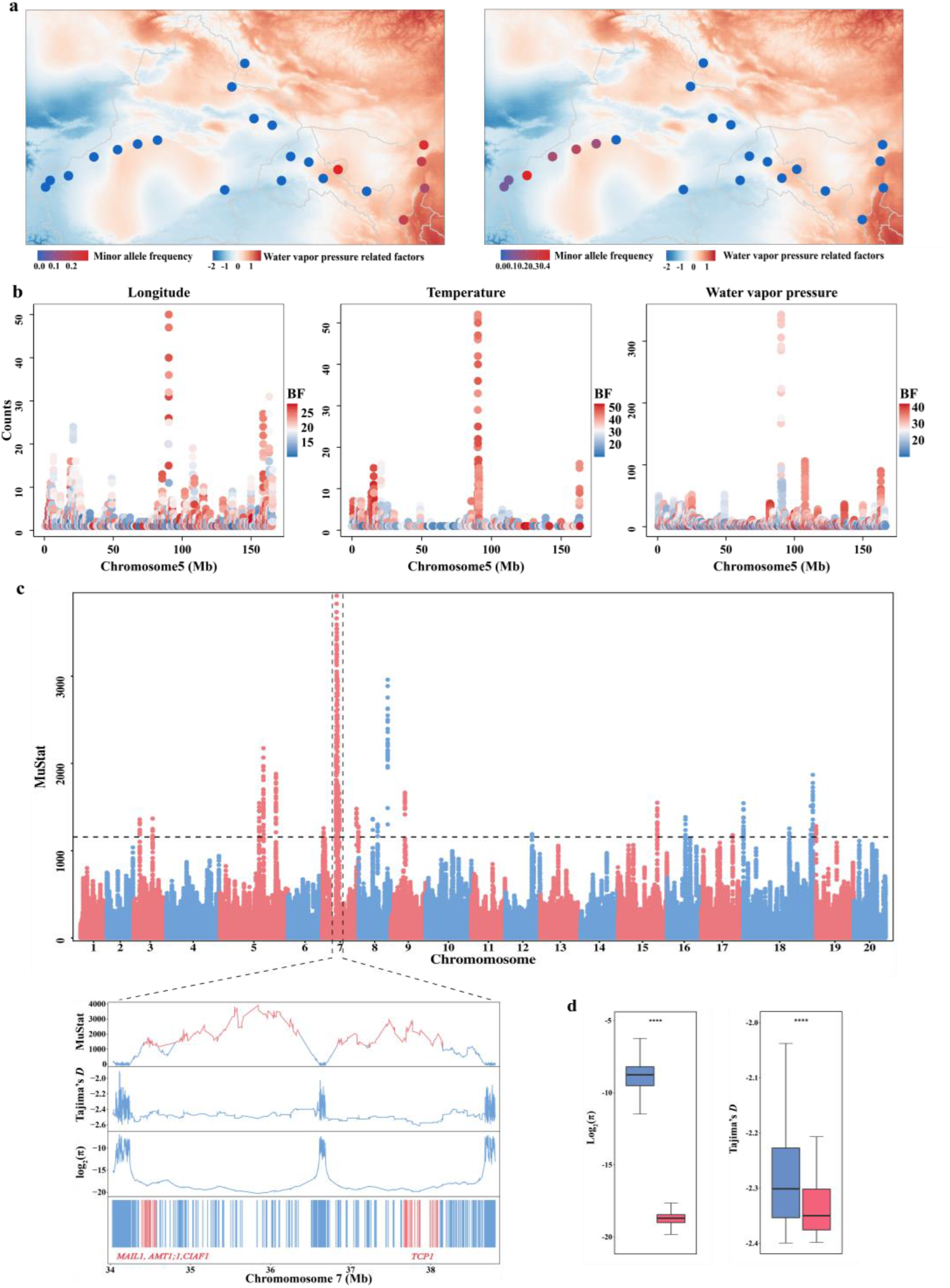
Genomic regions under local adaptation and selective sweeps for *G. przewalskii* populations. a) Two examples of candidate SNPs with allele frequency significantly correlated to water vapor pressure related factors (left for gene *CB5-E* and right for gene *UBQ1*). The background colors illustrate the distribution of environment variables and the colored dots illustrate the minor allele frequency. b) Chr.5 is enriched for candidate SNPs correlated to longitude, temperature and water vapor pressure related factors. c) The distribution of μ-statistic for selective sweeps across the reference genome. The horizontal dashed line marks the cutoff of μ-statistic (99.999% quantile). The μ-statistic, Tajima’s *D* and π distributions of the two largest windows of selective sweeps (Chr.7: 34.3 Mb - 36.6 Mb and Chr.7: 36.7 Mb - 38.3 Mb) are highlighted in the panels below. d) Genetic diversity (π) and Tajima’s *D* for candidate regions of selective sweeps (red boxes) and the genomic average values (blue boxes). Statistical significance was determined by two-sided student’s t-test. *p*-value <2.2e-16 for π and 4.03e-6 for Tajima’s *D*.

Environment-associated SNPs were randomly scattered across the whole genome except for region 89.89 Mb ∼ 90.46 Mb on chromosome 5 (Chr.5) that contained 501 SNPs, 12.58-fold as many as the average number of correlated SNP per Mb across the whole genome (Fisher’s exact test *p*-value < 2.2e-16) (**Fig. 3b and Fig. S15**). Interestingly, the number of SNPs located in the 89.89 Mb ∼ 90.46 Mb was positively correlated with linkage disequilibrium, estimated by *r*^2^ (Pearson’s correlation test *P* < 1.01×10^-10^, *ρ* = 0.73).

Up to 41.1% of all environment-associated SNPs were in transposable elements and tandem repeats (18.4% in Copia and 8.9% in Gypsy TE families). Approximately 14.8% were located in 10,516 protein-coding genes, i.e. approximately 20% (10,516 / 51,990) of all protein-coding genes (**Fig. S16**) and 8.0% were in proximal gene flanking regions. For all climate variables, nonsynonymous variants were significantly enriched among the loci strongly correlated with climate (*p*-value from 7.04e-18 to 8e-4), with fold-enrichment ratios ranging from 1.15 for aridity index to 1.38 for temperature (**Fig. S17**). Taken together, these results suggest that our scan effectively detects adaptive alleles.

### Selective sweeps

In total, 755 signatures of selective sweeps were identified (13.8 Mb; 0.7 % of the genome), of which 672 (89%) were located on Chr.5, Chr.7, Chr.8 and Chr.18. In particular, 439 regions (58%) were on Chr.7, 12.8-fold as many as the average number of selective sweeps across the whole genome (with the window size rescaled by the chromosome/genome length, Fisher’s exact test *p*-value < 2.2e-16), including the two longest regions (Chr.7: 34.3 Mb - 36.6 Mb and Chr.7: 36.7 Mb - 38.3 Mb, see **Fig. 3c**). 102 protein-coding genes were identified in all selective sweeps. Compared to the genome-wide average level, swept regions harbored significantly lower nucleotide diversity (mean 2.77 ×10^-6^ versus 2.79 ×10^-3^, student t-test *p*-value < 2.2e-16) as well as lower Tajima’s *D* (mean -2.453 versus -2.343, student t-test *p*-value = 4.03e-6, **Fig. 3d**).

### Relationship between different types of natural selection

In summary, comparison across species yielded 63 genes under stabilizing selection and two genes in *G. przewalskii* under directional selection. We also identified 10,516 genes putatively under local adaptation since they were X^t^X-outlier and their allele frequencies were significantly associated to environmental variables. 20.6% (13/63) of the genes under stabilizing selection were also involved in local adaptation while no genes under directional selection overlapped with local adaptation. Interestingly, no genes found to have experienced a selective sweep overlapped with other types of signatures of natural selection (**Fig. 4a**).

**Fig. 4.**
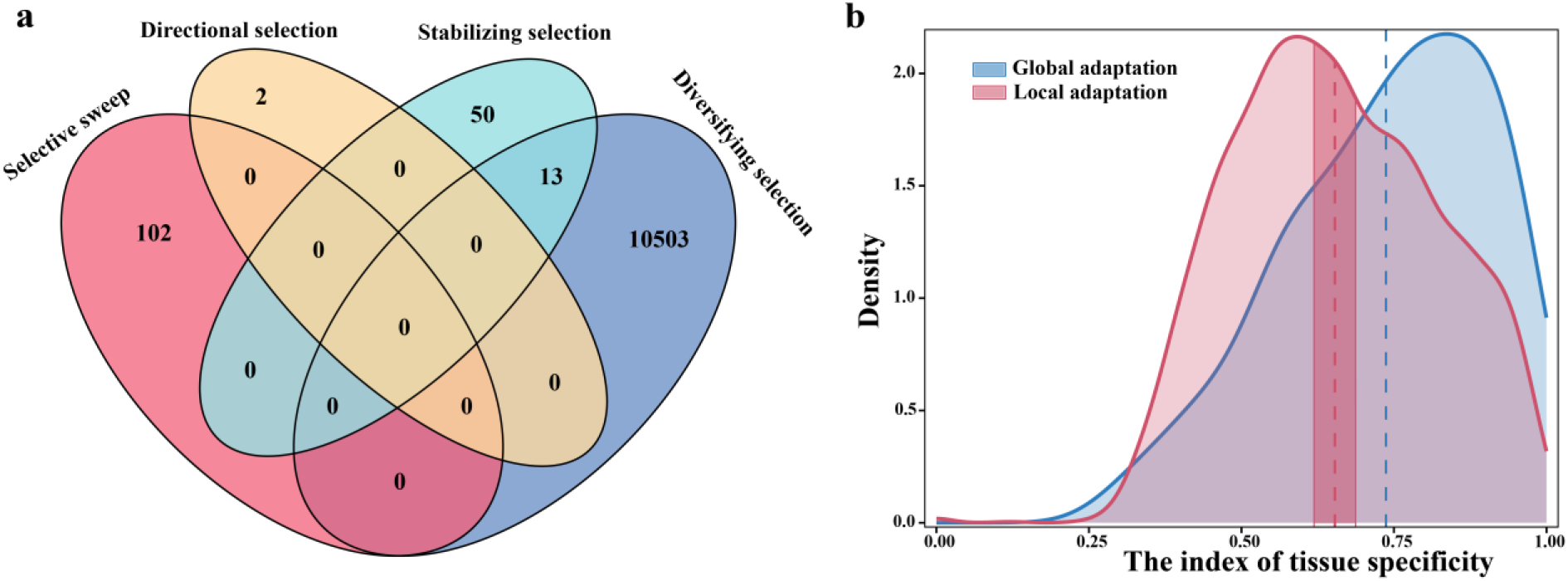
Comparison between global and local adaptation. a) Venn diagram showing the numbers of shared and specific genes identified for four types of natural selection in *G*. *przewalskii*. b) The distributions of tissue-specificity index for global (blue) and local adaptation (red). The dashed line represents the mean tissue-specificity index. The dark red box represents the confidence interval for the mean by bootstrapping with replacement.

2,061 of the genes associated to local adaptation belonged to expanded gene families, which were probably associated to the recent whole genome duplication (WGD) event of *G. przewalskii* (enrichment ratio = 1.38, Fisher’s exact test *p*-value < 2.2e-16). No such correlation could be detected for genes under global adaptation (See **Supplementary Methods S2** for the results of gene expansion and WGD). Furthermore, genes that experienced selective sweeps were significantly enriched in genes that were specific to *G. przewalskii* (enrichment ratio = 1.67, Fisher’s exact test *p*-value =0.007).

### Functional analysis of the genes under selection

Gene ontology enrichment analysis suggested that genes under global adaptation were more likely involved in the response to exogenous stimuli, mainly to water deprivation (8 genes), nutrient starvation (2), hypoxia (3), oxidative (4), temperature/heat (4), light/radiation (8), osmotic stress (5), or to endogenous hormone stimuli (14). 27 genes were related to development and morphogenesis, mainly for root (5) and cell wall (5), especially gene *YODA* for stomatal complex development (see details in **Table S19** and **Supplementary Methods S4)**. For genes under local adaptation, three major biological processes were significantly enriched: 1) response to environmental stimulus (such as response to temperature, light, UV and radiation), DNA repair processes and response to DNA damage stimulus; 2) plant growth, development and reproduction processes (e.g. flowering, development of pollen, zygote, trichoblast, root hair and seed); 3) biosynthesis and metabolism for compounds like pigment and cell wall polysaccharide (see **Table S21**). Genes under selective sweeps were mainly involved in photosynthesis (PSII associated light-harvesting complex II catabolic process, photosystem I assembly, and photoinhibition), metabolites biosynthesis (positive regulation of flavonoid biosynthesis, abscisic acid transport, and coenzyme A biosynthesis), and root development (lateral root branching, and regulation of root meristem growth. See details in **Table S23**). Genes that were both under global and local adaptation were identified that could be essential for the development of morphological traits and response to environmental stress. For example, genes were related to drought or salinity tolerant (*LACCASE14, AT5G05220, UBP12* and *CAMTA1*) ^16-19^, and cell wall development (*BXL2* and *BXL7*) ^20^(**Table S24**).

### Gene expression tissue-specificity

Gene expression analyses based on *Arabidopsis thaliana* Expression Atlas database further revealed that genes of global adaptation had significantly higher tissue-specificity than those of local adaptation (**Fig. 4b**). 93 *A. thaliana* orthologues genes of global adaptation had a mean tissue-specificity index of 0.737, while 6,638 orthologues genes of local adaptation had a mean of 0.653 (a bootstrapping 95% CI = 0.619, 0.687).

## Discussion

To understand how global and local adaptation to aridity have impacted the genome, we first tested for adaptation to aridity across 15 species, eleven of which were growing under arid environments and others not. We identified 118 candidate genes for adaptation to aridity that were shared across xerophytic species. Then, assuming that global and local adaptation are related phenomena, we focused on populations of one xerophytic species, *G. przewalskii*, and tested for local adaptation. Finally, we compared the sets of genes identified at the global and local scale. Below we discuss our findings with respect to global and local adaptation and the relationship between the two processes.

### Global adaptation to extreme environments

As was emphasized in the introduction, our definition of global adaptation will include both convergent and parallel evolution and will give a conservative estimate of the number of global adaptation within *G. przewalskii*. Studies of convergent evolution in plant adaptation to extreme environments generally led to the identification of a limited number of genes ^1^. For example, five and three genes were found harboring amino acid substitutions for repeated adaptations to CAM photosynthesis and carnivore diet, respectively ^21,22^. This is likely because complex traits are usually redundant, i.e. the same phenotype can be produced by many combinations of genes and alleles at those genes ^23^. Furthermore, the repertoire of genes associated to a given trait can also vary across species, either because they are present but not associated to the traits or because they are simply missing. Using genome scans and focusing on series of adaptive traits to environmental stress, 47 to 400 genes were identified in convergent adaptation to polar, alpine, or intertidal zones ^24-26^. In light of these studies, the number of genes, 118, found in the present study to be repeatedly involved in adaptation to arid environment across species that diverged over 100 million-years does not seem too surprising.

In contrast to most studies on the topic, our approach explicitly modeled convergent evolution by simultaneously analyzing the effects of phylogenetic relationship, genetic drift, directional selection and stabilizing selection. This enabled us to highlight the dominant role of stabilizing selection, which did not receive enough attention in studies of convergent evolution. For *G. przewalskii*’s adaptation to aridity, 63 genes were under stabilizing selection and only two were under directional selection. Stabilizing selection is a major evolutionary force maintaining fitness related traits at adaptive optima ^12^. A few substitutions are needed to lead to fitness improvement in extreme environments and cause rapid divergence from other species living under more benign environments. However, in the long term, stabilizing selection at the whole gene level is necessary to eliminate interruptive mutations and constrain the adaptive functions/traits. Therefore, searching for signatures of stabilizing selection across species growing in the same environment or with similar life history traits could be a more fruitful approach for identifying the genetic basis of repeated adaptation.

### Divergence of G. przewalskii populations and local adaptation

In addition to its long-term adaptation to aridity, we showed that *G. przewalskii* was also significantly shaped by local adaptation to the joint effect of geographic landscape and environmental factors such as precipitation and temperature. Orogenic changes occurring in deserts and mountains of northwest China fragmented the species distribution range, limited gene flow between populations and created new environmental regions, driving *G. przewalskii* population divergence ^27,28^. For instance, the Tarim lineage diverged from others during the late Miocene - early Pliocene (∼5.22Mya). This period corresponds to extreme aridification in the entire Tarim Basin due to the retreat of the Paratethys Ocean, the discontinuation of the oceanic water-vapor pathway between the Pamir and the Tian Shan mountain ranges ^29^, and the final and permanent dry-up of episodic lakes ^30^. Similarly, the other lineages diverged during the Pleistocene (2.75 ∼ 0.74 Mya), which was also associated to increased aridity and expansion of deserts (*e.g.* the Taklamakan Desert, the Badain Jaran Desert, Tengger Desert, the Gurbantunggut Desert) in northwest China ^28,31,32^. The quick uplift of the Kunlun Mountains and the Qinghai-Tibetan Plateau during the Pleistocene further blocked the moist air and caused the precipitation at Lop Nur to drop drastically from about 990 mm during the Pliocene to 275 mm during the Holocene, and to 17.4 mm today ^33^. The Turpan-Hami basin is the lowest in altitude and also the hottest place in China, with temperature reaching 48.9 ◦C ^34,35^. Strong winds in Turpan last from April to June and create the largest wind-eroded landforms in the Toksun Yadan National Geopark. The frequent hot and dry winds exacerbate the aridity and soil degradation in this region and force the dwarfism and fragmentation of the vegetation ^36^. Meanwhile the Hexi Corridor is the northwestern boundary of the East Asian summer monsoon where the synchronous variation of precipitation has a significant influence on the local vegetation ^37^. Finally, conditions also varied within each major geographical region. Both precipitation and temperature are heterogeneously distributed in the Alxa region ^38^ and populations of the Tarim lineage are scattered along the Tarim River and its tributaries. In summary, even though the whole distribution range of *G. przewalskii* reflects its adaptation to aridity, local environmental factors, that combine orogeny, the regression of the Tethys, and glaciations, also contributed to further adaptation.

### Global and local adaptation in Gymnocarpos

While global adaptation in the *Gymnocarpos* genus occurred about 35 Mya (∼70 N_e_ generations), i.e., when the ancestor of the genus *Gymnocarpos* diverged in the Southern Arabian Peninsula ^39^, divergence of *G. przewalskii* populations through drift and local adaptation is relatively recent, about 5.2 to 0.7 Mya (1.4 ∼ 10.4 N_e_ generations). Yet, although approximately 20% (10,516 / 51,990) of the protein-coding genes were involved in local adaptation, a mere 13 of those genes were found to be shared across species. This may seem particularly low, but seems in line with previous studies. For instance, while thousands of genes in lodgepole pine and interior spruce showed signature of local adaptation, only 47 locally adapted genes were shared between the two species ^25^. A possible explanation is that strong stabilizing selection on genes under global adaptation keeps removing genetic variation, thereby preventing these genes from responding to more recent environmental changes, especially if selection pressure in extreme environments is not relaxed. Another possible cause is that local adaptation genes rely on expanded gene families, something that was also reported in pine and spruce ^25^. Although our approach can also be easily applied to gene families of any copy number, the present study is conservative by limiting the search to single copy orthologs and therefore should underestimate the overlapping ratio. On the other hand, the results (13 out of 65) mean 20% of the genes under global adaptation for *G. przewalskii* are also under local adaptation.

Functional analyses showed that genes of both global and local adaptation may be represented by multiple family members with diverse functions, i.e. they are highly pleiotropic. For example, Laccase genes exist in bacteria, fungi, plants and animals, and play important roles in defense mechanism, stem tensile strength, and water transport capacity of plants under drought or salinity stress ^40,41^. Signals of purifying selection, selective sweeps, as well as gene duplication, were also detected in Laccase genes in soybean ^42^. Defects of *PPR* gene can cause changes of photosynthesis, leaf development, hypersensitivity to abiotic stress, and restoration of pollen fertility^19^. While the expression of UBP genes modulate drought resistance through stomatal closure ^43^. And UBP genes play an important role in circadian clock and photoperiodic flowering regulation^18^. Mutations in genes regulating stomatal development can also lead to dwarfism or tiny leaves (e.g. *YODA* gene ^44-46^) that may prevent xerophytic plants from losing water and increase resistance to wind damage. This may not come as a surprise considering that plants living in arid environments have to face multiple challenges like extreme heat and coldness, osmotic stress, DNA damage caused by intensive UV-radiation, low precipitation but high transpiration and numerous physiological and morphological changes are involved in adaptation to aridity, such as stomatal regulation, photosynthesis, DNA repair, metabolism and developmental changes. However, genes of global adaptation in general had stronger tissue-specificity in gene expression which means that they may only be crucial to some vital traits or at some key developmental stages, while genes of local adaptation may exhibit rather constant effects on broader tissues.

In conclusion, stabilizing selection plays a dominant role in global adaptation by maintaining individual genes with crucial function for adaptation over millions of years (approximate 70 N_e_ generations in the present case). In contrast, the effect of local adaptation is generally genome-wide and likely occurs through tuning of allele frequencies at a large group of genes and is also driven by gene/whole genome duplication events that took place relatively recently. The limited number of genes involved in both global and local adaptation could indicate that many of the genes associated with the latter could be the result of neofunctionalization or sub-functionalization ^25^.

## Materials and Methods

### Genome sequencing, assembly, and annotation

For reference genome sequencing of *G. przewalskii*, genomic DNA was extracted from young fresh needles of an adult individual (‘Wutuan-1’) growing in a dried riverbed at an elevation of 1315 meters in Wutuan, Xinjiang, China (41.47°N, 80.78°E). Paired-end short reads were generated using Illumina HiSeq 2000 platform. Long reads were generated using PacBio Sequel platform. Scaffolds were *de novo* assembled through SSPACE v3.0^47^ and anchored to 20 pseudochromosomes with the help of a Hi-C chromatin interaction map using LACHESIS ^48^ (**Fig. S2**). The completeness of the genome assembly was evaluated by Benchmarking Universal Single-Copy Orthologs (BUSCO) ^9^ and Core Eukaryotic Gene Mapping Approach (CEGMA) ^10^. We collected flower and needle tissues of *G. przewalskii* for RNA sequencing (RNA-seq), combined with homology-based and *de novo* approaches to predict genes in the *G. przewalskii* genome. Detail methods can be found in **Supplementary Methods S1**.

### Gene family clustering, phylogenetic reconstruction and estimation of divergence times

OrthoFinder v2.4.1 ^49^ was used to identify paralogues and orthologues between *G. przewalskii* and 15 other species. Eleven species were xerophytic: three belong to the Caryophyllales (*Hylocereus undatus*, *Simmondsia chinensis*, and *G. przewalskii*, referred to ‘CX’ hereafter) and eight (*Populus euphratica, Ammopiptanthus nanus, Lycium ruthenicum, Pistacia vera, Pugionium cornutum, Ricinus communis, Phaseolus acutifolius* and *Cicer arietinum*, ‘OX’) belong to other orders. Four non-xerophytic Caryophyllales species, growing under less challenging water and heat conditions, were chosen as control (*Spinacia oleracea*, *Beta vulgaris*, *Dianthus caryophyllus* and *Fagopyrum tataricum*, ‘CNX’, see **Fig. 1a**). *Oryza sativa* as an outgroup species. We then estimated the number of gene families shared between the fifteen species. “Common xerophytic” genes were defined as genes shared by at least four species in class CX and OX but that are absent in any CNX species. For genes with more than one copy, the best orthologs were chosen based on BLASTP (e-value < 1e-5). In total, the “common xerophytic” group included 118 gene families (containing 118 single copy orthologous genes in *G*. *przewalskii* and *S. chinensis*). The amino acid sequences were aligned by MAFFT ^50^ and retro-translated into codon alignment by PAL2NAL ^51^. The detailed methods of phylogenetic reconstruction, divergence time estimation, and gene family expansion and contraction are provided in the **Supplementary Methods S2.**

### The evolution of genes shared by xerophytic species

We counted the number of gene families shared between pairs of 15 species that were classified into xerophytic and non-xerophytic. We first regressed the number of shared gene families on the phylogenetic branch length or both branch length and life history trait contrast (*i.e.* xerophytic versus xerophytic, xerophytic vs non-xerophytic, non-xerophytic vs non-xerophytic) using a generalized linear model with negative binomial distribution. We also applied a linear mixed effect model with the fixed effect of life history trait and the random effect of phylogenetic branch length.

To better evaluate the roles of directional and stabilizing selection, and to discriminate them from genetic drift, we simulated genomic evolution of shared genes between species using an Ornstein-Uhlenbeck process, which can be described by the following stochastic differential equation:

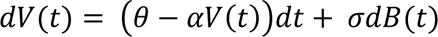

where the directional selection drives the traits to an optimum value, *θ* and the stabilizing selection maintains this optimum value with a strength *α*. The effect of genetic drift was simulated using a Brownian process *dB*(*t*) with the variance equals to σ^2^. Three models were generated: 1) the OUBM model with drift alone (*OU*_(*θ*=0,*α*=0,σ)_), 2) the OUSS model with random drift and the effect of stabilizing selection (*OU*_(*θ*=0,*α*,σ)_), and 3) the OUDS model with the effects of random genetic drift and both directional and stabilizing selection (*OU*_(*θ*,*α*,σ)_). The mean numbers of shared genes and the standard errors were summarized between pairs of species in six classes (“OX – OX”, “CX – CX”, “OX – CX”, “CX – CNX”, “OX – CNX”, and “CNX – CNX”). For each model, parameters (*θ*, *α*, σ) and the total number of genes in the first common ancestral node were drawn from the priors and 2e5 iterations were simulated using function ‘rcOU’ in R package ‘sde’^52^. Approximate Bayesian Computation was used to apply the model selection, goodness-of-fit test, parameter inference, and posterior predictive check using the R package ‘abc’^53^. All R codes are provided in **Supplementary file 2**.

### Estimating k_a_/k_s_ ratios

The *codeml* program with a free-ratio model in PAML ^14^ was used to estimate the *k_a_/k_s_* ratio for the orthologs in each branch of the species tree. To avoid division by zero, we normalized the *k_a_/k_s_* ratio by adding 1 to both *k_a_* and *k_s_*. The distributions of *k_a_/k_s_* ratios for the internal branches (*i.e.* ancestral branches that do not lead to one of the species in the study) and external branches (*i.e.* branches that lead to a given species) were plotted.

### Population genome resequencing and SNP identification in G. przewalskii

Whole genome resequencing was carried out for 177 *G. przewalskii* individuals (**Fig. 2a, Table S14**). The 150 bp paired-end short reads were obtained using Illumina HiSeq 2000 platform with a mean depth of 15×. Adaptor-trimmed paired-end reads from these individuals were aligned to reference genome sequence using BWA-MEM algorithm with default parameters^54^ and sorted by samtools v1.7^55^. To minimize the influence of mapping bias, a few filtering steps were performed before SNP calling: (1) The properly paired reads were kept; (2) sites mapping to different chromosomes were discarded; (3) PCR duplication introduced during library construction was marked using PICARD MarkDuplicates (http://broadinstitute.github.is/picard/). SNP calling was then conducted using BCFtools ^56^, with the command line ‘bcftools mpileup --fasta-ref ref.fasta sorted.bam | bcftools call -mO z -o raw.vcf.gz’. The resulting file included non-variant and variant (SNP and INDEL) sites. To obtain a high-quality dataset, we further employed several filtering steps using BCFtools ^56^ and Vcftools v0.1 ^57^: (1) a SNP was filtered out if SNP quality (QUAL) <30, or depth (DP) < 500 or > 5000, or mapping quality (MQ) < 25, or mean read depth < 8, or mean genotype quality (GQ) < 20, or quality by depth (QD) < 2, or any of the following statistic failed: read position bias (RPB), mapping quality bias (MQB), base quality bias (BQB), mapping quality vs strand bias (MQSB), strand odds ratio (SOR). (2) non-variants sites were filtered if QUAL < 30, or DP < 500 or DP >5 000, or MQ < 20, or MQSB < 0.03, or the mean read depth < 8. (3) SNPs were also discarded if more than two alleles were found, or more than 10% individuals failed, or the minor allele frequency (MAF) < 0.01. In the end, 235,810,604 high-quality sites including 28,250,759 SNPs were retained for the analysis of Stairway Plot2 ^58^. These SNPs were also used to perform an analysis of genome-environment association. For population clustering, admixture analysis, and migration barrier inference, we retained 897,368 SNPs by excluding SNPs in protein-coding regions and SNPs in linkage disequilibrium (LD coefficient *r*^2^ > 0.1) using PLINK v1.9 ^59^.

### Population structure, correlation analysis, migration barrier and demographic inference

To investigate differentiation among *G. przewalskii* populations, a Neighbor-Joining tree was built using MAGE v11.0.10 ^60^. PLINK v1.9 ^59^ and ADMIXTURE v1.3.0 ^61^ were further used for population structure analysis. Tests of “isolation-by-distance” (IBD) and “isolation-by-environment” (IBE) were performed by regressing *F*_ST_/(1-*F*_ST_) on the matrices of pairwise geographical distance and environmental distance using the Mantel test and the partial Mantel test implemented in R package ‘vegan’ ^62^. The spatial features of population structure and barriers to gene flow were visualized using Estimated Effective Migration Surfaces (EEMS) ^63^. Stairway plot2 ^58^ was used to estimate past changes in effective population size and Fastsimcoal2 v2.6.0 ^64^ to infer and compare demographic scenarios. See details in **Supplementary Methods S3.**

### Genome–Environment Association Analysis

In this analysis, we identified genetic signals of local adaptation by scanning population differentiation (X^t^X statistics) and correlating allele frequency to variation in environmental factors (See details in **Supplementary Methods S5**) using BayPass v2.2 ^65^. The X^t^X is analogous to *F*_ST_, but it accounts for the covariance structure of the populations, thus making the analysis robust to complex demographic histories. To reduce bias caused by sample size, we merged the smallest and geographically closest populations JB, SD and LD into one (LSJ), and merged M1 and M2 into one Mongolian population M12. Population M3 contains only two individuals and was excluded from this analysis. First, 897,368 non-genic SNPs free of linkage disequilibrium were used to generate the covariance-correlation matrix (Ω) between populations using the default options for the Markov Chain Monte Carlo (MCMC) algorithm under the core model. The matrix Ω was designed to account for the covariance structure of population allele frequencies caused by shared ancestors and the demographic history (including migration or ancient admixture events)^65^. See **Fig. S18** for a comparison of the covariance matrix and *F*_ST_ matrix. Second, all 28,250,759 SNPs were used to calculate X^t^X statistics under the core model. To identify outlying X^t^X, we performed a calibration procedure using pseudo-observed datasets (PODs) ^65^. For this step, R function simulate.baypass ( ) was used to generate the simulation data with 100,000 SNPs. We then ran BayPass using the same parameters again and defined 0.95 quantile values for X^t^X as a threshold. In the next step, the auxiliary covariate model (-auxmodel) was selected to scan all 28,250,759 high-quality SNPs for signals of local adaptation due to environmental differentiation using default settings but with the burnin increased to 10,000 iterations. The model introduces a Bayesian (binary) auxiliary variable δ to capture some linkage disequilibrium information within a number of pairs of consecutive markers when testing the correlation of allele frequencies to environmental factors. Finally, a Bayes Factor (BF) was derived by posterior mean of each SNP auxiliary variable (known as Posterior Inclusion Probability or PIP) to compare the alternative model (with effect of environmental differentiation) and the null model (environmental effect is not necessary). Twenty-seven representative environmental components of temperature, precipitation, wind speed, solar radiation, and water vapor pressure related together with longitude, latitude, aridity index, and soil salt conditions were standardized (-scalecov) and tested for regression with allele frequency of all SNPs using multivariate gaussian distribution. Candidate SNPs were selected with X^t^X >26 (0.95 quantile values for X^t^X) and BF >10 (the strong evidence suggested by^65^) for each environmental component, respectively. Details on the estimation of linkage disequilibrium (LD) are provided in the **Supplementary Methods S6.**

To evaluate the distribution of candidate SNPs along the genome, the genome was divided into seven regions using SnpEff v4.3 ^66^: 1) the genic regions containing 12.09% SNPs, 2) the proximal regions (within 2 kb flanking 5’ or 3’ end) containing 7.34% SNPs, 3) the distal regions (larger than 2 kb to 5’ or 3’ end) containing 80.57% SNPs. The genic regions were further divided into 4) the intronic regions containing 9.04% SNPs, 5) the UTR regions (0.84%), 6) the synonymous regions containing 0.79% SNPs, and 7) the nonsynonymous regions containing 1.42% SNPs. An empirical enrichment ratio test was carried out by comparing the percentages of candidate SNPs in the above regions to that of total SNPs:

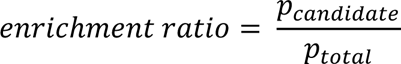

Fisher’s exact test was used to calculate the *p*-value.

### Selective sweep analysis

To identify genomic regions that may have experienced a selective sweep in *G. przewalskii*, we performed whole-genome screening using RAiSD (Raised Accuracy in Sweep Detection)^67^. The software introduced the μ statistic, a composite evaluation test that combines different signatures left by selective sweeps, namely local reduction in polymorphism, shift of the SFS towards both low and high frequency variants and specific pattern of linkage disequilibrium. Phased haplotypes and imputed missing genotypes were obtained using Beagle v5.2 ^68^. 80,347,610 SNPs were used to calculate the μ statistic with sliding windows containing 50 consecutive SNPs each. Nucleotide diversity (π) and Tajima’s *D* were calculated for each sliding window with the software PopGenome ^69^. Values of the μ statistic higher than the 99.999% quantile (1157) of its distribution were considered as candidate regions under selection.

### Gene ontology analysis

Gene Ontology (GO) enrichment analysis was performed for selected candidate genes using *Arabidopsis thaliana* orthologues (blasting with e-value < 1e-5) in PANTHER Classification System on GO Ontology database https://doi.org/10.5281/zenodo.7186998 (released 7 October 2022, http://geneontology.org/). Fisher’s exact test was used to determine the significance of GO terms, and *p*-values were further adjusted by false discovery rate (FDR) to correct for multiple testing.

## Tissue-specificity index

To calculate tissue specificity, we downloaded Expression Atlas database for *Arabidopsis thaliana*^70^ (accession: E-MTAB-7978^71^). We calculated the mean TPM (Transcripts per million) across all developmental stages and sub-tissue types within the tissue type (23 in total) for subsequent analyses. The tissue-specificity index of gene expression (τ) was calculated using the methods presented in ^72,73^.

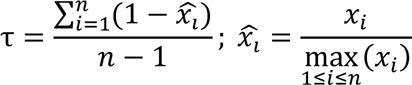

where *n* is the number of tissues and *x*^_*l*_is the mean TPM for a given tissue type normalized by the maximum mean TPM across *n* tissue types. In total, we obtained τ for 30,074 *A. thaliana* genes. For genes of global adaptation and local adaptation, we found 93 and 6,638 *A. thaliana* orthologues (blasting with e-value < 1e-5), respectively. Confidence intervals for the mean tissue-specificity index of the local adaptive gene were calculated based on 10000 bootstrap replicates with 93 samples every time.

## Data availability

All sequencing data used in this study have been deposited in NCBI SRA database with the Bioproject number PRJNA849347 for raw data of reference genome assembly and annotation. And PRJNA885256 for population resequencing libraries. The reference genome and gene annotations have also been deposited in the Genome Warehouse in National Genomics Data Center (https://ngdc.cncb.ac.cn/gwh) under the accession number GWHBKGY00000000. A

## Supporting information

supplemental methods and results

## Acknowledgements

This work was supported by the State Key Basic Research and Development Plan of China (grant no. 2017YFA0605104) granted to Yingxiong Qiu and the National Natural Science Foundation of China (31972946) and the Fundamental Research Funds for the Central Universities 2-2050205-21-688 granted to Jun Chen. The support provided by the China Scholarship Council (CSC) during a visit of Ruirui Fu to Uppsala University is acknowledged (no. 202206320333).

## Author contributions

J.C. designed and managed the project. All analyses were performed by R.F., with contributions to *k_a_/k_s_* calculation from Y.Z. R.F. and Y.L. performed the DNA extraction. Z.Y., R.L., P.L., and Y.Q. contributed plant material collection. R.F., J.C. and M.L. wrote the manuscript.

## Competing interests

The authors declare no competing interests.

## Notes

### Competing Interest Statement

The authors have declared no competing interest.

