## supplemental methods and results for "Global and local adaptation to aridity in a desert plant *Gymnocarpos przewalskii*"

**Supporting Information**

**Methods S1** The genome assembly and annotation of *G. przewalskii*

**Methods S2** Comparative genomic analysis

**Methods S3** Population genomics analysis

**Methods S4** Functional analysis of genes under selection

**Methods S5** Environmental factors

**Methods S6** Linkage disequilibrium

**Fig. S1** *k*-mer analysis for *G. przewalskii* genome size estimation. The x-and y-axes show *k*-mer depth and *k*-mer frequency, respectively.

**Fig. S2** (Left) Circos plot of 20 chromosomes of *G. przewalskii* reference genome, showing the density distributions of (a) repeat element; (b) GC content; (c) SNP number; (d) gene density. The four metrics are calculated in 1 Mb sliding windows. The synteny blocks are shown with lines connecting genes in the center. (Right) Hi-C interaction heat map for *G. przewalskii* reference genome showing interactions between 20 chromosomes.

**Fig. S3** (a) Venn diagram shows that most of predicted gene models are supported by more than one method. (b) Distributions of gene models from *G. przewalskii* and four other species. (c) Venn diagram of functional annotations performed by four databases.

**Fig. S4** Distribution of Synonymous substitution rate (*K_s_*) for paralogous genes within the genomes of *G. przewalskii*, *Hylocereus undatus*, *Simmondsia chinensis*, *Spinacia oleracea*, *Vitis vinifera*, *Beta vulgaris, Dianthus caryophyllus,* and *Fagopyrum tataricum*.

**Fig. S5** The plot of syntenic depths in *G. przewalskii*. (a) *G. przewalskii* vs. *S. oleracea* genome; (b) *G. przewalskii* vs. *V. vinifera* genome.

**Fig. S6** Macrosynteny between genomic regions of *G. przewalskii*, *S. oleracea* and *V. vinifera*. Synteny patterns for 2:1 between *G. przewalskii* and two reference genomes support the independent WGD event of *G. przewalskii*.

**Fig. S7** Gene ontologies (biological processes) for 7,378 duplicated genes for recent WGD associated with expansion gene family. Only those of stress response, growth and developmental processes are shown.

**Fig. S8** The cross-validation errors against *K* values in Admixture analysis.

**Fig. S9** Population structure of *G. przewalskii*. left) Neighbor Joining phylogenetic tree based on intergenic single-nucleotide polymorphisms (SNPs) with LD removed, using *G. decandrus* as outgroup. right) Principal component analysis (PCA) with the first two principal components. Colors correspond to the phylogenetic tree grouping.

**Fig. S10**The genetic diversity (π), neutrality test statistic (Tajima’s *D*) and divergence (*F*_ST_) of four major lineages. Values above the inside of the circle represent nucleotide diversity (π) for the lineage, Values below the inside of the circle represent Tajima’s *D*, and values between lines indicate population divergence (*F*_ST_).

**Fig. S11** The result of Stairway Plot2 on all lineages for *G. przewalskii.* The thick lines represent the median values of population sizes and the thin lines show the 5% and 95% confident intervals.

**Fig. S12** Schematic diagram of ten priori demographic models with different migration events for *G. przewalskii* by fastsimcoal2.

**Fig. S13** Likelihood ratio G-statistics distribution. The likelihood ratio G-statistics (*CLR* = log10(*CL*_O_/*CL*_E)_, where *CL_O_* and *CL_E_* are the observed and estimated maximum composite likelihood, respectively) was checked how data fit the chosen model. A non-significant *p*-value of this test suggests that the observed SFS is well compatible with the model. The red line is the observed CLR.

**Fig. S14** The distribution of *ka* (left), *ks* (right) for “common xerophytic” genes.

**Fig. S15** The distributions of the number of SNPs under diversifying selection for 20 chromosomes. Each dot shows the SNP number for 100 kb windows sliding in increments of 10 kb.

**Fig. S16** The distributions of the number of SNPs and genes under diversifying selection for different environmental factors. (Left) Distribution of number of SNPs associated to environmental factors in protein-coding genes. (Right) Distribution of the number of genes associated to environmental factors. Vertical black bars indicate SNPs or genes that are associated to various environmental factors. Horizontal colored bars correspond to SNPs or genes associated to one environmental factor. salt, salinity; temp, temperature; prec, precipitation; wind, wind speed; lon, longitude; A.I., aridity index; srad, solar radiation; lat, latitude; vapr, water vapor pressure.

**Fig. S17** Enrichment analysis of different classes of SNPs under diversifying selection for nine environmental factors. Enrichments shown are relative to the proportion of each class of SNPs in the genome overall. The horizontal dashed line shows the expected enrichment under the null hypothesis of no enrichment. The colored dots show the enrichment values for each class of SNPs. Gray dots represent 10,000 null permutations of site categories. Statistical significance was determined by Fisher’s exact test (*p*-values: * < 0.05, ** < 0.01, *** < 0.001, **** < 0.0001. genic, SNPs in genic regions; intron, intronic SNPs; inter(>2k), SNPs in distal regions (larger than 2 kb to 5’ or 3’ end); inter(<2k), SNPs in proximal regions (within 2 kb flanking 5’ or 3’ end); nonsys, nonsynonymous SNPs (amino acid changing SNPs); synon, synonymous SNPs; TE, Transposable element.

**Fig. S18** The covariance matrix of *G. przewalskii* populations (left) and *F*_ST_ matrix of *G. przewalskii* populations (right).

**Table S1** *G. przewalskii* reference genome sequencing library statistics.

**Table S2** Estimation of the *G. przewalskii* genome size based on 17-mer statistics.

**Table S3** Summary of contigs and scaffolds for *G. przewalskii* reference genome.

**Table S4** Genome assembly completeness evaluation by BUSCO.

**Table S5** Genome assembly completeness evaluation by CEGMA.

**Table S6** The statistical results of reads coverage for the *G. przewalskii* genome.

**Table S7** The statistical results of repeat sequence for the *G. przewalskii* genome.

**Table S8** Classification of transposable elements (TEs) identified in the *G. przewalskii* genome,
including their length (in bp) and the percent of genome.

**Table S9** Summary of protein-coding genes identification by three sources.

**Table S10** The statistical results of gene structure of closely related species.

**Table S11** Summary of function annotation for gene.

**Table S12** Summary statistics of non-coding RNAs in the *G. przewalskii* genome.

**Table S13** The significant GO term of duplicated genes from recent WGD associated with expansion gene family in *G. przewalskii*. (see separate file)

**Table S14** Sample information of *G. przewalskii* for resequencing. (see separate file)

**Table S15** The matrix of *F*_ST_ for all populations in the study. (see separate file)

**Table S16** Details of single environment factors from WorldClim database used for the populations of this study. (see separate file)

**Table S17** Comparison likelihood of 10 models performed in fastsimcoal2.

**Table S18** Estimation of the demographic parameters with 95% highest posterior density (HPD) for the best model. (model 7)

**Table S19** The GO term of “common xerophytic” genes under stabilizing selection. (see separate file)

**Table S20** The GO term of “common xerophytic” genes under directional selection. (see separate file)

**Table S21** The significant GO term of local adaptation in *G. przewalskii* from Baypass. (see separate file)

**Table S22** The gene function of local adaptation in Chr.5: 89.89 Mb-90.46 Mb for *G. przewalskii*. (see separate file)

**Table S23** The significant GO term of candidate genes under selective sweep. (see separate file)

**Table S24** The function of genes under convergent evolution and local adaptation. (see separate file)

**Methods S1 The genome assembly and annotation of** ***G. przewalskii***

***1.1 Genome sequencing, size estimation and assembly***

An adult *G. przewalskii* (‘Wutuan-1’) used for sequencing the species reference genome was from Wutuan, Xinjiang, China (41.47°N, 80.78°E), growing in dried riverbeds at an elevation of 1315 meters. Genomic DNA was extracted from young fresh needles with Plant Genomic DNA Kit (TOLOBIO, Shanghai, China) following the manufacturer’s instructions from fresh leaves which were harvested and frozen immediately in liquid nitrogen. Selected high-quality DNA samples were sent to Novogene for both Illumina and PacBio sequencing. Paired-end genomic sequence library was constructed with an insert size of 350 bp. Sequencing was implemented on the Illumina HiSeq 2000 platform (Illumina, NEB, USA) and generated 150-bp paired-end reads. The PacBio single-molecule real-time (SMRT) sequencing library of 20-kb insert size was generated with a SMRTbell Template Prep Kit (PacBio) and sequenced on a PacBio sequel platform (Pacific Biosciences, CA, USA) with 12 cells. In total 593 Gb raw reads were obtained with read depth ~ 256× (**Table S1**). All sequencing reads were trimmed to remove adaptors and enhance quality.

The genome size of *G.przewalskii* was estimated using *k*-mer method^1^ (Genome Size = *k*-mer_num/Peak_depth). In order to calculate and plot the *k*-mer frequency distribution, we first generated the 17-mer distribution of sequencing reads from 208,139.9 Mb Illumina short reads and then calculated the genome size (G) based on the above formula. The number of *k*-mer was 130,281.3 Mb and the main peak of *k*-mer depth was equal to 56. Therefore, the genome size was estimated to be 2,326.45 Mb and was revised to 2,310.3 Mb by filtering out false *k*-mers (**Fig. S1, Table S2**).

We performed genome assembly by combining long PacBio reads, Illumina short reads, and Hi-C sequencing reads. The PacBio reads, corrected by CANU v1.8^2^, were first applied to generate the primary contigs using FALCON v0.3.0^3^ (<https://github.com/PacificBiosciences/FALCON/>). Subsequently, contigs were polished by mapping all filtered PacBio reads with QUIVER^4^ and were further corrected using Illumina short paired-end reads by PILON v1.22^5^. These improved contigs were assembled into scaffolds through SSPACE v3.0^6^. Finally, the clean reads of Hi-C library were aligned to scaffolds using BWA v0.7.12^7^ and the uniquely mapped reads were retained for subsequent assembly. The scaffolds of 1,978 Mb (94.44% of the whole-genome) were divided into subgroups, sorted and oriented into 20 pseudochromosomes of *G. przewalskii* using LACHESIS^8^ (**Fig. S2**). The final assembled genome size was 2,095 Mb with scaffold N50 of 91.85 Mb (**Table S3**).

The completeness of genome assembly was evaluated by Benchmarking Universal Single-Copy Orthologs (BUSCO)^9^ and Core Eukaryotic Gene Mapping Approach (CEGMA)^10^. 88.4% of 1440 single-copy orthologs were retained and 8.8 % were missing by BUSCO (**Table S4**). On the other hand, 236 of 248 (95.16%) Core Eukaryotic genes were retrieved by CEGMA (**Table S5**). To further evaluate the consistency of the assembly, all the paired-end reads were mapped to the reference using BWA v0.7.12^7^. The mapping rate of paired-end reads was 98.2% and the coverage rate was 97.7 % (**Table S6**). Taken together, these results showed that the genome of *G. przewalskii* was well assembled, with high-quality, completeness, and accuracy.

***1.2 Genome annotation***

Tandem repeat sequences were identified by Tandem Repeats Finder v4.09(TRF)^11^. Transposable elements (TEs) were identified with a combination of homology-based and *de novo* approaches. Firstly, we used RepeatMasker v.3.3.0^12^ to detect interspersed repeats and low complexity DNA sequences. And RepeatProteinMasker^13^ was used to find repeat sequences at the protein level with the TE protein database. Secondly, *de novo* identification was implemented by RepeatModeler v1.0.8^14^, LTR_FINDERv1.0.5^15^ and RepeatScout^16^ and then annotation was carried out using RepeatMasker v3.3.0^12^. Finally, all repeat identification results from the different software packages were integrated and redundancy was eliminated to produce the final repeat annotation. For summary, 61.58% of the genome is composed of repetitive sequences (**Table S7**), of which Class I (retrotransposons) and Class II (DNA transposons) comprised 59.18% and 0.13% of the genome, respectively. For retrotransposons, particularly long terminal repeats (LTRs) account for 59.06% of the genome (**Table S8**).

Protein-coding genes were identified by combining three complementary methods: homology-based, *de novo* prediction and transcriptome-based predictions. For homology-based predictions, protein sequences of five species including *Fagopyrum tataricum*, *Beta vulgaris*, *Chenopodium quinoa*, *Dianthus caryophyllus*, *Fagopyrum esculentum* were downloaded and aligned to *G. przewalskii* genome by TBALSTN (e-value ≤ 1e-5) as implemented in BLAST^17^. BLAST hits were concatenated by SOLAR v.0.0.19^18^ and GENEWISE v.2.2.0^19^ was used to predict gene structures and define gene models contained in each protein region. For *de novo* prediction, five software including AUGUSTUS v.2.5.5^20^, GENSCAN v.1.0^21^, GLIMMERHMM v.3.0.1^22^, GENEID v.1.4.4^23^ and SNAP^24^ were utilized to predict protein-coding regions in the repeat-masked genome. For transcriptome-based predictions, flower and leaf tissue samples were collected for Illumina RNA-seq sequencing. Short reads were aligned to the reference genome using tophat v.2.0.11^25^ and genome-based transcript assembly was conducted using cufflinks v.2.2.1^26^. Next, the assembled transcripts were aligned to the reference genome and filtered by Program to Assemble Spliced Alignment v2.3.3 (PASA) ^27^ with complete open reading frames. All predictions of gene models yielded by the above three sources were integrated and redundancy was eliminated to generate a consensus gene set using evidencemodeler v.1.1.1^28^. Finally, the non-redundant set was updated by PASA v2.3.3 to generate the information about untranslated regions (UTRs) and alternative splicing and to obtain final gene models (**Table S9** and **Fig. S3a**). In total, we obtained 51,990 protein-coding genes with an average length of 1.04 kb and an average of 4.23 exons per gene. And 49,641 (95.48%) of these genes were distributed on the assembled 20 chromosomes of *G. przewalskii*. The comparison to five other genomes of sequenced species has shown that the genome of *G. przewalskii* is among the species of the best quality (**Table S10** and **Fig. S3b**).

Functional annotation of protein-coding genes was achieved using BLASTP (e-value ≤ 1e-5) against public databases including NCBI non-redundant protein database (NR), SwissProt and Kyoto Encyclopedia of Genes Genomes (KEGG). Only genes with the best match and highest score were retained. Protein domains were annotated by searching InterPro and Pfam using InterProScan v4.8^29^ and HMMER v3.1^30^, respectively. GO annotations were accomplished using the Blast2GO pipeline v3.1.3^31^. In summary, 93.60% (48,670) of genes in the *G. przewalskii* genome were annotated (**Table S11, Fig. S3c**).

Transfer RNA (tRNA) annotation was carried out using tRNAscan-SE v1.3.1^32^. For the prediction of conserved ribosomal RNA (rRNAs), we searched rRNA sequences by aligning against sequences of closely related species (*Fagopyrum tataricum*, *Beta vulgaris*, *Chenopodium quinoa*, *Dianthus caryophyllus*, *Fagopyrum esculentum*) rRNA sequence using BLAST. Other ncRNAs, including microRNA (miRNA) and small nuclear RNA (snRNA), were identified by INFERNAL v1.1.1^33^ against the Rfam^34^ database. In the aggregate, we detected 794 microRNA (miRNA), 1,435 transfer RNA (tRNA), 380 ribosomal RNA (rRNA), and 728 small nuclear (snRNA) in the genome sequence (**Table S12**).

**Methods S2 Comparative genomic analysis**

***2.1 Genome synteny and whole-genome duplication***

Orthologous and paralogous genes were identified by protein sequence similarity searching within and between pairs of genomes of *G. przewalskii*, *Hylocereus undatus*, *Simmondsia chinensis*, *Spinacia oleracea*, *Vitis vinifera*, *Beta vulgaris, Dianthus caryophyllus, Fagopyrum tataricum* using BLASTP (e-value < 1e-5)^17^. Collinear blocks between pairs of genomes were identified by the program WGDI^35^ with the option ‘-icl’. Synonymous substitution rates (*K*_s_) for orthologous genes were calculated using the PAML^36^ package under the Nei-Gojobori model and the median value for each collinear block was used to obtain the distribution of *K*_s_. The whole genome duplication events (WGD) were identified using the peaks of *K*_s_ distribution by the program WGDI with the option ‘-pf’. The package JCVI (<https://github.com/tanghaibao/jcvi>) was used to visualize the genomic collinearity.

To estimate the time of WGD, we first estimated the evolutionary rate of *G.* *przewalskii*. A total number of 3,824 single-copy orthologous genes for *G. przewalskii* and three closely related species in Caryophyllales (*Spinacia oleracea*, *Beta vulgaris*, *Dianthus caryophyllus*) were identified by OrthoFinder v2.4.1^37^. The fourfold degenerate sites were extracted by an in-house Perl script and the overall substitutions per site per year (*r*) were calculated based on the concatenated sequences and the divergence time between *D. caryophyllus* and *S. oleracea* (median: 54 Myr) using ‘baseml’ in PAML^36^. The divergence time to WGD was then estimated from the formula T = *K*_s_ / 2*r*, where *r* equaled 8.37 × 10^-9^ per site per year and *K*_s_ was derived from the peak value of the above *K*_s_ distribution.

***2.2 Gene Family Clusters***

The protein-coding sequences of *G. przewalskii* and 15 other species were downloaded from Phytozome v12 (https://phytozome.jgi.doe.gov/pz/portal.html), the NCBI website (<https://www.ncbi.nlm.nih.gov/>) and the genome warehouse in Beijing Institute of Genomics (BIG) data center (<http://bigd.big.ac.cn/gwh>) for gene family analysis. With *Oryza sativa* as an outgroup species, the other 15 species could be classified into three groups based on habitat and phylogenetic relationships, including 1) four non-xerophytic species in Caryophyllales order (*Spinacia oleracea*, *Beta vulgaris*, *Dianthus caryophyllus, Fagopyrum tataricum,*‘CNX’) growing in a broad range of water supply, and 11 xerophytic species that grow in semi-arid/arid environment with 2) three species in the Caryophyllales order (*Hylocereus undatus*, *Simmondsia chinensis*, and *G. przewalskii*, ‘CX’ hereafter) and 3) eight outside of the Caryophyllales (*Populus euphratica, Ammopiptanthus nanus, Lycium ruthenicum, Pistacia vera, Pugionium cornutum, Ricinus communis, Phaseolus acutifolius, Cicer arietinum*, ‘OX’). To perform gene family clustering analysis, only the longest transcript isoforms were retained for each gene. Next, genes that encode fewer than 50 amino acids were removed. Then an all-versus-all comparison of protein sequences of 16 species was performed using BLASTP ^17^ with an e-value cutoff lower than 1e-5. Orthologous gene families were identified and grouped by OrthoFinder v2.4.1^37^ with default parameter settings. Gene families shared by at least two species were retained for further analysis.

***2.3 Phylogenetic Analysis and Divergence Time Estimate***

In total, 122 single-copy orthologous gene families were identified from the above gene family clustering analysis and used to reconstruct the phylogenetic relationship. Protein-coding DNA sequence (CDS) alignments of each single-copy family were obtained by the protein sequence-guided alignment using MAFFT v7.471^38^ and PAL2NAL^39^. The maximum likelihood phylogenetic tree was constructed based on a concatenated CDS matrix using RAxML v8.2.12^40^ with the “GTR+G+I” nucleotide substitution model. A bootstrap of 1000 iterations was applied to calculate the supporting score for each phylogenetic branch. The divergence time of ancestral nodes was inferred using the ‘mcmctree’ package in PAML^36^ with the ‘relaxed-clock (clock = 2)’ model and ‘HKY85’ model. The fossil-based divergence time between *O. sativa* - *B. vulgaris* (115 - 308 Mya), *B.vulgaris* - *F. tataricum* (73 - 91 Mya), *B. vulgaris* - *P. euphratica* (111 - 131 Mya), *P. euphratica* - *C. arietinum* (101 - 131 Mya), *P. euphratica* - *R. communis* (70 - 86 Mya) were obtained from the TimeTree database (<http://www.timetree.org/>) and used to calibrate the tree.

***2.4 Gene family expansion and contraction***

According to the results of gene family clustering analysis, only gene families shared by at least two species were preserved, and gene families were filtered out if more than 100 copies were found in any single species. The 19,193 gene families were generated for gene family expansion and contraction analysis. CAFE v.4.2.1 ^41^ was used to infer the gene families’ expansion and contraction along all phylogenetic branches for all 16 genomes with a significance threshold of *p*-value $\leq$0.05.

***2.5 The influence of gene family expansion and whole genome duplications***

For 19,193 gene families shared by two or more species, we explored the number of families that experienced expansion and contraction for each species. A mean of 3081, 3586, and 2499 gene families were under expansion, which corresponded to 23.5%, 27.1% and 19.9% of total examined gene families for OX, CX and CNX groups, respectively.

To search for the causes of gene expansion, we further tested the correlation between expanded gene families and the ones experienced whole genome duplication. With an estimate of synonymous substitutions rate of 8.37 × 10^-9^ per base per year, the distribution of *K*_s_ had revealed two WGD events of *G. przewalskii* genome (**Fig. S4**), one around 121.6 Mya corresponding to the γ triplication event of core eudicots, and the other around 19.4 Mya, a recent duplication event specific to *G. przewalskii*. To prove this, we performed syntenic analysis by comparing *G. przewalskii* genome to *S. oleracea* ^42^ and *V. vinifera* ^43^, both of which are paleohexaploid without any recent WGD events. The results revealed that up to 11% and 21% of *V. vinifera* and *S. oleracea* genomes, respectively, had two ortholog copies within *G. przewalskii* genome, and barely had higher ortholog copy numbers (**Fig. S5**). While for *G. przewalskii* genome only 1:1 syntenic genes could be identified in the other two genomes. Similarly, the microsynteny profile revealed a 2:1 synteny share between *G. przewalskii* and the other two genomes (**Fig. S6**). Particularly, one between Chr.12 (27.3%), Chr.15 (66.7%) of *G. przewalskii* and Chr.18 of *V. vinifera* and another between Chr.14 (21.7%), Chr.15 (30.4%) of *G. przewalskii* and Chr.5 of *S. oleracea*. All these confirmed that *G. przewalskii* had experienced a specific recent WGD event. Besides, we also identified recent WGD events specific to *H. undatus* and *S. chinensis*, respectively (**Fig. S4**). While for group CNX no WGD events were identified except *F. tataricum*. For group OX, previous reports have shown that they share WGD events at family and genus levels except for *R. communis* and *P. vera*^44-51^*.*

For 5,845 gene families under expansion in *G. przewalskii*, 3,245 (7,717 genes) were identified as the result of two WGD events (*K*_s_, 0.05-0.58 and 1.71-2.36, respectively), of which 236 families including 339 genes in the γ event and 3,150 families with 7,378 genes in the recent WGD event. For 1,464 gene families under expansion in *S. chinensis*, 395 (842 genes) were identified as the result of two WGD events (*K*_s_, 0.63-1.19 and 1.6-2.38, respectively), of which 130 families including 197 genes in the γ event and 323 families with 645 genes in the recent WGD event. For 3,448 gene families under expansion in *H. undatus*, 1,456 (2,998 genes) were identified as the result of two WGD events (*K*_s_, 0.61-0.98 and 1.6-2.37, respectively), of which 336 families including 541 genes in the γ event and 1,278 families with 2,457 genes in the recent WGD event. These results suggested that the recent WGD event should be the main cause of gene expansion in *G. przewalskii* (enrichment ratio = 1.40, Fisher’s exact test *p*-value < 2.2e-16) and two other xerophytes of Caryophyllales (enrichment ratio = 2.03, Fisher’s exact test *p*-value < 2.2e-16 in *S. chinensis*, and enrichment ratio = 2.08, Fisher’s exact test *p*-value < 2.2e-16 in *H. undatus*, respectively).

The analysis of gene ontology enrichment showed 7,378 duplicated genes from the recent WGD event associated with the expansion gene family in *G. przewalskii* were enriched in multiple GO categories. Some genes showed significant functional representation of organ developments and stress responses, including anther development, flower development, root hair elongation, leaf morphogenesis, seed maturation, response to cold, response to salt stress, and response to water deprivation. And other genes associated with signaling pathways (such as red or far-red light and plant hormones mediated signaling pathways) and cell wall biogenesis (e.g. cellulose biosynthetic process) (**Fig. S7,** see a complete list in **Table S13**).

**Methods S3 Population genomics analysis**

***3.1 Plant material and population genome resequencing***

The genus *Gymnocarpos* is a group of xerophytic plants with about ten species living in the arid regions of Asia and Africa^52,53^. The aridification since the mid-late Miocene had accelerated the diversification of the genus ^52^. *Gymnocarpos przewalskii* Bunge ex Maximowicz (2n=2×=40^54^) is a unique species in the arid regions of northwest China (including Xinjiang, Qinghai, Gansu, Ningxia, and Inner Mongolia) and Mongolia. It is a deciduous perennial shrublets, usually growing along ancient watercourses at an elevation of 800-2500 meters (<http://www.efloras.org/>).

In this study, 177 individuals were sampled from 26 natural populations of *G. przewalskii* covering its distribution range from the westmost corn of Xingjiang (74.67ºE, 39.32ºN) to the mid-Inner Mongolia in the east (106.81ºE, 39.26ºN), and from Ningxia in the south (105.02ºE, 37.44ºN) to West Mongolia in the north (91.56ºE, 46.42ºN). In addition, we obtained one individual of *G. decandrus* from Israel (35.39ºE, 31.58ºN**, Table S14**) to serve as an outgroup for phylogenetic analysis.

A total of 177 *G. przewalskii* and one *G. decandrus* individuals were collected for genome re-sequenced. Total DNA was extracted with Plant Genomic DNA Kit (TOLOBIO, Shanghai, China) following the manufacturer’s instructions. About 10 µg genomic DNA from each sample was used to construct a sequencing library to obtain paired-end short reads with a length of 150 bp on an Illumina HiSeq 2000 platform (Illumina, NEB, USA) with a mean depth of 15×.

***3.2 Population structure***

A Neighbor-Joining tree was built based on the genotype matrix of 897,368 SNPs for 177 *G.* *przewalskii* individuals using MAGE v11.0.10^55^, with the outgroup *G. decandrus* to root the tree. A Principal Component analysis (PCA) was also carried out using PLINK v1.9^56^, and the first two eigenvectors of the largest contributions to the variance were plotted. Ancestral component inference and population admixture analysis were performed using ADMIXTURE v1.3.0^57^ on all *G. przewalskii* samples. ADMIXTURE analysis was run for *K* (the number of ancestral components) from one to nine with a bootstrap of 1000 iterations. The optimal *K* value was chosen based on fivefold cross-validation (**Fig. S8**). Genetic divergence (*F*_ST_) and genetic diversity (pairwise nucleotide difference π) for each lineage were estimated using Vcftools^58^ with a sliding window size of 100 kb and a step size of 10 kb. Tajima’s *D* was also calculated for 100 kb nonoverlapping windows along the genome.

The Neighbor-Joining phylogenetic tree supported all individuals could be grouped into four clusters: 1) the Tarim cluster that includes the westernmost populations; 2) the Hami cluster of the northernmost populations together with two Mongolian populations (M1 and M2); 3) the Hexi cluster of central populations; and 4) the Alxa cluster of the easternmost populations and one Mongolian population M3 (**Fig. S9**). The same four lineages were also recovered by PCA and Admixture analyses (with optimal *K* = 4, **Fig. S9**). The latter also exhibited admixed ancestry components in population JT and MQ where Hexi and Alxa lineages are in contact, and in population LY where Hami, Hexi and Alxa lineages met. Genetic divergence as measured by *F*_ST_ was limited between the four lineages, varying from 0.05 between Tarim and Hexi to 0.083 between Tarim and Hami (**Fig. S10**), as well as between populations (*F*_ST_ from 0.012 to 0.198, **Table S15**). The Tarim lineage harbored the lowest level of genetic diversity ($\pi=4.85\times{10}^{-4}$), Alxa and Hexi lineages had intermediate values ($7.43\times{10}^{-4}$ and 7.8$4\times{10}^{-4}$) and Hami the highest (9.12$\times{10}^{-4}$, **Fig. S10**).

***3.3 Migration barrier inference***

To visualize spatial features of population structure and identify barriers to gene flow, we used Estimated Effective Migration Surfaces (EEMS)^59^. The method uses locality information of geo-referenced genetic samples and pairwise genetic dissimilarity matrix derived from genome-wide nucleotide variations to highlight geographic regions where genetic similarity decays faster or slower than expected under an “Isolation-by-Distance” model. A deme size was used to determine the geographic grid size and the set of migration routes. The observed pairwise genetic dissimilarity matrix were produced using the program ‘bed2diffs_v2’ implemented in the EEMS package. The expected genetic dissimilarity between pairs of populations was calculated by integrating overall possible migration routes. The estimation procedure adjusted the migration rates for all edges in the graph so that the genetic differences expected under a stepping-stone model matched the observed genetic differences. To reduce the potential influence of grid size, a different number of demes (50, 100, 200, 300, 400, and 500) was chosen, and for each size five runs were performed using ‘runeems_snps’ with 15,000,000 MCMC iterations and 6,000,000 burnins. A combined estimate averaged across all runs of different numbers of demes was obtained using the package ‘rEEMSplots’ in R. Finally, EEMS estimates were interpolated across habitat space to visualize potential corridors and barriers of gene flow.

The result confirmed the isolation of Tarim from the other three lineages by revealing a migration barrier separating Tarim from the others. The migration barrier is centered around the Lop Nur, with the Kumtag Desert in the southeast, the Taklamakan Desert in the southwest, and the Tianshan Mountains and the Turpan Basin in the north. Close to the central region, the plant diversity is extremely low due to the extreme aridity and salinity, extreme heat in summer and extreme cold in winter, and abrasive sandstorm ^60,61^. Two regions of high effective migration rates were also found, one around Aral City in the west where four rivers (the Hotan River, the Yarkant River, the Kashgar River, and the Aksu River) flow into the Tarim River, the longest inland river in China. The other is called Hexi Corridor with the Heihe River (the second longest inland river in China) and the Shule River (the second longest inland river in the Hexi Corridor), cutting through the migration barrier in the east.

***3.4 Correlations of geographical distance, environmental factors and genetic distance***

To detect the effects of geographic distance and environmental differences on genetic differentiation between *G.* *przewalskii* populations, tests of “isolation-by-distance” (IBD) and “isolation-by-environment” (IBE) were performed by regressing *F*_ST_/(1-*F*_ST_) on the matrices of pairwise geographical distance and environmental distance using the Mantel test implemented in R package ‘vegan’^62^. The matrix of geographical distance was generated by the R package ‘fossil’^63^. The environmental distance matrices were constructed for the precipitation, temperature, solar radiation, wind speed, and water vapor pressure related environmental factors as well as an overall matrix *E* based on all 104 environmental factors (including salinity, see details in **Table S16**) using the R package ‘cluster’^64^. Matrices of aridity index, longitude, and latitude were also tested for IBE. To avoid correlation between matrices, the independent contribution of each matrix was evaluated by controlling for the effects of all others using the partial Mantel test.

The correlation between genetic divergence *F*_ST_/(1-*F*_ST_) and geographic distance or environmental distance was significantly positive when regressed separately (coef. = 0.47, *p*-value =0.001, and coef. = 0.41, *p*-value = 0.003, respectively). Partial Mantel tests were also applied to control the covariance of geographic distance and environmental factors. The coefficients were reduced but remained significant (coef. = 0.37, *p*-value =0.001 for the geographic distance and coef. = 0.27, *p*-value = 0.044 for the environmental distance, respectively), suggesting population divergence of *G. przewalskii* should be the combined result of genetic drift and local adaptation to environmental factors. We also performed Mantel and partial Mantel tests separately within Tarim and within other clusters to evaluate the contribution of the migration barrier. In this case, the environmental distance did not show any significant effect across populations of the Tarim region while the effect of geographic distance became stronger on both sides of the migration barrier (coef. = 0.75, *p*-value = 0.006 for Tarim populations, and coef. = 0.6, *p*-value = 0.001 for others, respectively). For individual environmental factors, longitude, latitude and precipitation contributed significantly to divergence among Tarim populations while temperature, solar radiation, and wind speed were also found significant in populations to the north and the east of the barrier.

***3.5 Nonparametric demographic inference by*** ***Stairway Plot2***

Based on the results of the population structure analyses, we used a nonparametric approach to infer the demographic history of *G. przewalskii* based on the site frequency spectrum (SFS) using Stairway Plot2^65^ with 500 bootstraps (ninput 500). The folded SFS of four clusters was [generate](javascript:;)d using SoFoS (<https://github.com/CartwrightLab/SoFoS>) with genetic polymorphism data scaled to 110 ,64, 110 and 70 from Tarim lineage to Alxa lineage, respectively. The result showed all lineages of *G.przewalskii* experienced a continuous decline in population size (Ne) from the Last Glacial Maximum to very recent (Ne ≈ 4,000) (**Fig. S11**).

***3.6 Demographic inference based on 2D-SFS (Fastsimcoal2)***

To fully understand the divergence history and population dynamics of *G. przewalskii,* we built and compared ten demographic scenarios based on the results of clustering analyses and stairway plot2^65^, using Fastsimcoal2 v2.6.0^66^ (**Fig. S12**). The origin of admixed populations (JT and MQ) was tested for admixture between Hexi and Alxa lineages (model 2, 4, 6, 8, and10) or derived from Alxa lineage (model 1, 3, 5, 7, and 9). Migrations were also tested for different ancestral and descendant populations (no migration: model 1&2; ancestral migration: model 3&4; descendant migration: model 5&6; no migration between Tarim: model 7&8; full migration: model 9&10).

The minor allele spectra (a.k.a. site frequency spectrum, SFS) of all four lineages as well as the admixed populations (JT & MQ) were derived using easySFS (<https://github.com/isaacovercast/easySFS>) with the number of alleles projected to eight. For each model, coalescent simulations were run for 100,000 iterations and 40 ECM cycles were used to maximize the expected model likelihood. The best model was selected based on the highest Akaike’s weight value^66^. For the best-supported model, 100 replicates were run to gain the best point estimates of all parameters. To obtain 95% confidence intervals of parameters, a parametric posterior bootstrap was performed for 100 replicates. The goodness-of-fit for the best model was evaluated using the likelihood ratio *G*-statistic (*CLR* = log10(*CL_0_*/*CL_E_*))^66^. *CL_0_* and *CL_E_* are the observed and estimated maximum composite likelihood, respectively. A non-significant *p*-value of this test indicated that the observed SFS is well explained by the model (**Fig. S13**).

We found that ancestral migration between Tarim and the three other lineages (m_Tarim_others_ = 2.12e-4 and m_others_Tarim_ = 5.79e-6) right after the split of the Tarim lineage about 5.22 Mya (95% HPD: 5.28 ~ 2.90 Mya) and ceased after the Hami lineage diverged about 2.7 Mya (95% HPD: 1.2 ~ 2.7 Mya) was the best-supported model based on AIC’s weight (model 7 in **Table S17**). Migration persisted between Hami, Hexi, and Alxa lineages, at a slightly higher rate from west to east (2.61e-5 ~ 1.07e-4) than in the opposite direction (3.86e-6 ~ 5.01e-5). Hexi and Alxa lineages diverged about 1.42 Mya (95% HPD: 1.44 ~ 0.13 Mya). Population size contractions were identified in all lineages around 56 Kya (78,570 ~ 545 years ago). The best-supported model also indicated that the signature of admixture between JT and MQ is the result of ancestral polymorphisms instead of secondary contact (**Table S17** for results of model comparison and **Table S18** for parameter estimates).

**Methods S4 Functional analysis of genes under selection**

For genes under stabilizing selection in “common xerophytic” genes, though not significant enriched signals were still identified (fold ratio >1) including 40 (42.1%, 40/95) genes in 39 hierarchical GO categories involved in response to exogenous stimuli (e.g., response to water deprivation, oxidation, temperature/heat, and light/radiation) (**Table S19**). 27 genes were enriched in 20 GO categories related to development and morphogenesis, mainly for root (5) and cell wall (5), especially the gene *YODA* for stomatal complex development. *YODA* regulates stomatal density and spacing patterns in response to variable light conditions ^67,68^, drought stress ^69^, and heat stress ^70^. Mutant plants are also found extremely dwarfing ^67^. Another gene, *YCF4*, is involved in the assembly and stability of photosystem I ^71^.Among genes under directional selection, six of nine annotated genes are found for the response to stimulus (water, hypoxia, radiation, ABA, and jasmonic acid, e.g., *HUP44*, *AT3G16210,* *AT5G49525*, and *P1R3*). (**Table S20**).

For candidate genes for local adaptation, gene ontology analysis showed that three major biological processes were significantly enriched, including 1) response to environmental stimulus; 2) plant growth, development and reproduction processes; 3) biosynthesis and metabolism for compounds (see **Table S21** for the detailed list). Genes in the region Chr.5: 89.89 Mb ~ 90.46 Mb were found related to the regulators of various growth and development processes, and response to stimulus in plants (**Table S22**). Particularly, *CB5-E* was involved in response to salt, cold and multiple phytohormone in Chinese Cabbage^72^. *UBQ1* was involved in response to heat in tomato ^73^.

For 102 genes under selective sweeps, 13 GO categories were significantly overrepresented with *p*-value < 0.05 (Fisher’s exact test, see **Table S23**), mainly involved in photosynthesis, metabolites biosynthesis, and root development. For example, the *FTSH5* is essential for photosystem II repair and chloroplast biogenesis^74^. Three genes (*MAIL1*, *AMT1;1*, *CIAF1*) were located in Chr.7: 34.3 Mb - 36.6 Mb region. *MAIL1* is essential for development of the primary root^75^. Loss of function mutations display extremely short primary root, leaf development is later and smaller^75,76^. The gene *AMT1;1* is involved in lateral root branching. *AMT1;1* encodes a plasma membrane-localized ammonium transporter protein and gene is required for ammonium uptake in poplar ^77^. The overexpression mutant significantly improves salt tolerance during the early root growth stage after seed germination by alleviating ammonia toxicity caused by salt stress^78^. The gene *CIAF1* is related to mitochondrial respiratory chain complex I assembly. And mitochondrial complex I is necessary to maintain optimal photosynthetic performance ^79,80^. Another gene *TCP1* was located in Chr.7:36.7 Mb - 38.3 Mb region and involved in the longitudinal elongation of leaves ^81,82^.

**Methods S5 Environmental factors**

To assess the correlation between genetic differentiation and environment variables, we first obtained soil salt conditions (SQ5) from Harmonized World Soil Database v 1.2(<https://www.fao.org/soils-portal/soil-survey/soil-maps-and-databases/harmonized-world-soil-database-v12/en/>). Eighty-four environmental and 19 climate variables for all sampling sites were extracted from the WorldClim dataset (<http://www.worldclim.org/>) using the R package ‘raster’ (<https://cran.r-project.org/web/packages/raster/index.html>) and averaged over the period from 1970-2000. We classified them into five basic categories: 1) temperature related factors, 2) precipitation related factors, 3) wind speed related factors, 4) solar radiation related factors and 5) water vapor pressure related factors. Principal component analysis (PCA) was performed for the factors in each of five categories using the R package ‘FactoMineR’^83^. PC components with a cumulative contribution larger than 99% were retained as representatives of that category for association analyses between genetic differentiation and environment. In total, of 27 representative components were retained and used to calculate five environmental distance matrices using the R package ‘cluster’^64^. An overall environmental distance matrix *E* was also generated using the same method but based on all 104 environmental and climatic factors (including SQ5). Additionally, the aridity index (A. I.), the ratio of annual precipitation to annual potential evapotranspiration (PET), were obtained using the ‘Thornthwaite’ function implemented in the R package ‘SPEI’^84^ and averaged across the years 2000 to 2018. The aridity index, longitude and latitude were also used as three independent factors for association analyses between genetic divergence and environmental changes. See all information about all environmental and climatic factors in **Table S16**.

**Methods S6 Linkage disequilibrium**

Phased haplotypes and imputed missing genotypes were performed using Beagle v5.2^85^. Pairwise linkage disequilibrium (*r*^2^) was calculated using emeraLD v.0.1^86^ with a sliding window of 10 kb.

**Supplementary Figures**

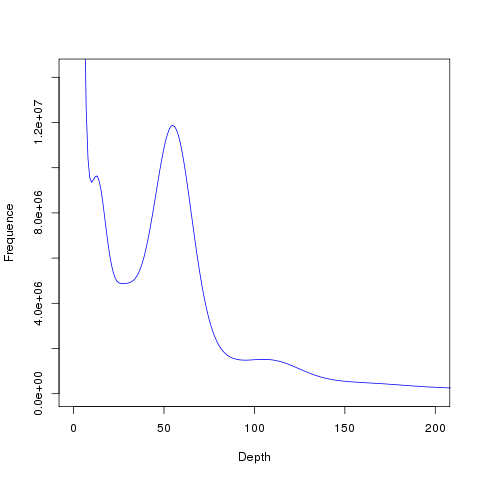

**Fig. S1** *k*-mer analysis for *G. przewalskii* genome size estimation. The x-and y-axes show *k*-mer depth and *k*-mer frequency, respectively.

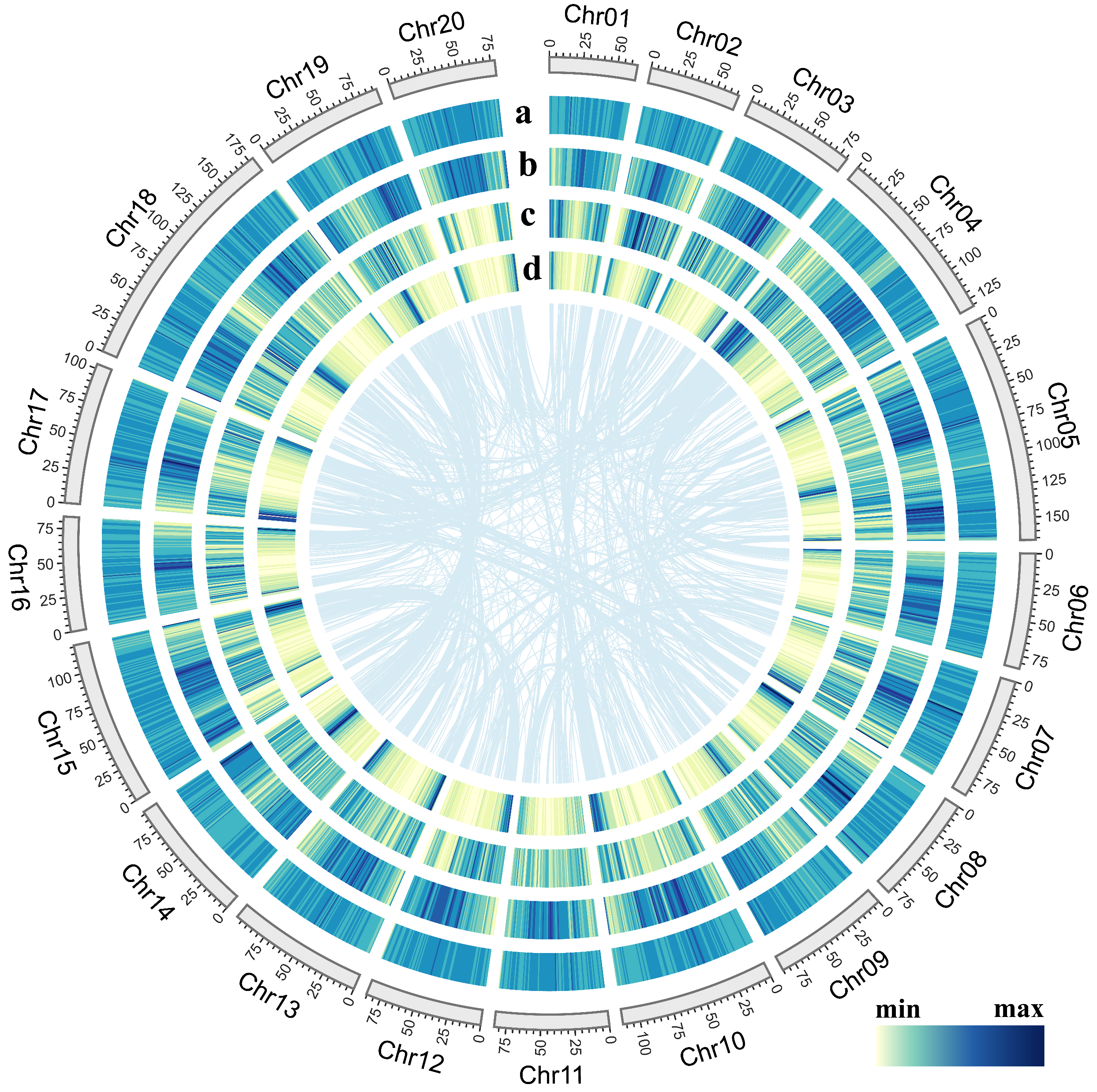

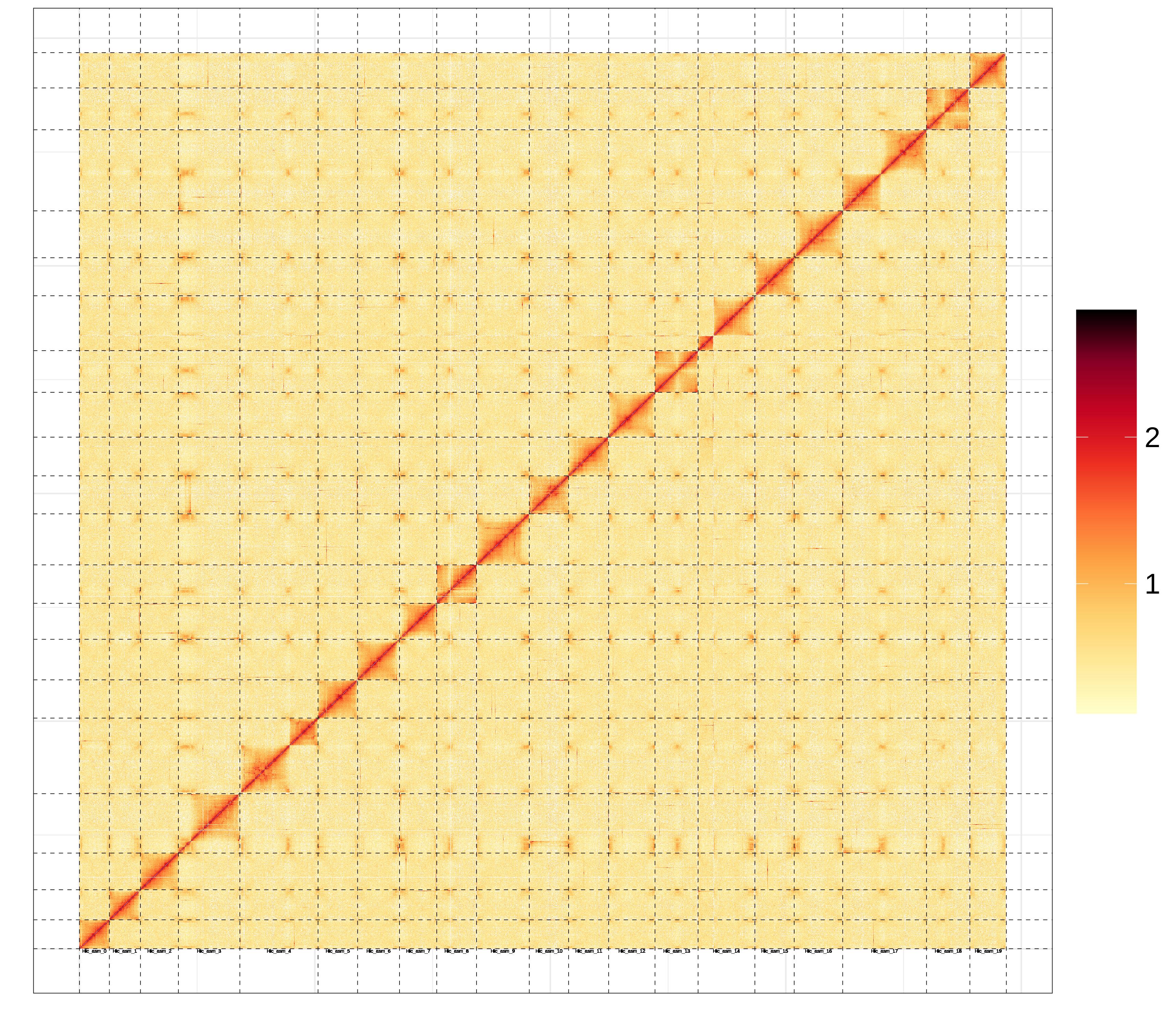

**Fig. S2** (Left) Circos plot of 20 chromosomes of *G. przewalskii* reference genome, showing the density distributions of (a) repeat element; (b) GC content; (c) SNP number; (d) gene density. The four metrics are calculated in 1 Mb sliding windows. The synteny blocks are shown with lines connecting genes in the center. (Right) Hi-C interaction heat map for *G. przewalskii* reference genome showing interactions between 20 chromosomes.

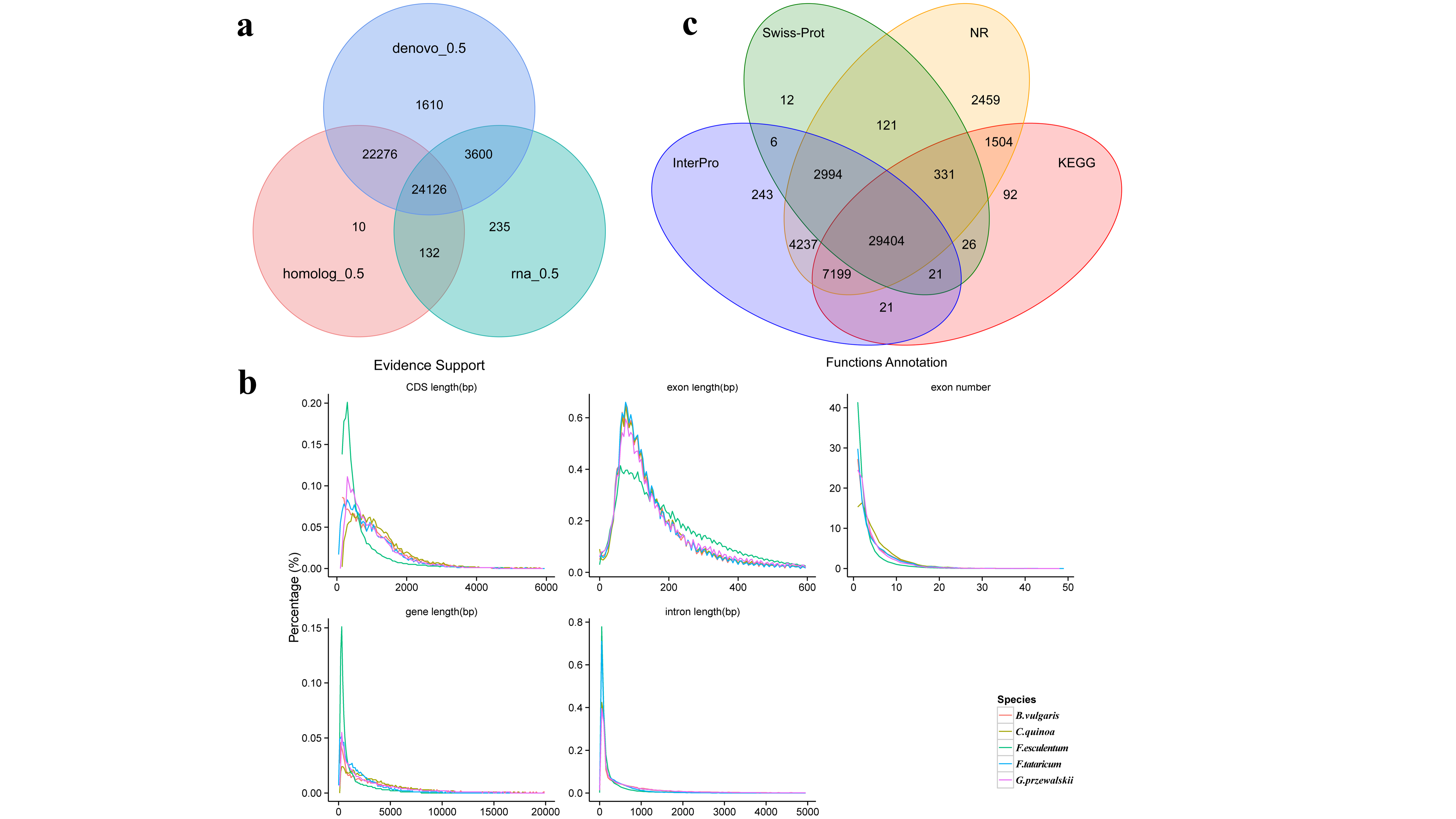

**Fig. S3** (a) Venn diagram shows that most of predicted gene models are supported by more than one method. (b) Distributions of gene models from *G. przewalskii* and four other species. (c) Venn diagram of functional annotations performed by four databases.

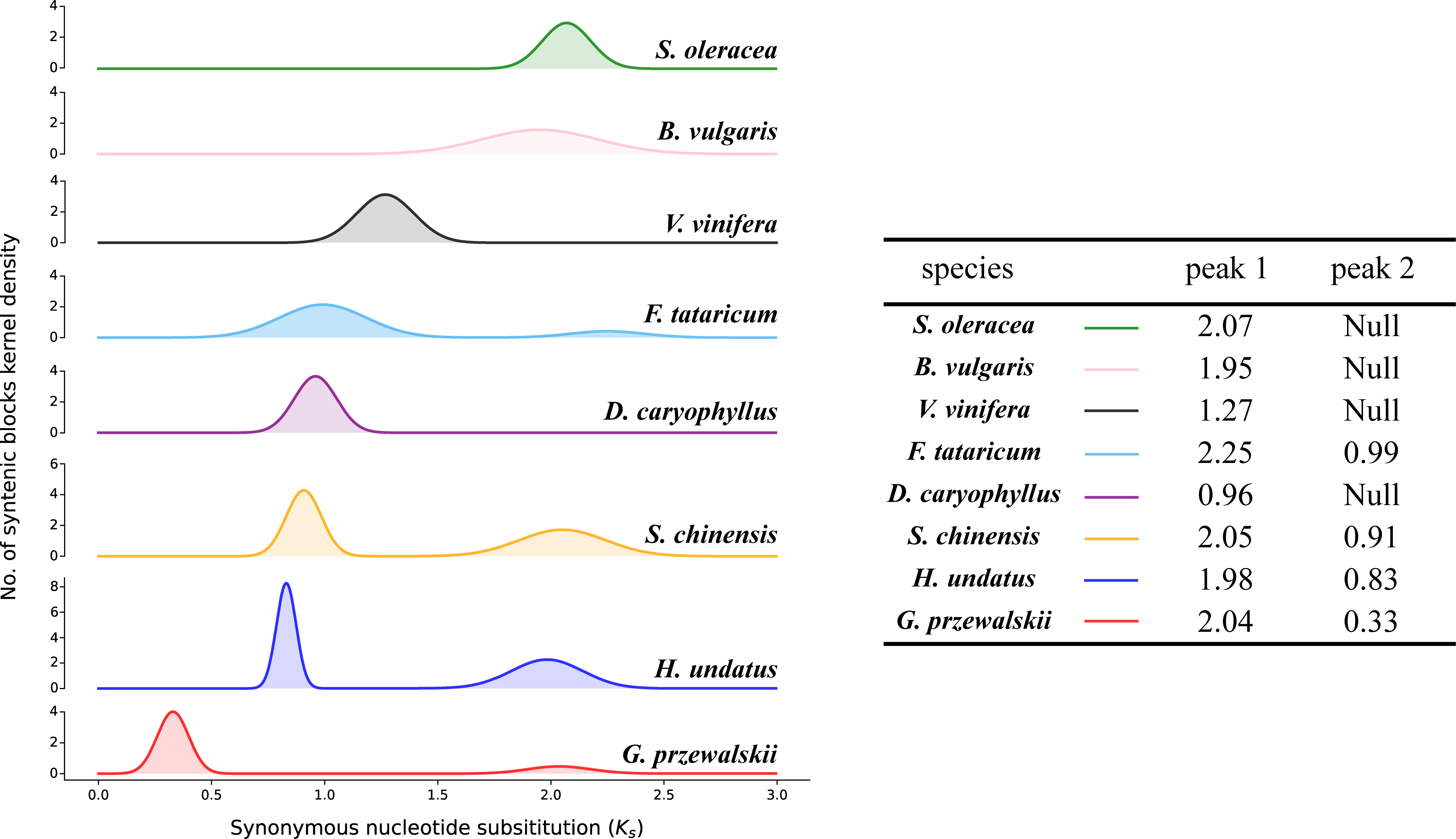

**Fig. S4** Distribution of Synonymous substitution rate (*K_s_*) for paralogous genes within the genomes of *G. przewalskii*, *Hylocereus undatus*, *Simmondsia chinensis*, *Spinacia oleracea*, *Vitis vinifera*, *Beta vulgaris, Dianthus caryophyllus,* and *Fagopyrum tataricum*.

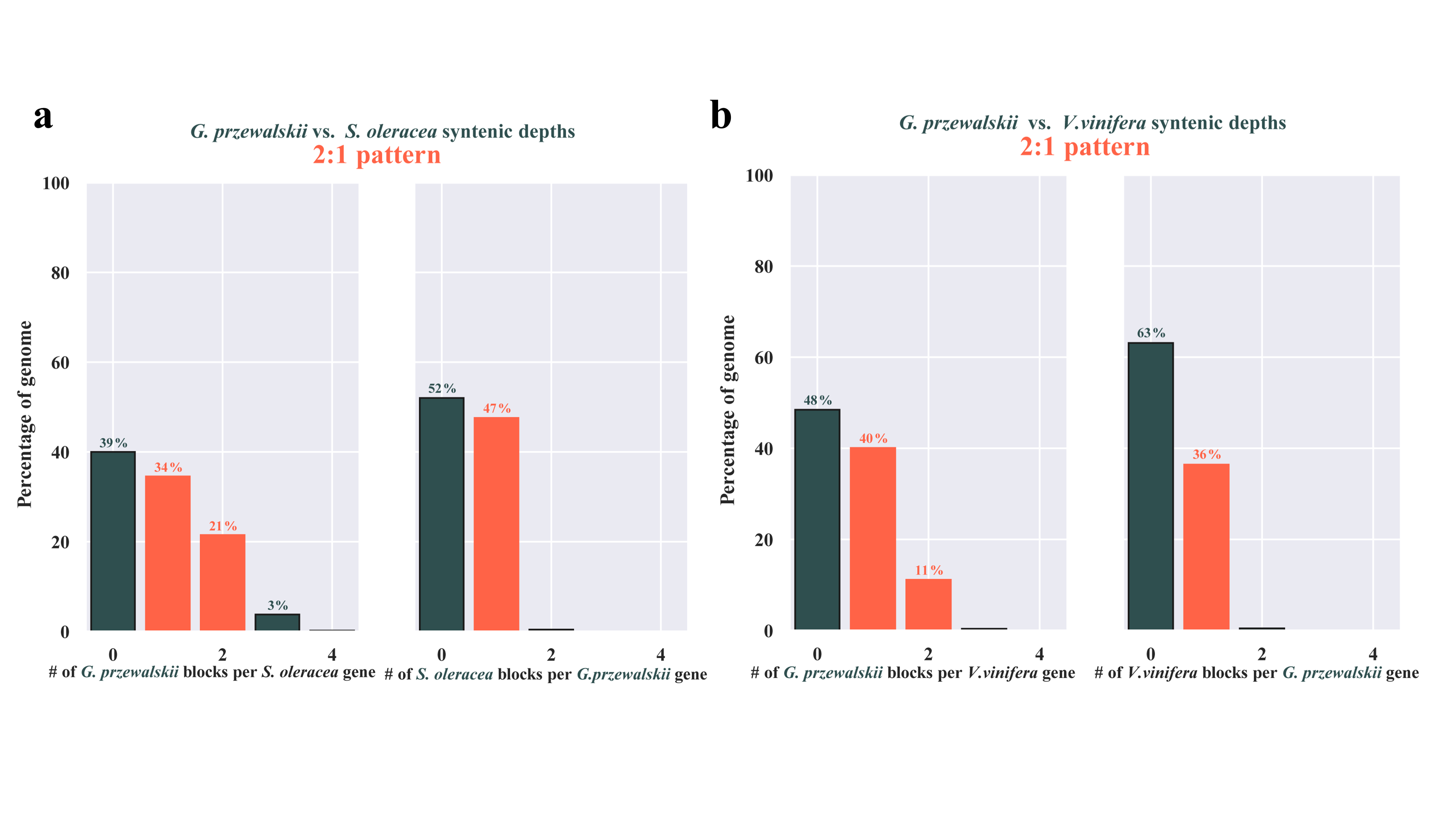

**Fig. S5** The plot of syntenic depths in *G. przewalskii*. (a) *G. przewalskii* vs. *S. oleracea* genome; (b) *G. przewalskii* vs. *V. vinifera* genome.

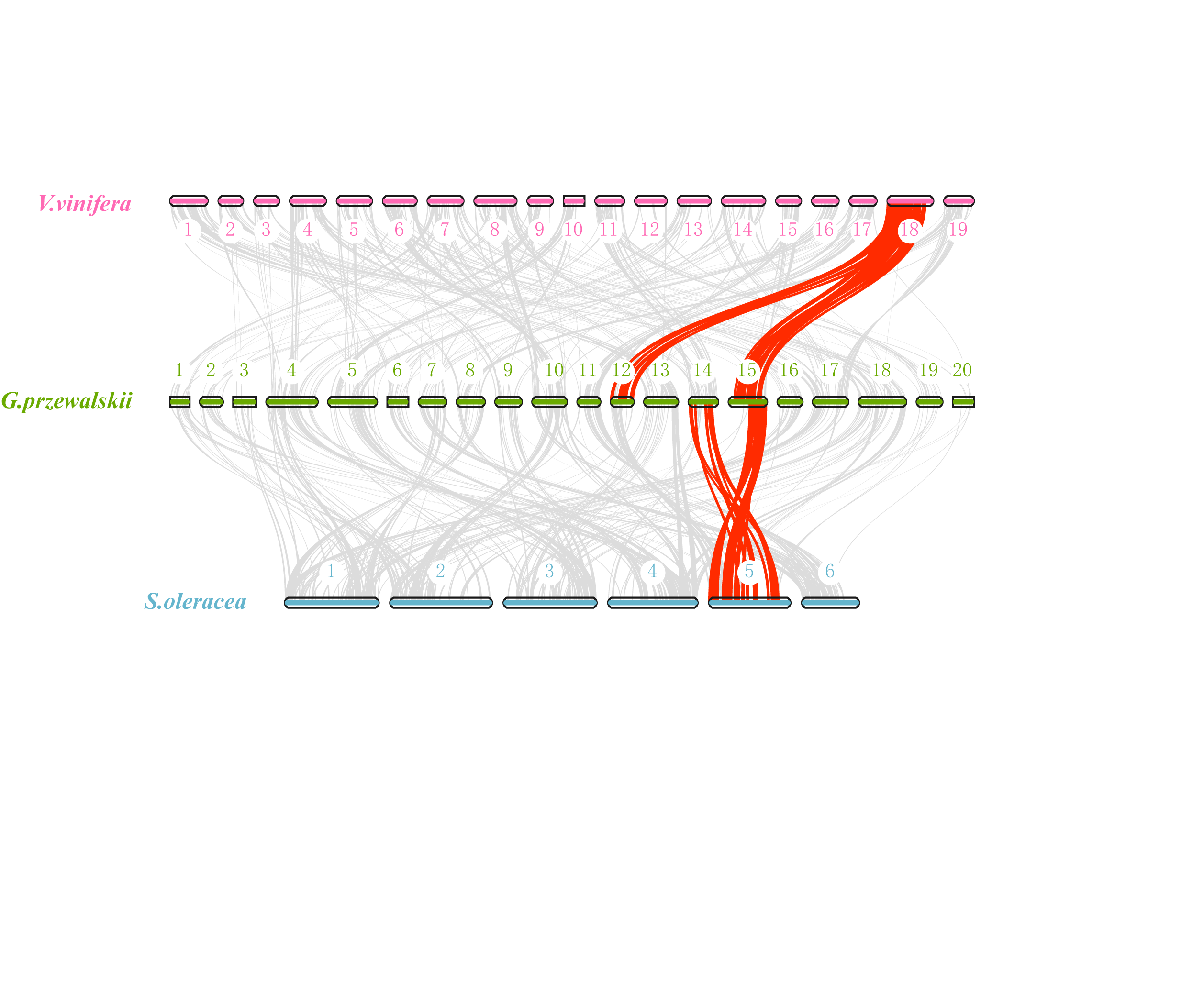

**Fig. S6** Macrosynteny between genomic regions of *G. przewalskii*, *S. oleracea* and *V. vinifera*. Synteny patterns for 2:1 between *G. przewalskii* and two reference genomes support the independent WGD event of *G. przewalskii*.

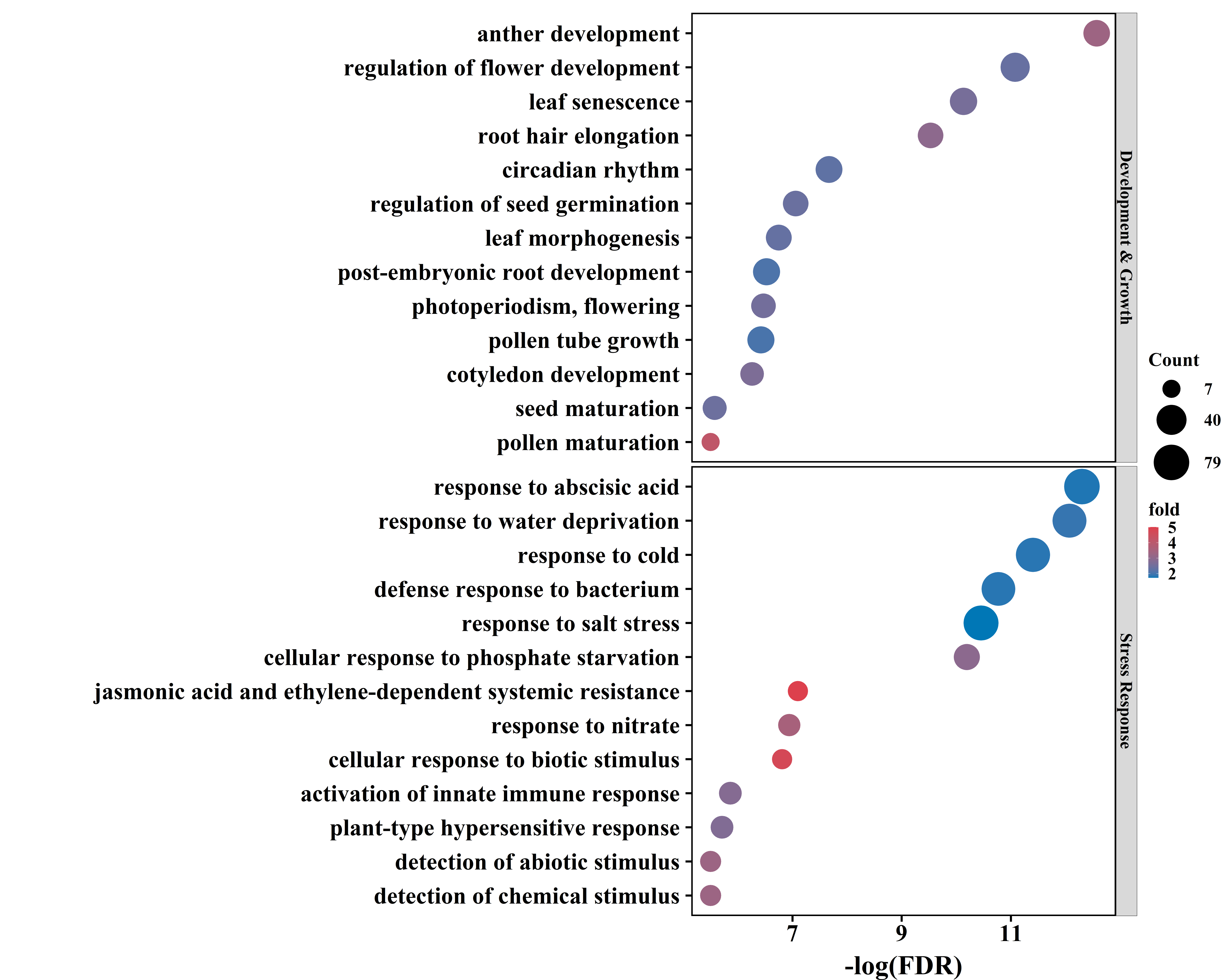

**Fig. S7** Gene ontologies (biological processes) for 7,378 duplicated genes for recent WGD associated with expansion gene family. Only those of stress response, growth and developmental processes are shown.

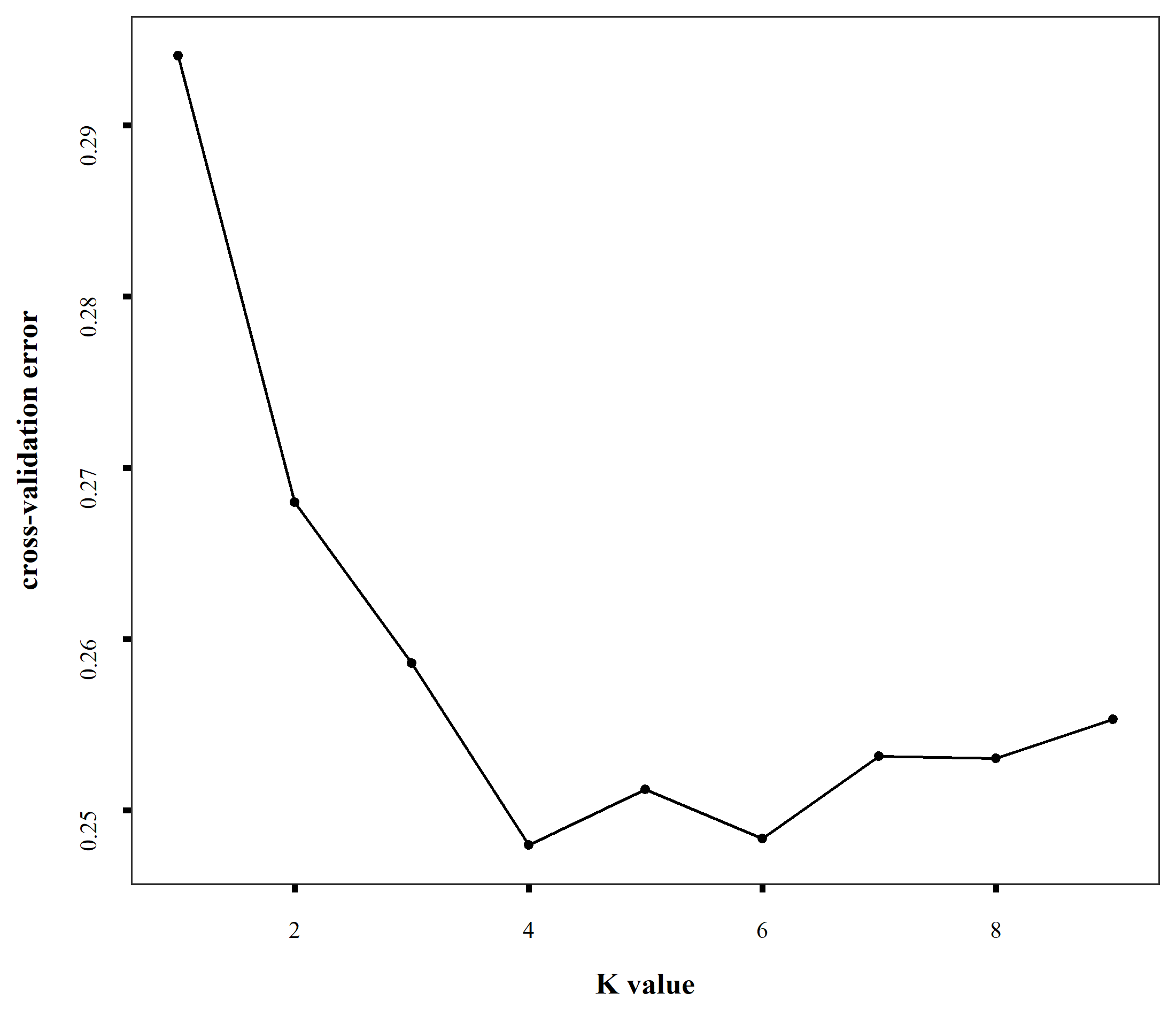

**Fig. S8** The cross-validation errors against *K* values in Admixture analysis.

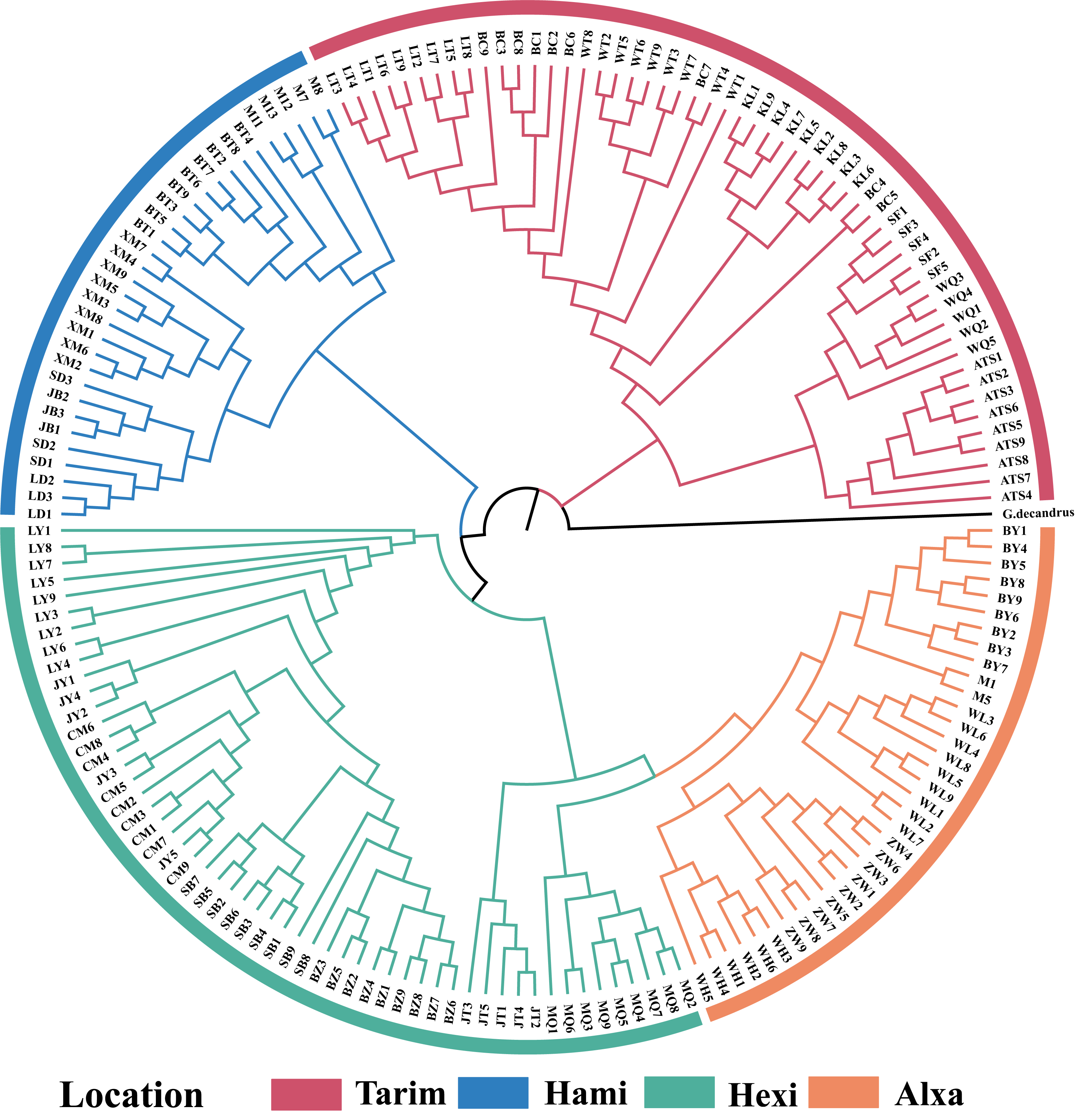

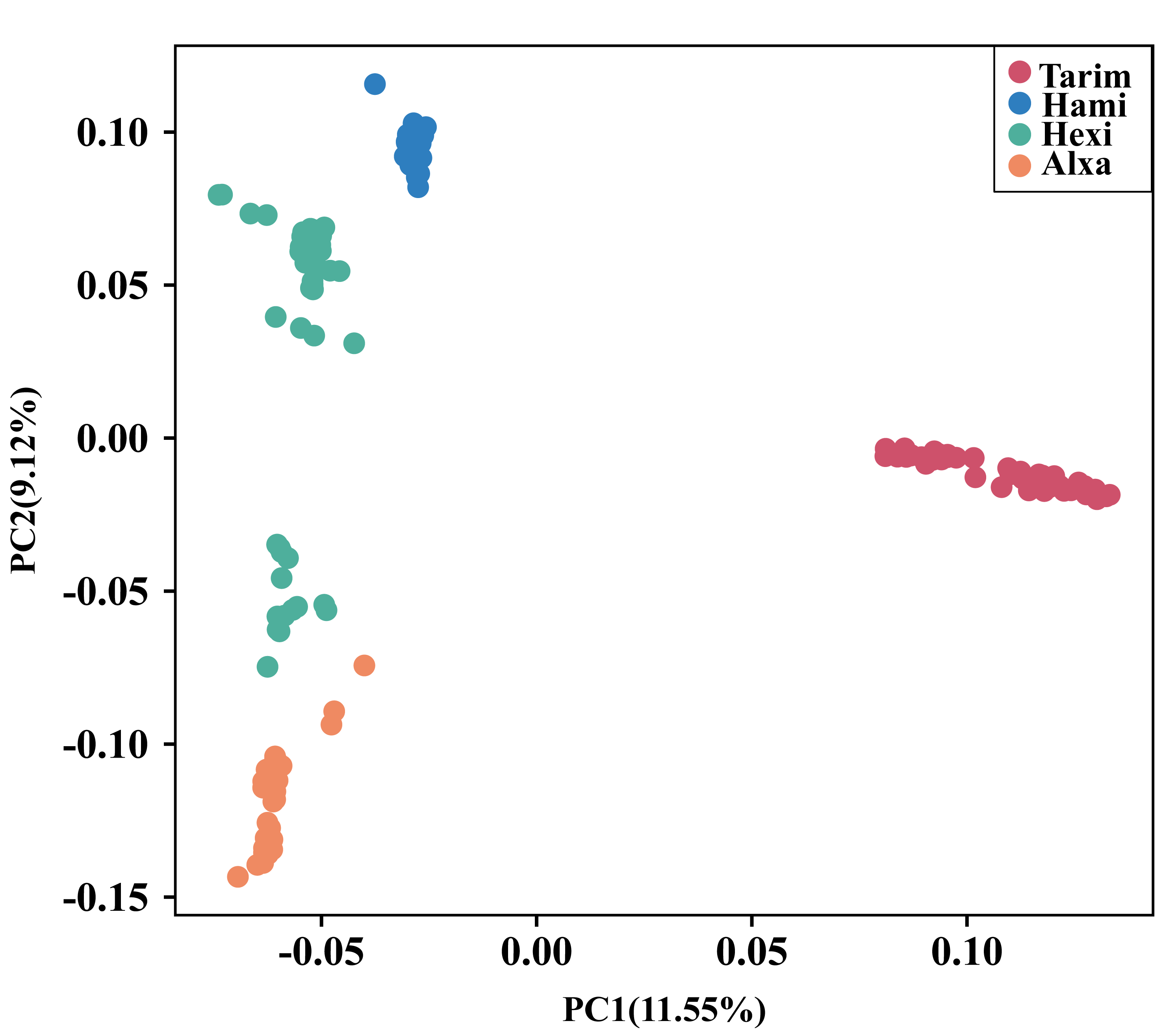

**Fig. S9** Population structure of *G. przewalskii*. left) Neighbor Joining phylogenetic tree based on intergenic single-nucleotide polymorphisms (SNPs) with LD removed, using *G. decandrus* as outgroup. right) Principal component analysis (PCA) with the first two principal components. Colors correspond to the phylogenetic tree grouping.

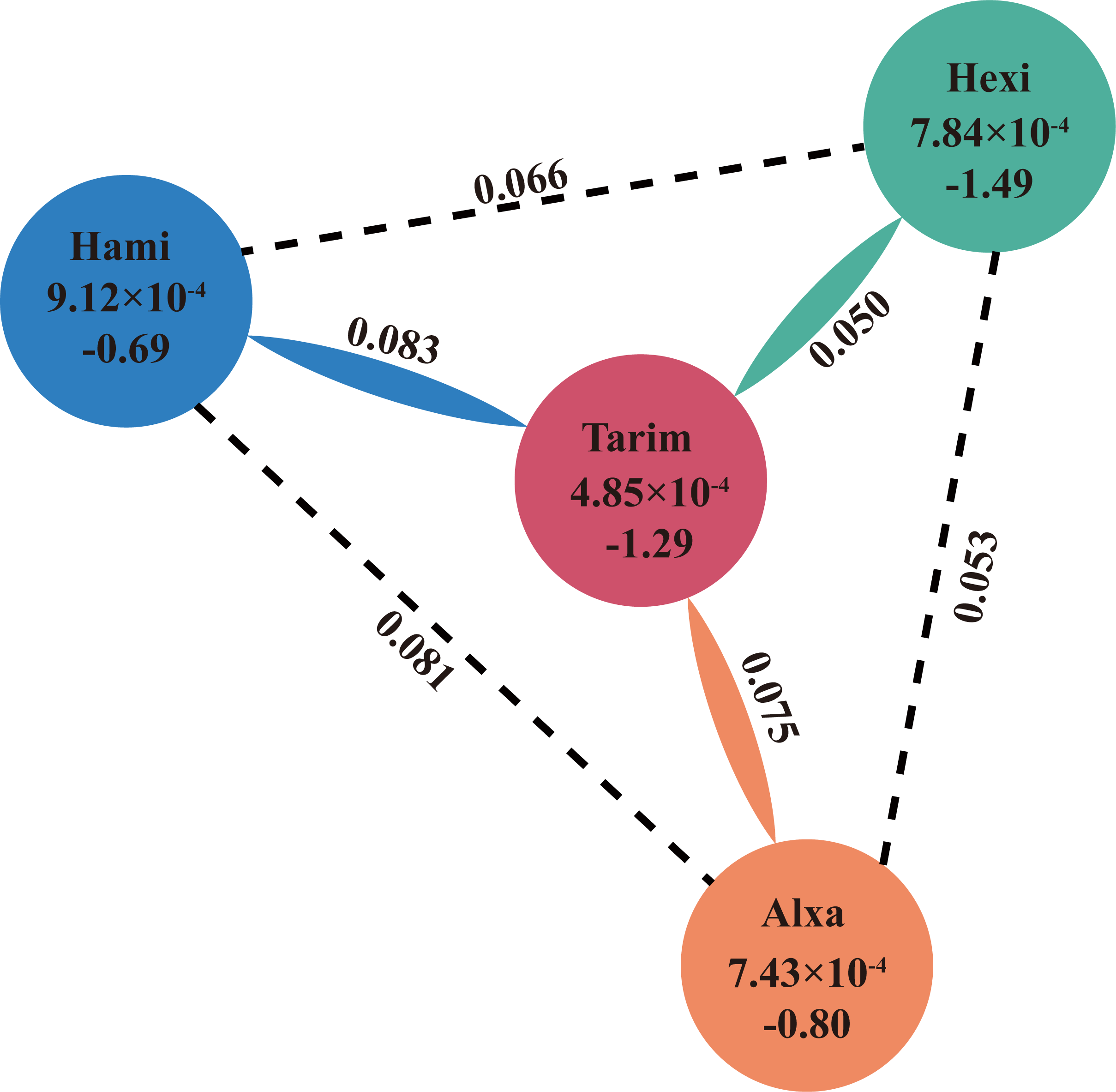

**Fig. S10** The genetic diversity (π), neutrality test statistic (Tajima’s *D*) and divergence (*F*_ST_) of four major lineages. Values above the inside of the circle represent nucleotide diversity (π) for the lineage, Values below the inside of the circle represent Tajima’s *D*, and values between lines indicate population divergence (*F*_ST_).

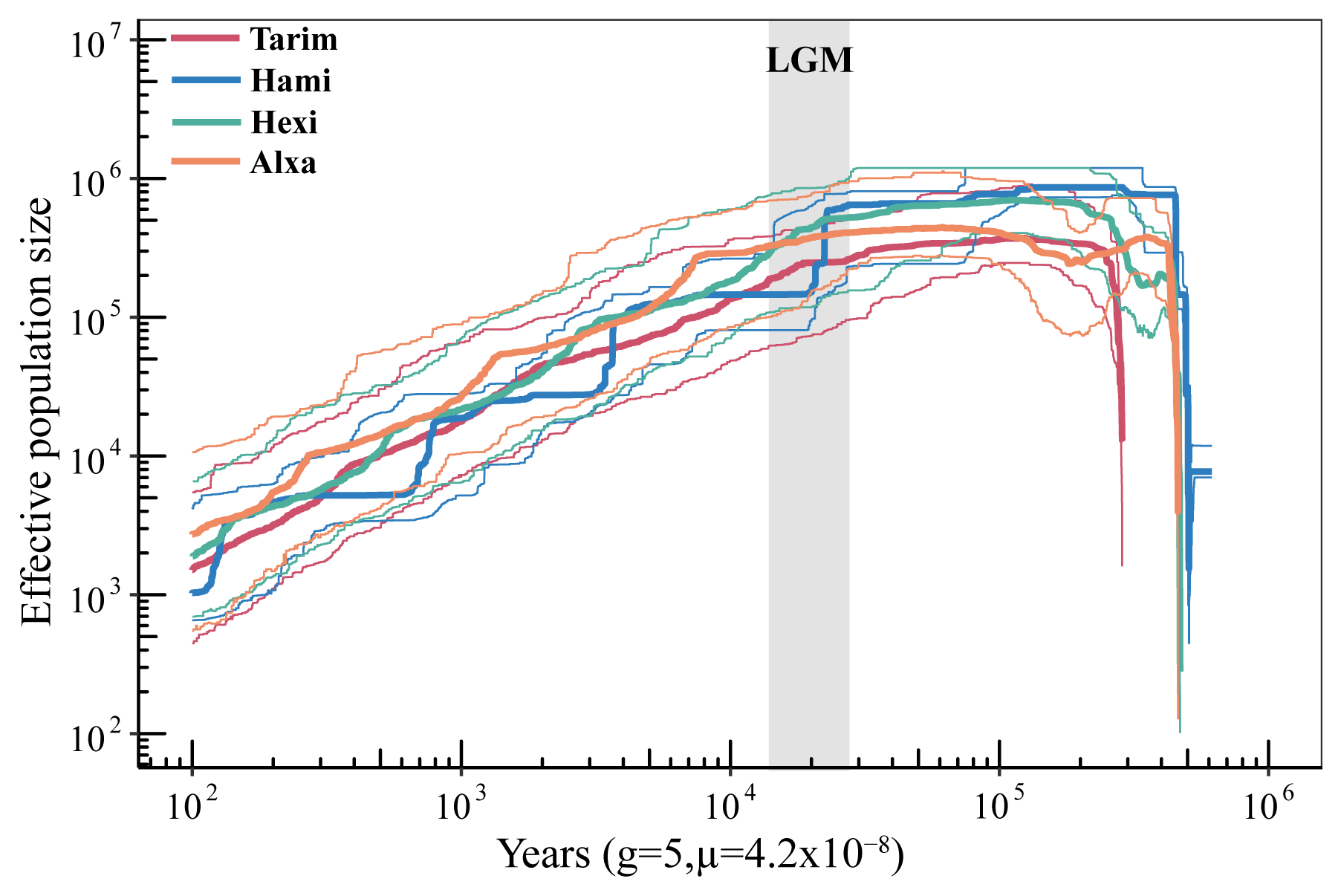

**Fig. S11** The result of Stairway Plot2 on all lineages for *G.przewalskii.* The thick lines represent the median values of population sizes and the thin lines show the 5% and 95% confident intervals.

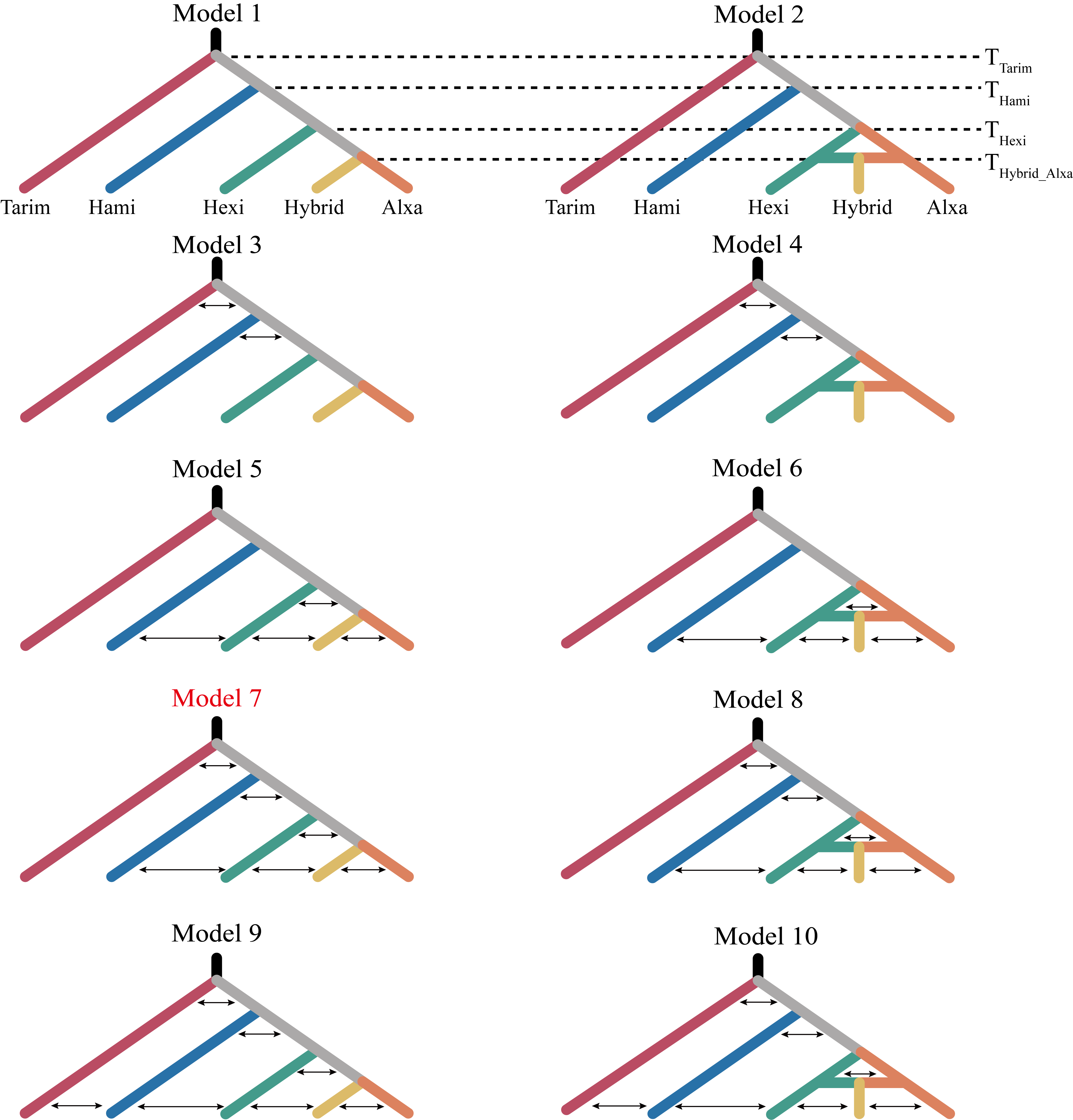

**Fig. S12** Schematic diagram of ten priori demographic models with different migration events for *G. przewalskii* by fastsimcoal2.

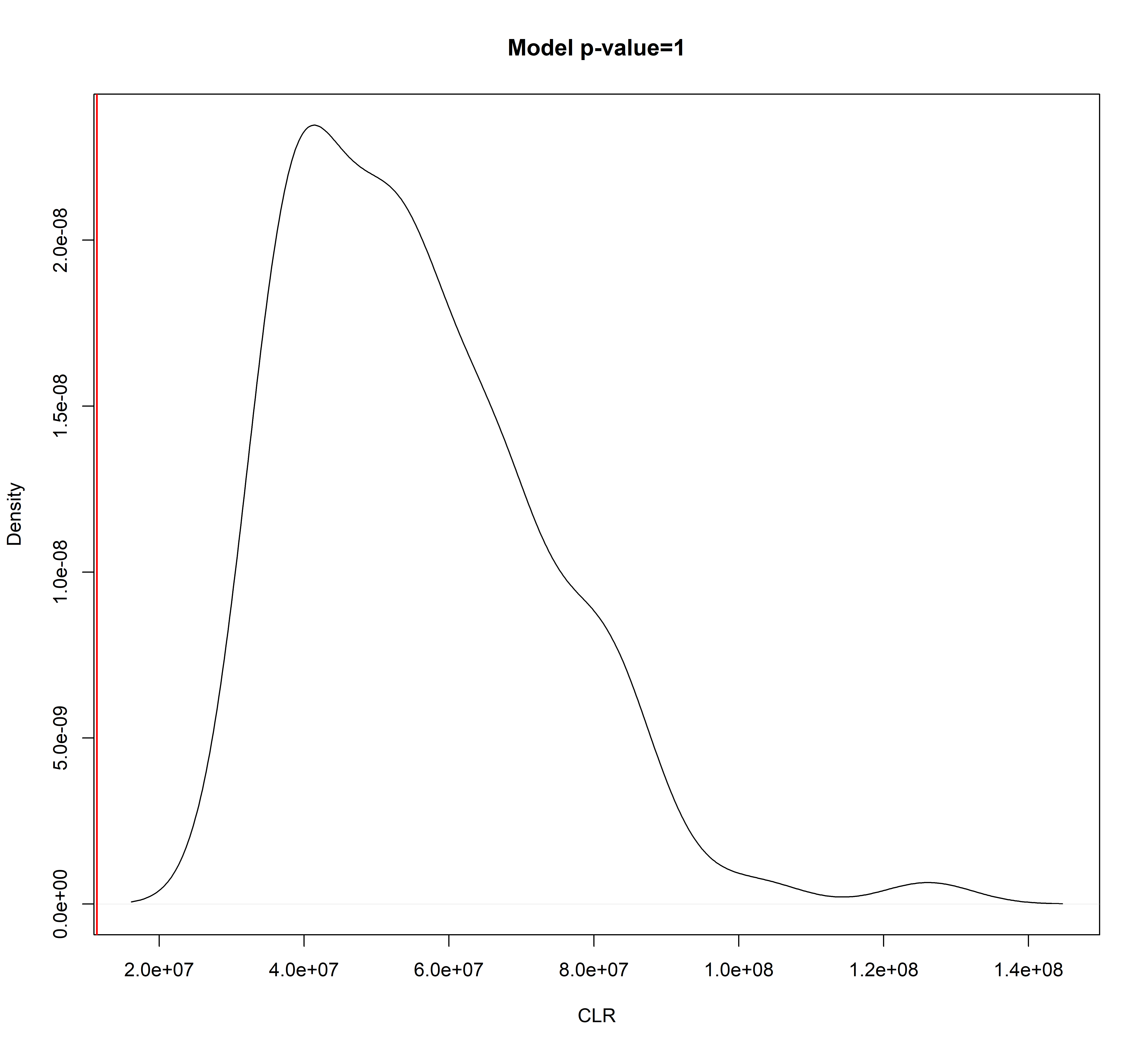

**Fig. S13** Likelihood ratio G-statistics distribution. The likelihood ratio G-statistics (*CLR* = log10(*CL*_O_/*CL*_E)_, where *CL_O_* and *CL_E_* are the observed and estimated maximum composite likelihood, respectively) was checked how data fit the chosen model. A non-significant *p*-value of this test suggests that the observed SFS is well compatible with the model. The red line is the observed CLR.

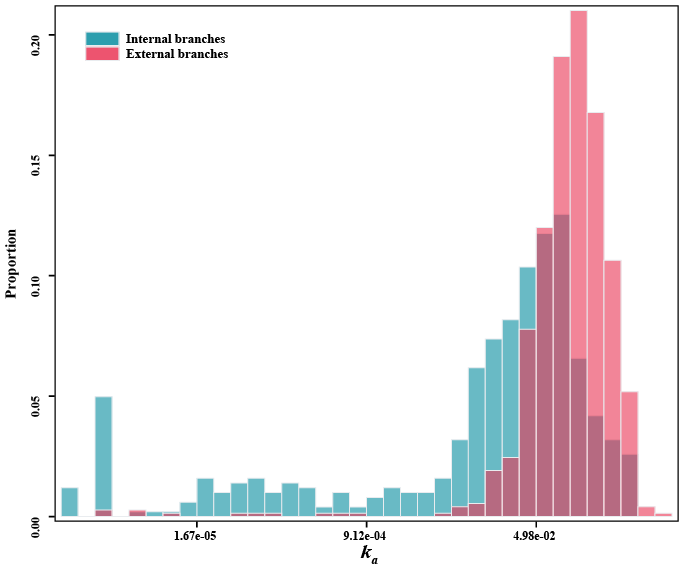

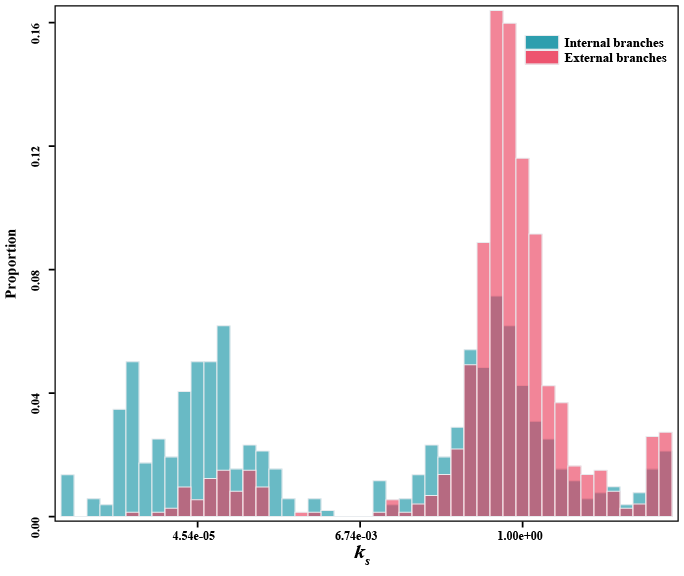

**Fig. S14** The distribution of *k_a_* (left), *k_s_* (right) for “common xerophytic” genes.

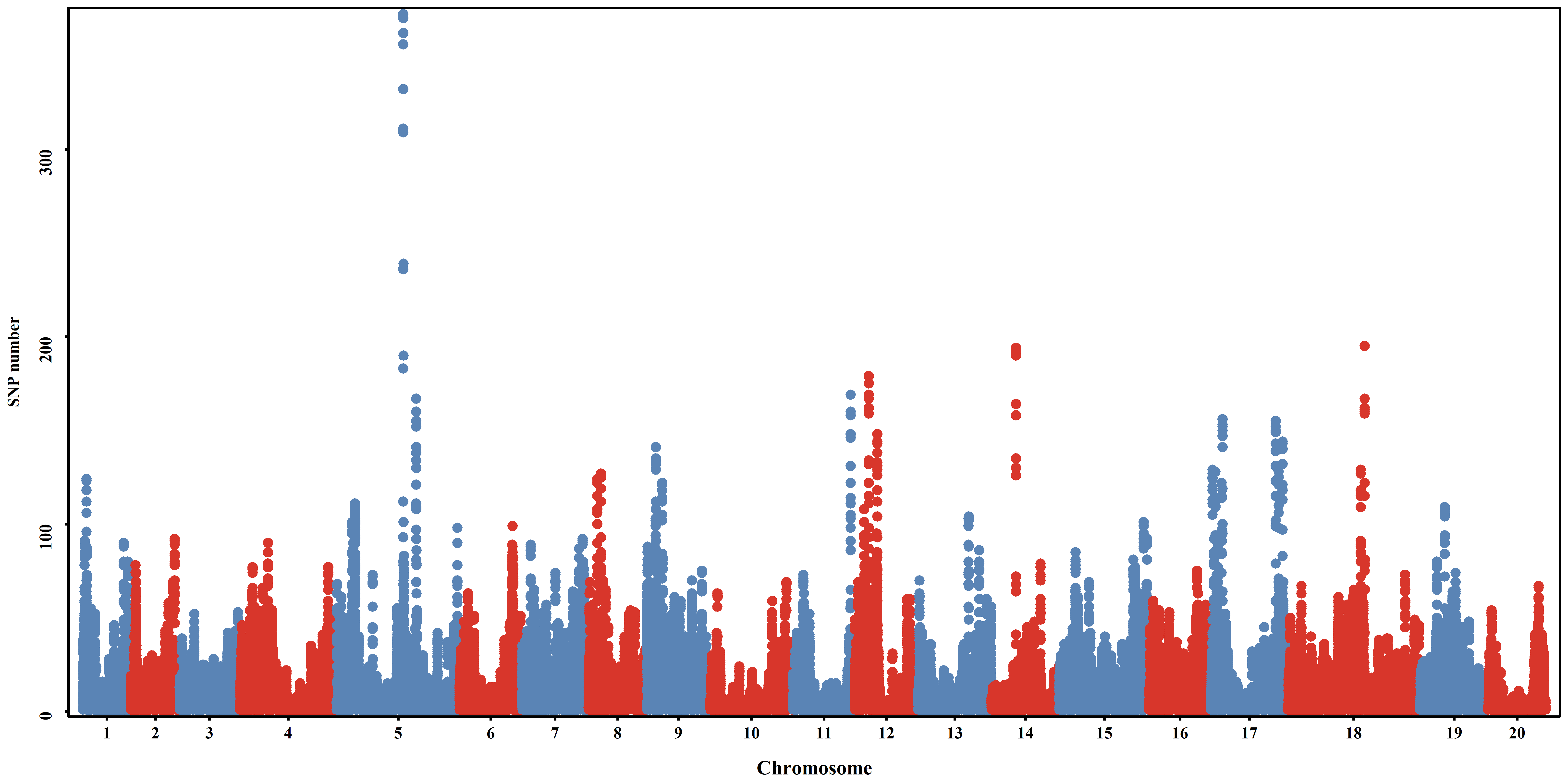

**Fig. S15** The distributions of the number of SNPs under diversifying selection for 20 chromosomes. Each dot shows the SNP number for 100 kb windows sliding in increments of 10 kb.

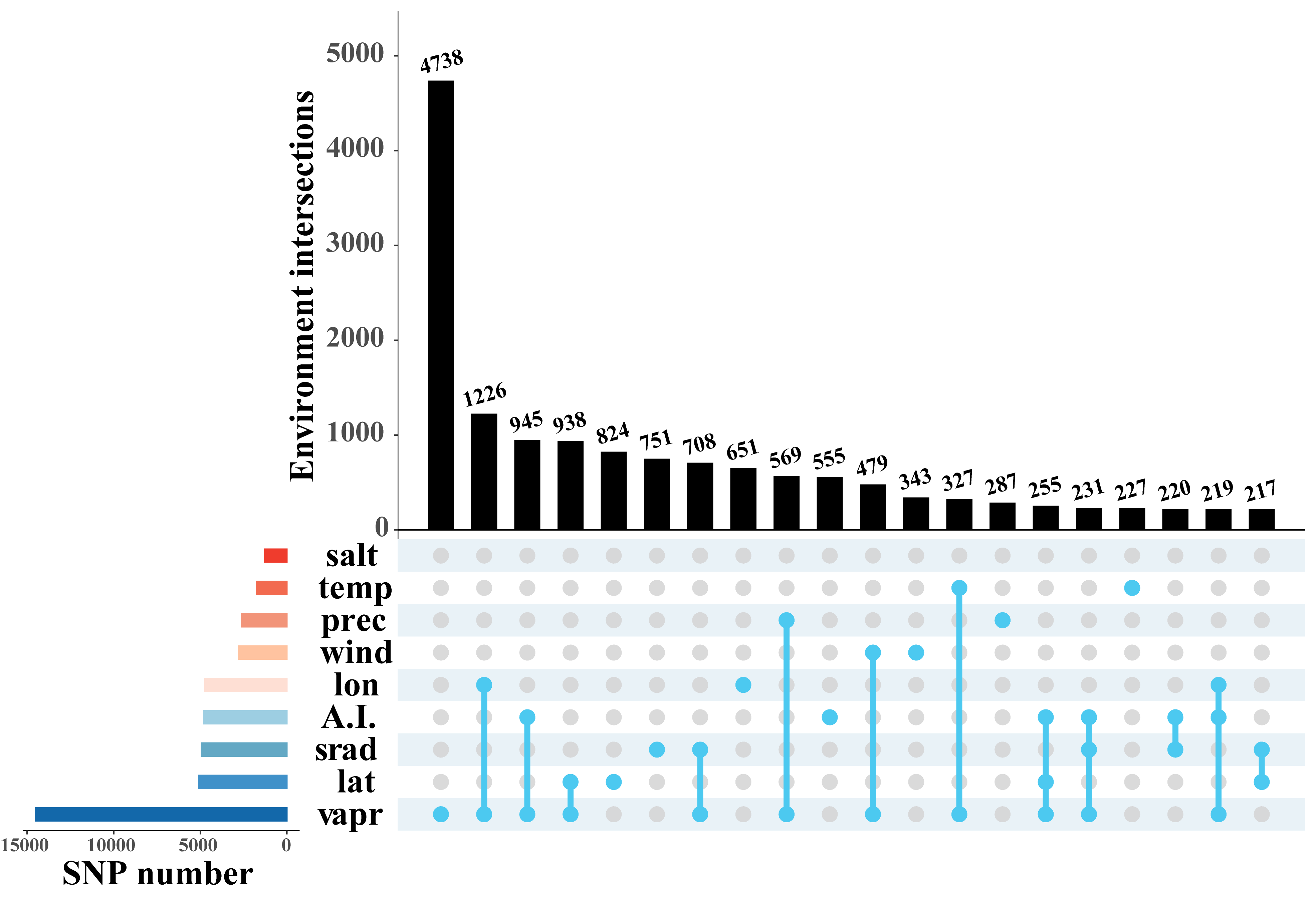

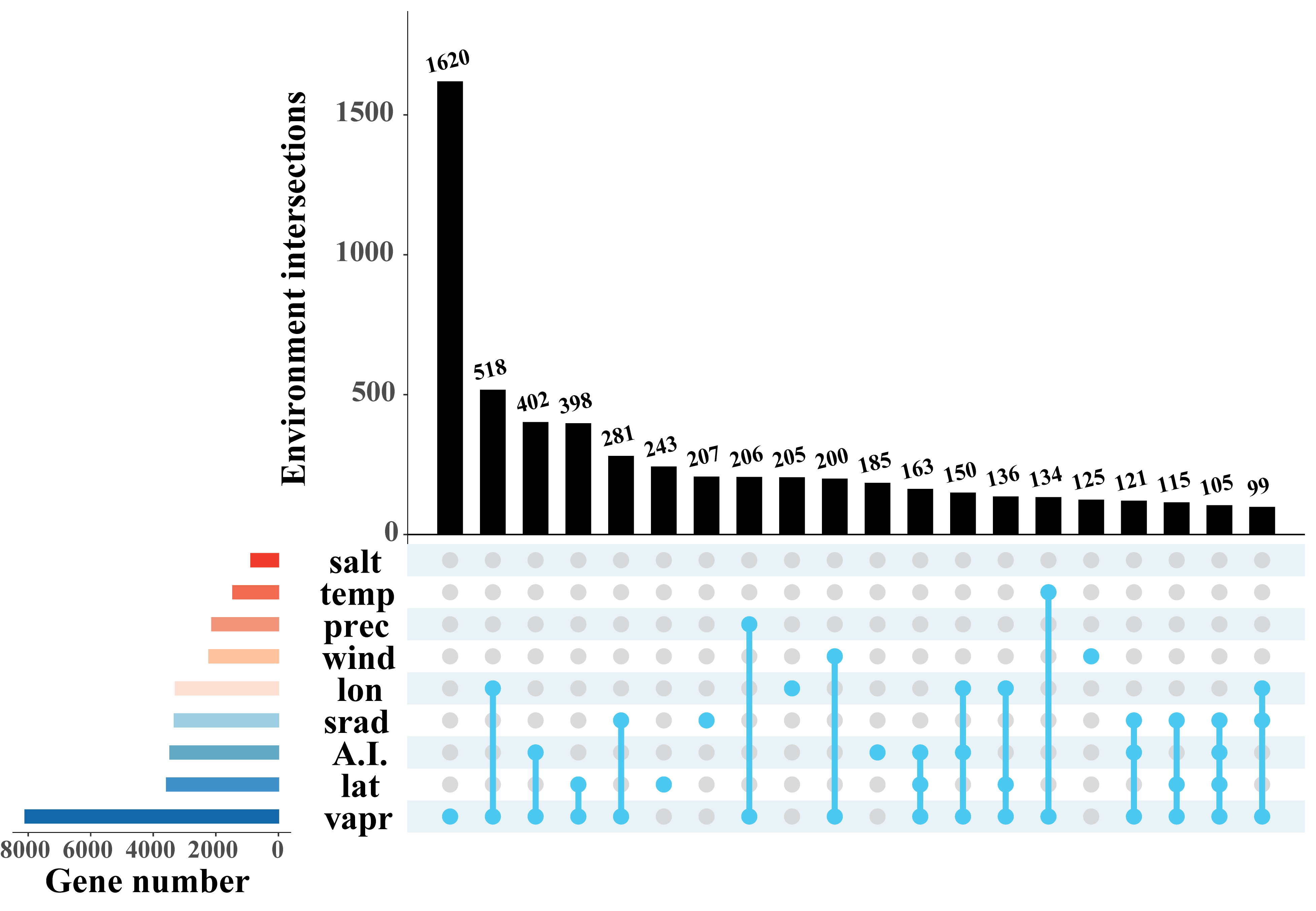

**Fig. S16** The distributions of the number of SNPs and genes under diversifying selection for different environmental factors. (Left) Distribution of number of SNPs associated to environmental factors in protein-coding genes. (Right) Distribution of the number of genes associated to environmental factors. Vertical black bars indicate SNPs or genes that are associated to various environmental factors. Horizontal colored bars correspond to SNPs or genes associated to one environmental factor. salt, salinity; temp, temperature; prec, precipitation; wind, wind speed; lon, longitude; A.I., aridity index; srad, solar radiation; lat, latitude; vapr, water vapor pressure.

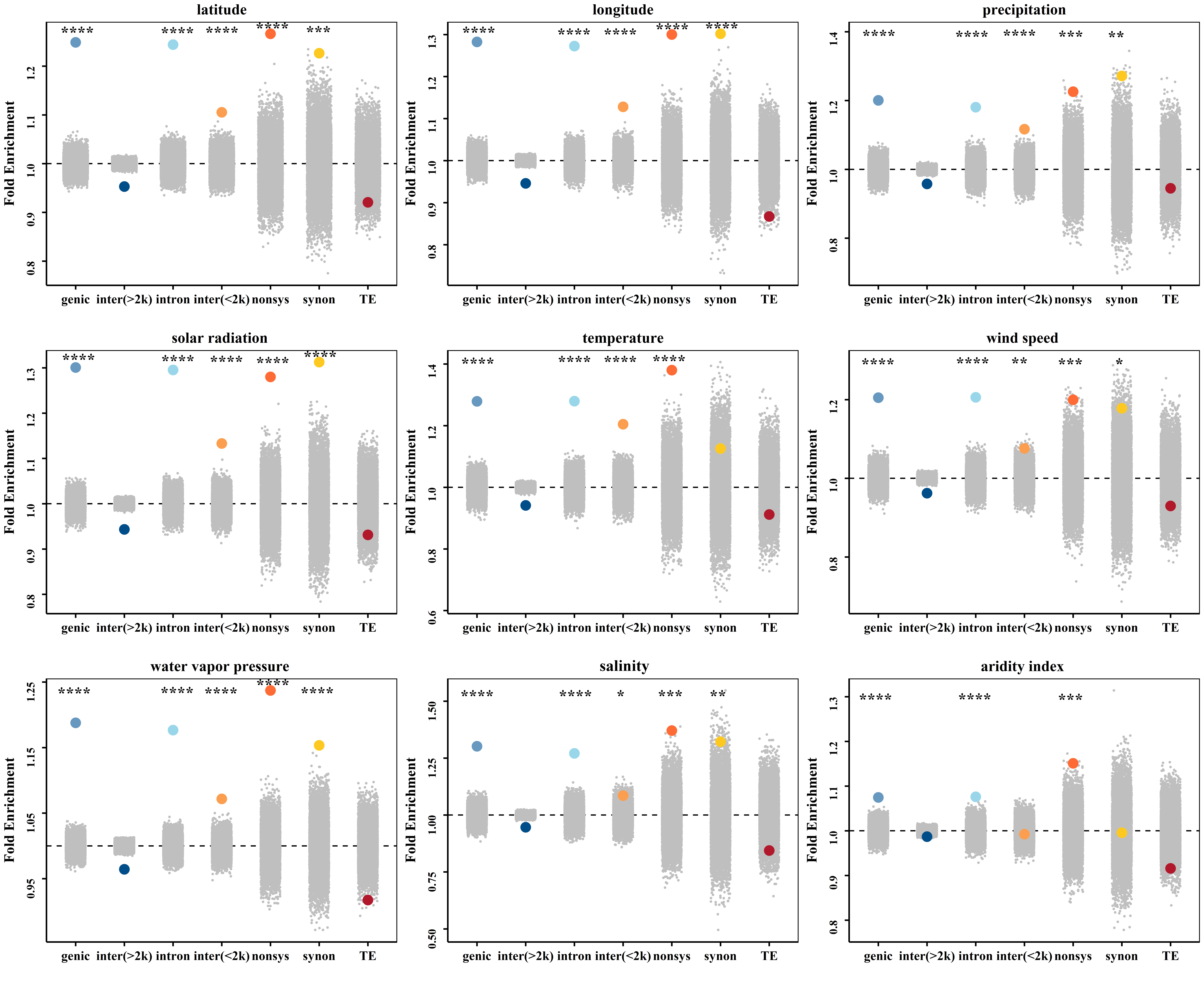

**Fig. S17** Enrichment analysis of different classes of SNPs under diversifying selection for nine environmental factors. Enrichments shown are relative to the proportion of each class of SNPs in the genome overall. The horizontal dashed line shows the expected enrichment under the null hypothesis of no enrichment. The colored dots show the enrichment values for each class of SNPs. Gray dots represent 10,000 null permutations of site categories. Statistical significance was determined by Fisher’s exact test (*p*-values: * < 0.05, ** < 0.01, *** < 0.001, **** < 0.0001. genic, SNPs in genic regions; intron, intronic SNPs; inter(>2k), SNPs in distal regions (larger than 2 kb to 5’ or 3’ end); inter(<2k), SNPs in proximal regions (within 2 kb flanking 5’ or 3’ end); nonsys, nonsynonymous SNPs (amino acid changing SNPs); synon, synonymous SNPs; TE, Transposable element.

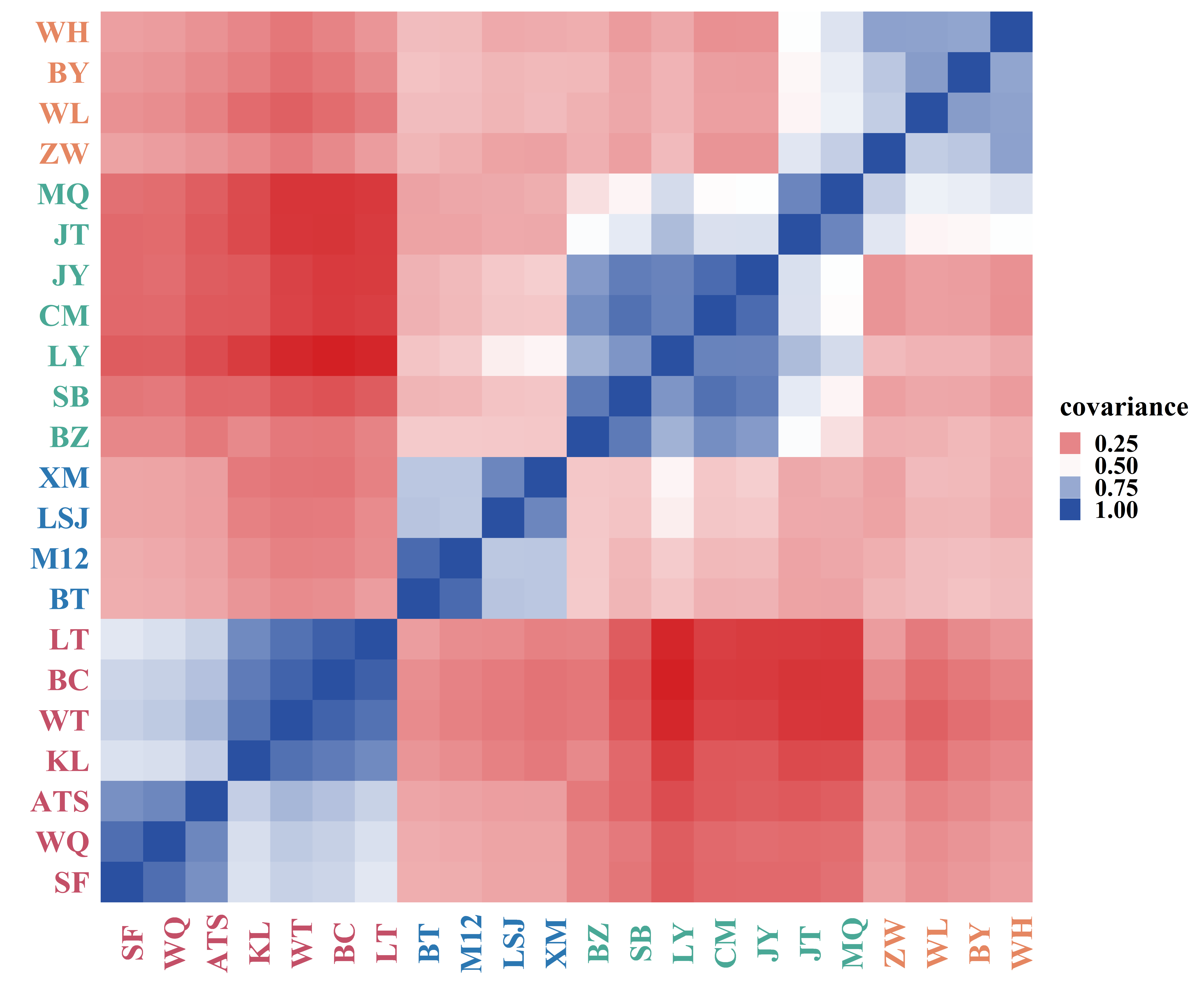

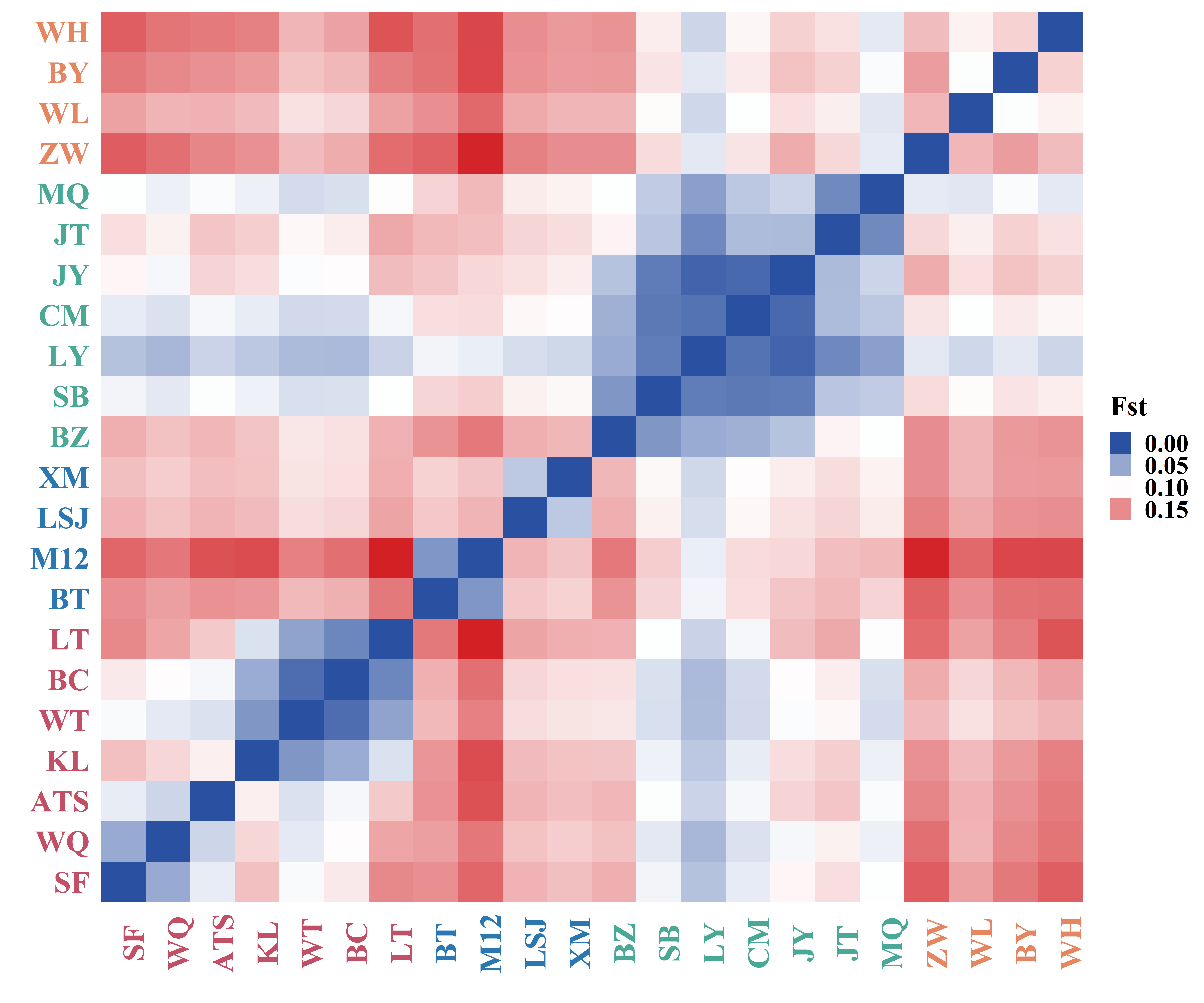

**Fig. S18** The covariance matrix of *G. przewalskii* populations (left) and *F*_ST_ matrix of *G. przewalskii* populations (right).

**Supplementary Tables**

**Table S1** *G. przewalskii* reference genome sequencing library statistics.

| **Pair-end libraries** | **Insert size** | **Total data (G)** | **Read length (bp)** | **Depth (X)** |
| --- | --- | --- | --- | --- |
| **Illumina reads** | 350bp | 208 | 150 | 90.04 |
| **PacBio reads** | / | 385 | / | 166.67 |
| **Total** | / | 593.00 | / | 256.71 |

**Table S2** Estimation of the *G. przewalskii* genome size based on 17-mer statistics.

| **K-mer** | **K-mer number** | **K-mer Depth** | **Genome size (Mb)** | **Revised genome size (Mb)** | **Used Base (Mb)** | **Repeat** **_rate(%)** | **Heterozygous_rate(%)** |
| --- | --- | --- | --- | --- | --- | --- | --- |
| 17 | 130,281,321,749 | 56 | 2,326.45 | 2,310.3 | 208,139.9 | 78.43 | 0.91 |

**Table S3** Summary of contigs and scaffolds for *G. przewalskii* reference genome.

|  | **length** | | **number** | |
| --- | --- | --- | --- | --- |
|  | **contig** | **scaffold** | **contig** | **scaffold** |
| **N50** | 705,529 | 91,849,704 | 848 | 9 |
| **N60** | 559,559 | 85,552,657 | 1,180 | 11 |
| **N70** | 431,901 | 83,722,907 | 1,608 | 14 |
| **N80** | 301,017 | 80,826,546 | 2,188 | 16 |
| **N90** | 176,532 | 66,518,824 | 3,083 | 19 |
| **Max** | 4,740,395 | 178,545,894 | / | / |
| **Total** | 2,094,568,126 | 2,095,003,727 | 5,700 | 1,343 |

**Table S4** Genome assembly completeness evaluation by BUSCO.

| **C:88.4%, S:72.0%, D:16.4%, F:2.8%, M:8.8%, n:1440** | |
| --- | --- |
| **Type** | **Percentage (%)** |
| **Complete BUSCOs (C)** | 88.4 |
| **Complete and single-copy BUSCOs (S)** | 72.0 |
| **Complete and duplicated BUSCOs (D)** | 16.4 |
| **Fragmented BUSCOs (F)** | 2.8 |
| **Missing BUSCOs (M)** | 8.8 |

**Table S5** Genome assembly completeness evaluation by CEGMA.

| **Description** | **Completely mapped CEGs** | **Completely+ Partially** |
| --- | --- | --- |
| **Number of CEGs present in the assembly** | 233 | 236 |
| **Completeness of the genome (%)** | 93.95 | 95.16 |

**Table S6** The statistical results of reads coverage for the *G.przewalskii* genome.

|  | **Type** | **Percentage (%)** |
| --- | --- | --- |
| **Reads** | Mapping rate (%) | 98.20 |
| **Genome** | Average sequencing depth | 59.67 |
|  | Coverage (%) | 97.71 |
|  | Coverage at least 4X (%) | 97.02 |
|  | Coverage at least 10X (%) | 95.67 |
|  | Coverage at least 20X (%) | 90.65 |

**Table S7** The statistical results of repeat sequence for the *G. przewalskii* genome.

| **Type** | **Repeat Size(bp)** | **Percent of genome (%)** |
| --- | --- | --- |
| **Tandem repeat** | 87,054,001 | 4.16 |
| **Repeatmasker** | 1,175,837,192 | 56.13 |
| **Proteinmask** | 492,641,154 | 23.52 |
| **Total** | 1,290,122,583 | 61.58 |

***:** Total is the non-redundant result of the above categories after removing the overlap between them

**Table S8** Classification of transposable elements (TEs) identified in the *G .przewalskii* genome,
including their length (in bp) and the percent of genome.

| **Type** | **Denovo+Repbase** | | **TE Proteins** | | **Combined TEs** | |
| --- | --- | --- | --- | --- | --- | --- |
|  | **Length(bp)** | **Percent of genome (%)** | **Length(bp)** | **Percent of genome (%)** | **Length(bp)** | **Percent of genome (%)** |
| **DNA** | 1,992,704 | 0.10 | 855,250 | 0.04 | 2,767,653 | 0.13 |
| **LINE** | 1,362,415 | 0.07 | 1,404,935 | 0.07 | 2,607,510 | 0.12 |
| **SINE** | 21,757 | 0.00 | 0 | 0 | 21,757 | 0.00 |
| **LTR** | 1,163,606,690 | 55.54 | 490,381,728 | 23.41 | 1,237,306,717 | 59.06 |
| **Unknown** | 10,267,031 | 0.49 | 0 | 0 | 10,267,031 | 0.49 |
| **Total** | 1,175,837,192 | 56.13 | 492,641,154 | 23.52 | 1,248,165,421 | 59.58 |

*: This result does not include tandem repeat sequences. Combined TEs were the result of combining the two methods and removing redundancy. Total is the non-redundant result of the above categories after removing the overlap between them

**Table S9** Summary of protein-coding genes identification by three sources.

|  | Gene set | number | mean transcript length (bp) | mean CDS length (bp) | mean exons per gene | mean exon length (bp) | mean intron length (bp) |
| --- | --- | --- | --- | --- | --- | --- | --- |
| *De novo* | Augustus | 115,025 | 2,606.84 | 796.62 | 3.17 | 251.33 | 834.38 |
|  | GlimmerHMM | 191,883 | 9,656.40 | 526.57 | 2.65 | 198.71 | 5,533.54 |
|  | SNAP | 82,344 | 7,303.91 | 464.65 | 3.15 | 147.64 | 3,185.13 |
|  | Geneid | 175,802 | 4,052.88 | 613.70 | 3.38 | 181.79 | 1,447.57 |
|  | Genscan | 114,204 | 11,039.74 | 944.91 | 4.94 | 191.46 | 2,565.16 |
| Homolog | *F.tataricum* | 46,509 | 2,695.64 | 876.11 | 3.39 | 258.73 | 762.52 |
|  | *B.vulgaris* | 48,574 | 2,951.57 | 938.54 | 3.58 | 261.92 | 779.23 |
|  | *C.quinoa* | 43,673 | 3,326.41 | 968.10 | 3.77 | 256.82 | 851.49 |
|  | *D.caryophyllus* | 40,027 | 3,472.68 | 1,012.67 | 3.85 | 263.12 | 863.54 |
|  | *F.esculentum* | 39,502 | 2,708.26 | 896.72 | 3.39 | 264.40 | 757.48 |
| RNAseq | PASA | 78,206 | 4,930.98 | 1,029.70 | 5.26 | 195.70 | 915.46 |
|  | Transcripts | 49,596 | 9,508.30 | 1,833.89 | 5.81 | 315.63 | 1,595.46 |
|  | EVM | 123,463 | 2,812.02 | 770.62 | 3.11 | 247.87 | 967.98 |
|  | Pasa-update* | 123,288 | 2,817.87 | 771.07 | 3.10 | 248.64 | 974.14 |
|  | **Final set*** | **51,990** | **4,188.29** | **1,039.87** | **4.23** | **245.92** | **975.22** |

*: these datasets include UTR regions

**Table S10** The statistical results of gene structure of closely related species.

| species | Number genes | mean transcript length (bp) | mean CDS length (bp) | mean exons per genes | mean exon length (bp) | mean intron length (bp) |
| --- | --- | --- | --- | --- | --- | --- |
| *G.przewalskii* | 51,990 | 4,188.29 | 1,039.87 | 4.23 | 245.92 | 975.22 |
| *F.esculentum* | 286,768 | 1,351.68 | 742.47 | 3.06 | 242.75 | 295.93 |
| *C.quinoa* | 44,734 | 4,370.05 | 1,275.31 | 5.46 | 233.75 | 694.53 |
| *F.tataricum* | 34,554 | 2,236.74 | 1,012.80 | 4.58 | 221.19 | 341.98 |
| *B.vulgaris* | 24,045 | 4,709.11 | 1,143.88 | 4.78 | 239.07 | 941.98 |
| *D.caryophyllus* | 43,491 | 2,772.12 | 1,118.16 | 4.74 | 235.73 | 441.84 |

**Table S11** Summary of function annotation for gene.

|  | **Number** | **Percent (%)** |
| --- | --- | --- |
| **Total** | 51,990 | - |
| **Swissprot** | 32,915 | 63.30 |
| **NR** | 48,249 | 92.80 |
| **KEGG** | 38,598 | 74.20 |
| **InterPro** | 44,125 | 84.90 |
| **GO** | 23,286 | 44.80 |
| **Pfam** | 35,094 | 67.50 |
| **Annotated** | 48,670 | 93.60 |
| **Unannotated** | 3,320 | 6.40 |

**Table S12** Summary statistics of non-coding RNAs in the *G. przewalskii* genome.

| **Type** | **Classification** | **Number** | **Average length(bp)** | **Total length(bp)** | **Percent (%)** |
| --- | --- | --- | --- | --- | --- |
| **miRNA** | | 794 | 119.14 | 94,595 | 0.004515 |
| **tRNA** | | 1,435 | 74.82 | 107,363 | 0.005125 |
| **rRNA** | **rRNA** | 380 | 187.69 | 71,321 | 0.003404 |
|  | **18S** | 80 | 464.81 | 37,185 | 0.001775 |
|  | **28S** | 70 | 132.06 | 9,244 | 0.000441 |
|  | **5.8S** | 13 | 155.15 | 2,017 | 0.000096 |
|  | **5S** | 217 | 105.41 | 22,875 | 0.001092 |
| **snRNA** | **snRNA** | 728 | 120.21 | 87,510 | 0.004177 |
|  | **CD-box** | 370 | 104.35 | 38,608 | 0.001843 |
|  | **HACA-box** | 75 | 130.75 | 9,806 | 0.000468 |
|  | **splicing** | 282 | 138.07 | 38,935 | 0.001858 |
|  | **scaRNA** | 1 | 161 | 161 | 0.000008 |
|  | **Unknown** | 0 | 0 | 0 | 0 |

**Table S17** Comparison likelihood of 10 models performed in fastsimcoal2.

| Scenario | # Parameters | Delta likelihood | AIC | AIC’s weight |
| --- | --- | --- | --- | --- |
| Model 1 | 16 | 299996.781 | 32415813.05 | 0 |
| Model 2 | 16 | 257195.316 | 32218711.01 | 0 |
| Model 3 | 20 | 226248.783 | 32076204.96 | 0 |
| Model 4 | 20 | 241800.621 | 32147817.82 | 0 |
| Model 5 | 24 | 215774.43 | 32027970.79 | 0 |
| Model 6 | 24 | 211975.947 | 32010484.12 | 0 |
| Model 7 | **28** | **192135.426** | **31919117.15** | **1** |
| Model 8 | 28 | 197830.687 | 31945348.8 | 0 |
| Model 9 | 30 | 193405.875 | 31921370.61 | 0 |
| Model 10 | 30 | 199405.875 | 31952807.88 | 0 |

note: A bold line indicates the best-supported model.

**Table S18** Estimation of the demographic parameters with 95% highest posterior density (HPD) for the best model (model 7).

| Parameter | Point estimation | 95% CI |
| --- | --- | --- |
| Current population sizes | | |
| N_Tarim_ | 87,193 | 24,647 ~ 135,945 |
| N_Hami_ | 48,802 | 1,426 ~ 75,464 |
| N_Hexi_ | 82,607 | 7,696 ~ 134,113 |
| N_Hybrid_ | 70,359 | 5,846 ~ 130,051 |
| N_Alxa_ | 44,217 | 5,354 ~ 70,556 |
| Historical population size |  |  |
| N_ANC_ | 1,079,003 | 767,223 ~ 1,129,462 |
| N_Hybrid_Alxa_ | 88,476 | 751 ~ 562,800 |
| N_Hexi_ Alxa_ | 83,184 | 14,297 ~ 1,095,167 |
| N_Hexi_Hami_ | 601,142 | 152,184 ~ 1,131,964 |
| N_Tarim_ | 25,239,928 | 1,717,836 ~ 36,909,467 |
| N_Hybrid_ | 390,563 | 3,942 ~ 684,193 |
| N_Hami_ | 14,126,810 | 71,067 ~ 19,912,634 |
| N_Alxa_ | 12,799,581 | 224,892~18,408,982 |
| N_Hexi_ | 23,912,410 | 1,076,296 ~ 35,892,421 |
| Divergence times (ya) |  |  |
| T_Tarim_ | 5,224,220 | 2,902,460 ~ 5,284,340 |
| T_Hami_ | 2,749,000 | 1,201,515 ~ 2,717,535 |
| T_Hexi_ | 1,424,755 | 132,545 ~ 1,441,165 |
| T_Hybrid_Alxa_ | 742,810 | 925 ~ 763,005 |
| [Contract](javascript:;) times (ya) |  |  |
| T_Contract_ | 56,575 | 545 ~ 78,570 |
| Recent migration rate (10^-6^) |  |  |
| m_Hexi_Hami_ | 3.86 | 1.30 ~ 224 |
| m_Hami_Hexi_ | 26.10 | 3.09 ~ 225 |
| m_Hybrid_Hexi_ | 7.34 | 1.50 ~ 303 |
| m_Hexi_Hybrid_ | 107 | 7.32 ~ 997 |
| m_Alxa_Hybrid_ | 50.10 | 3.34 ~ 873 |
| m_Hybrid_Alxa_ | 62.8 | 3.56 ~ 1060 |
| Ancient migration rate (10^-6^) |  |  |
| m_(Hybrid_Alxa)_Hexi_ | 18.30 | 2.03 ~ 90.8 |
| m_Hexi_(Hybrid_Alxa)_ | 51.3 | 3.85 ~ 636 |
| m_(Hexi_Hybrid_Alxa)_Hami_ | 6.13 | 1.44 ~ 117 |
| m_Hami_(Hexi_Hybrid_Alxa)_ | 364 | 2.04 ~ 544 |
| m_(Hami_Hexi_Hybrid_Alxa)_Tarim_ | 5.79 | 2.91 ~ 220 |
| m_Tarim_(Hami_Hexi_Hybrid_Alxa)_ | 212 | 3.90 ~ 694 |
