## supplemental method for "Global and local adaptation to aridity in a desert plant *Gymnocarpos przewalskii*"

### Supplementary text :

Ornstein Uhlenbeck process to simulate gene number shared by two species

J Chen

Last update: 2022-04-21

#### Contents

##### functions

```
## OU prorcess
simul<-function(alpha0 = 10^(-10), alpha1=10^(-10), sigma = 1, x0 = 0.05, nsim= 20000,
                theta0=0, theta1=0){
   # alpha: stabilizing selection (a0~0, a1 selection),
   # sigma: brownian proces (drift),
   # x0: time to tips, nsim: number of genes,
   # theta: optimal value of positive selection (theta0=0, theta1 selectoin)

   # ancestral nodes
   node1 = rcOU(n = nsim, Dt = 1-0.91, x0=0, theta=c(theta0, alpha0, sigma))
   node2 = rcOU(n = nsim, Dt = 1-0.78, x0=node1, theta=c(theta0, alpha0, sigma))
   node3 = rcOU(n = nsim, Dt = 1-0.71, x0=node2, theta=c(theta0, alpha0, sigma))
   node4 = rcOU(n = nsim, Dt = 1-0.69, x0=node3, theta=c(theta0, alpha0, sigma))
   node5 = rcOU(n = nsim, Dt = 1-0.48, x0=node4, theta=c(theta0, alpha0, sigma))
   node6 = rcOU(n = nsim, Dt = 1-0.3,  x0=node2, theta=c(theta0, alpha0, sigma))
   node7 = rcOU(n = nsim, Dt = 1-0.26, x0=node6, theta=c(theta0, alpha0, sigma))
   node8 = rcOU(n = nsim, Dt = 1-0.8,  x0=0,     theta=c(theta0, alpha0, sigma))
   node9 = rcOU(n = nsim, Dt = 1-0.69, x0=node8, theta=c(theta0, alpha0, sigma))
   node10 = rcOU(n = nsim, Dt = 1-0.53, x0=node9, theta=c(theta0, alpha0, sigma))
   node11 = rcOU(n = nsim, Dt = 1-0.46, x0=node10, theta=c(theta0, alpha0, sigma))
   node12 = rcOU(n = nsim, Dt = 1-0.35, x0=node11, theta=c(theta0, alpha0, sigma))
   node13 = rcOU(n = nsim, Dt = 1-0.26, x0=node11, theta=c(theta0, alpha0, sigma))

   # xerophyte species
   Peu = rcOU(n = nsim, Dt = 0.48, x0=node5, theta=c(theta1, alpha1, sigma))
   Rco = rcOU(n = nsim, Dt = 0.48, x0=node5, theta=c(theta1, alpha1, sigma))
   Pve = rcOU(n = nsim, Dt = 0.69, x0=node4, theta=c(theta1, alpha1, sigma))
   Pco = rcOU(n = nsim, Dt = 0.71, x0=node3, theta=c(theta1, alpha1, sigma))
   Pac = rcOU(n = nsim, Dt = 0.26, x0=node7, theta=c(theta1, alpha1, sigma))
   Car = rcOU(n = nsim, Dt = 0.26, x0=node7, theta=c(theta1, alpha1, sigma))
   Ana = rcOU(n = nsim, Dt = 0.3,  x0=node6, theta=c(theta1, alpha1, sigma))
   Lru = rcOU(n = nsim, Dt = 0.91, x0=node1, theta=c(theta1, alpha1, sigma))

   Sch = rcOU(n = nsim, Dt = 0.69, x0=node9, theta=c(theta1, alpha1, sigma))
   Hun = rcOU(n = nsim, Dt = 0.53, x0=node10, theta=c(theta1, alpha1, sigma))
   Gpr = rcOU(n = nsim, Dt = 0.35, x0=node12, theta=c(theta1, alpha1, sigma))
```

```
   # non-xerophyte species
   Fta = rcOU(n = nsim, Dt = 0.8, x0=node8, theta=c(theta0, alpha0, sigma))
   Dca = rcOU(n = nsim, Dt = 0.35, x0=node12, theta=c(theta0, alpha0, sigma))
   Sol = rcOU(n = nsim, Dt = 0.26, x0=node13, theta=c(theta0, alpha0, sigma))
   Bvu = rcOU(n = nsim, Dt = 0.26, x0=node13, theta=c(theta0, alpha0, sigma))

   return(list("Peu"=Peu, "Rco"=Rco, "Pve"=Pve, "Pco"=Pco, "Pac"=Pac, "Car"=Car,
               "Ana"=Ana, "Lru"=Lru, "Sch"=Sch, "Hun"=Hun, "Gpr"=Gpr,"Fta"=Fta,
               "Dca"=Dca, "Sol"=Sol, "Bvu"=Bvu))
}

## calculate the difference between two species
Dstat <- function(x,y){
## simple model to calculate gene difference between two species, diff<0 similar enough to be in the sa
    diff= abs(x-y)
    if(is.na(diff)){
        return(NA)
    }else if(diff>=0.5){ # quite arbitary cutoff
        return(0)
    }else{
        return(1)
    }
}
## count shared genes between pairs of species in difference LHT classes

get_shared_genes=function(dataset){
    num.spec = length(dataset)
    group1 = c(rep('Xero',11), rep('NonX',4))
    group2 = c(rep('OX',8), rep('CX',3), rep('CNX',4))
    k=choose(num.spec, 2)
    shared_genes=numeric(k)
    comb1 = character(k)
    comb2 = character(k)
    ord = character(k)
    m=1
    for(i in 1:(num.spec-1)){
        for(j in (i+1):num.spec){
            shared_genes[m] = sum(mapply(Dstat, x=dataset[[i]],y=dataset[[j]]))
            comb1[m] = paste(group1[i], group1[j], sep='-')
            comb2[m] = paste(group2[i], group2[j], sep='-')
            ord[m] = paste(i,j,sep='-')
            m = m + 1
     }
    }
    res = data.frame(sharedGenes=shared_genes, group1= comb1, group2=comb2)
    res$group2[res$group2=='CNX-CX'] = 'CX-CNX'
    res$group2[res$group2=='CNX-OX'] = 'OX-CNX'
    res$group2[res$group2=='CX-OX'] = 'OX-CX'
    res$ord = ord
    return(res)
}

## plot simulated and observed data
```

```
plot_sim=function(obs, sim){
    data=rbind(sim[,c('sharedGenes', 'group1','group2')], obs[,c('sharedGenes', 'group1','group2')])
    data$cate=rep(c('sim','obs'),each=105)

    data$genes_tf1 = data$sharedGenes
    # data$genes_tf2 = data$sharedGenes
    xv = sim$sharedGenes - mean(sim$sharedGenes)
    yv = obs$sharedGenes - mean(obs$sharedGenes)
    se = sqrt(sum((yv-xv)^2))/104
    data$genes_tf1 = c(xv, yv)
    # data$genes_tf1[data$cate=='sim']=data$genes_tf1[data$cate=='sim'] - mean(data$genes_tf1[data$cate
    # data$genes_tf1[data$cate=='obs']=data$genes_tf1[data$cate=='obs'] - mean(data$genes_tf1[data$cate

    # data$genes_tf2[data$cate=='sim']=data$genes_tf2[data$cate=='sim']- mean(data$genes_tf2[data$cate=
    # data$genes_tf2[data$cate=='obs']=data$genes_tf2[data$cate=='obs']- mean(data$genes_tf2[data$cate=

    data$group1=factor(data$group1, levels=c('Xero-Xero', 'Xero-NonX', 'NonX-NonX'))
    data$group2=factor(data$group2, levels=c('CNX-CNX', 'CX-CNX', 'OX-CNX','CX-CX','OX-CX','OX-OX'))

    p1<-ggplot(data, aes(colour=factor(cate),y=genes_tf1, x=group1))+geom_boxplot()+xlab('')+ylab('#sha
    p2<-ggplot(data, aes(colour=factor(cate),y=genes_tf1, x=group2))+geom_boxplot()+xlab('')+ylab('#sha
    grid.arrange(p1, p2, ncol=2)

    # return(se)
}

# plot observed, null model, purifying model and full model
plot_sim2=
function(obs, sim1,sim2,sim3){
    data=rbind(sim1[,c('sharedGenes', 'group1','group2')],
               sim2[,c('sharedGenes', 'group1','group2')],
               sim3[,c('sharedGenes', 'group1','group2')],
               obs[,c('sharedGenes', 'group1','group2')])
    #
    data$cate=factor(rep(c('null','neg','full','obs'),each=105), levels=c('obs','null','neg','full'))


    yv = obs$sharedGenes - mean(obs$sharedGenes)
    xv1 = sim1$sharedGenes - mean(sim1$sharedGenes)
    xv2 = sim2$sharedGenes - mean(sim2$sharedGenes)
    xv3 = sim3$sharedGenes - mean(sim3$sharedGenes)
    se1 = sqrt(sum((yv-xv1)^2))/104
    se2 = sqrt(sum((yv-xv2)^2))/104
    se3 = sqrt(sum((yv-xv3)^2))/104

    data$genes_tf1 = c(xv1,xv2,xv3,yv)
    data$group1=factor(data$group1, levels=c('Xero-Xero', 'Xero-NonX', 'NonX-NonX'))
    data$group2=factor(data$group2, levels=c('CNX-CNX', 'CX-CNX', 'OX-CNX','CX-CX','OX-CX','OX-OX'))

    p1<-ggplot(data, aes(colour=cate,y=genes_tf1, x=group1))+geom_boxplot()+xlab('')+ylab('#shared gene
    p2<-ggplot(data, aes(colour=cate,y=genes_tf1, x=group2))+geom_boxplot()+xlab('')+ylab('#shared gene
    grid.arrange(p1, p2, ncol=2)
```

```
    # return(c(se1,se2,se3))
}

#### calculate sum stats for abc inference
summary_sim=function(x, group, FUN=mean){
    tapply(x, factor(group), FUN)
}

read_simulations=function(files, num.lines){
    n = length(files)
    res = NULL
    for(i in 1:n){
        dat = fread(files[i], header=T)
        res = rbind(res, dat[1:num.lines, ])
    }
    return(res)
}

read_prior = function(file, n=1e4, num.line){
    prior = read.table(file,header=T)
    res = NULL
    for(i in 1:20){
        prior.index = prior[((i-1)*n+1):(i*n), ]
        prior.index = prior.index[1:m,]
        res = rbind(res, prior.index)

    }
    return(res)
}

quiet=function (..., messages = FALSE, cat = FALSE)
{
    if (!cat) {
        sink(tempfile())
        on.exit(sink())
    }
    out <- if (messages)
        eval(...)
    else suppressMessages(eval(...))
    # out
}
```

We selected 11 xerophyte plants including 3 in Caryophyllales and 8 out of Caryophyllales to compare with 4 non-xerophyte plants (with broad range of water supply) to test the correlation of number of shared genes to drift effect (phylogenetic branch length) and selection (within similar drought environment).

read and plot tree

```
tree<-read.tree(file='~/Desktop/ZJU/Gymnocarpos/correlation/abc/Rmd/cafemcmc.tree')
plot(tree)
nodelabels(c(0,0:11,13,12))
```

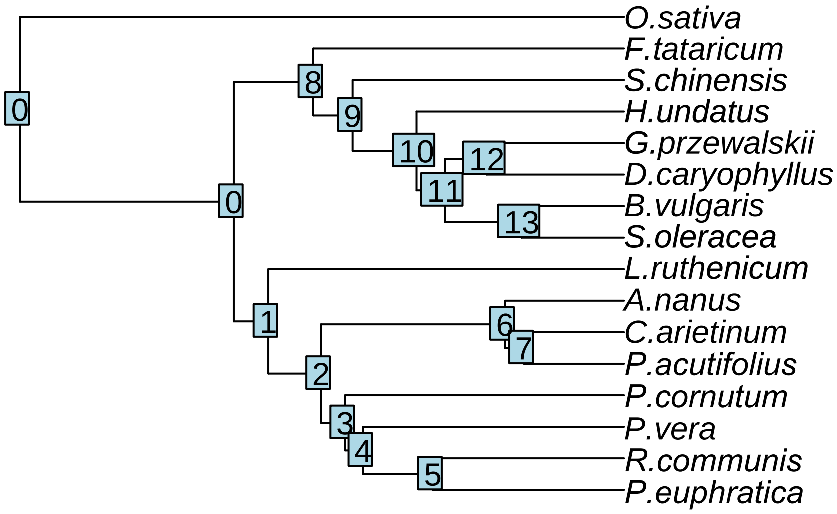


The observation (1 xerophyte vs xerophyte, 2 xerophyte vs non-xerophyte 3 non vs non)

```
obs<-read.table("~/Desktop/ZJU/Gymnocarpos/correlation/abc/Rmd/pairspecies_overlap_genefamily_final_orde
xyplot(sharedGenes~branch | group1,data =obs)
```

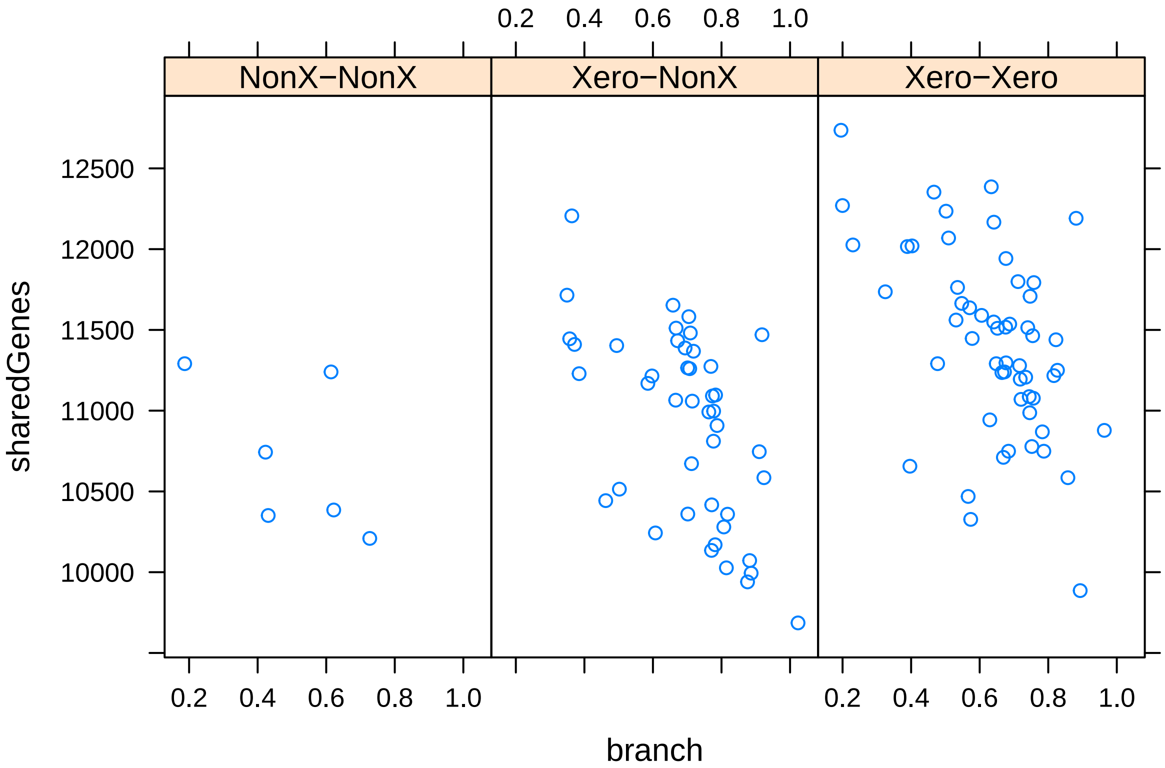


```
# it seems the effect of branch length is not random
# use negative binomial dist to model count data
m1<-glm.nb(obs$sharedGenes ~ obs$branch)
m2<-glm.nb(obs$sharedGenes ~ obs$branch + obs$group1)
anova(m1,m2)
```

```
## Likelihood ratio tests of Negative Binomial Models
##
## Response: obs$sharedGenes
```

```
##                      Model    theta Resid. df    2 x log-lik.   Test    df
## 1                obs$branch 426.9552       103       -1623.057
## 2 obs$branch + obs$group1 561.5540       101       -1595.496 1 vs 2     2
##   LR stat.      Pr(Chi)
## 1
## 2 27.56088 1.035692e-06
```

```
# also significant effect of life history traits (group)
summary(m2)
```

```
##
## Call:
## glm.nb(formula = obs$sharedGenes ~ obs$branch + obs$group1, init.theta = 561.553952,
##     link = log)
##
## Deviance Residuals:
##     Min       1Q   Median       3Q      Max
## -2.5075  -0.5855   0.1081   0.6712   2.4796
##
## Coefficients:
##                      Estimate Std. Error z value Pr(>|z|)
## (Intercept)           9.35917    0.02176 430.086  < 2e-16 ***
## obs$branch           -0.16200    0.02537  -6.385 1.71e-10 ***
## obs$group1Xero-NonX  0.05112    0.01949   2.622  0.00873 **
## obs$group1Xero-Xero  0.08676    0.01893   4.582 4.60e-06 ***
## ---
## Signif. codes:  0 '***' 0.001 '**' 0.01 '*' 0.05 '.' 0.1 ' ' 1
##
## (Dispersion parameter for Negative Binomial(561.554) family taken to be 1)
##
##     Null deviance: 178.65  on 104  degrees of freedom
## Residual deviance: 105.07  on 101  degrees of freedom
## AIC: 1605.5
##
## Number of Fisher Scoring iterations: 1
##
##
##               Theta:  561.6
##           Std. Err.:  81.4
##
##  2 x log-likelihood:  -1595.496
```

```
rsq(m1,adj=T) # variance explained by branch length, a pseduo R2
```

```
## [1] 0.2319826
```

```
rsq(m2,adj=T) # by both branch length and selection
```

```
## [1] 0.4019365
```

```
#we can also test for random effect of branch length
m4=lme(sharedGenes~group1, random=~1|branch, data=obs)
anova.lme(m4)
```

```
##             numDF denDF  F-value p-value
## (Intercept)     1   102 39577.16  <.0001
## group1          2   102    11.48  <.0001
```

```r
summary(m4)
```

```
## Linear mixed-effects model fit by REML
##   Data: obs
##        AIC      BIC    logLik
##   1605.367 1618.492 -797.6835
##
## Random effects:
##  Formula: ~1 | branch
##         (Intercept) Residual
## StdDev:    538.4654 201.9245
##
## Fixed effects:  sharedGenes ~ group1
##                     Value Std.Error  DF  t-value p-value
## (Intercept)     10703.167  234.7759 102 45.58885  0.0000
## group1Xero-NonX   209.106  250.2720 102  0.83552  0.4054
## group1Xero-Xero   714.306  247.2505 102  2.88900  0.0047
##  Correlation:
##                 (Intr) g1X-NX
## group1Xero-NonX -0.938
## group1Xero-Xero -0.950  0.891
##
## Standardized Within-Group Residuals:
##         Min          Q1         Med          Q3         Max
## -0.93506082 -0.26283079  0.04867853  0.27825028  0.80504417
##
## Number of Observations: 105
## Number of Groups: 105
```

```r
# it seems selection effect is still signficant and we found more shared genes in xerophyte paris

# more visualization
ggplot(obs, aes(y= sharedGenes, x=factor(group1,levels=c('Xero-Xero', 'Xero-NonX', 'NonX-NonX')),fill=g
geom_boxplot(outlier.colour="black", outlier.shape=16,outlier.size=1, notch=FALSE,position=position_dodg
xlab('')+ylab('#shared genes')+geom_dotplot(binaxis='y', stackdir='center', dotsize=0.5,position=positio
scale_fill_brewer(palette="RdBu") + theme_minimal()+theme(legend.position = "none", axis.title.y = eleme
```

```
## Bin width defaults to 1/30 of the range of the data. Pick better value with `binwidth`.
```

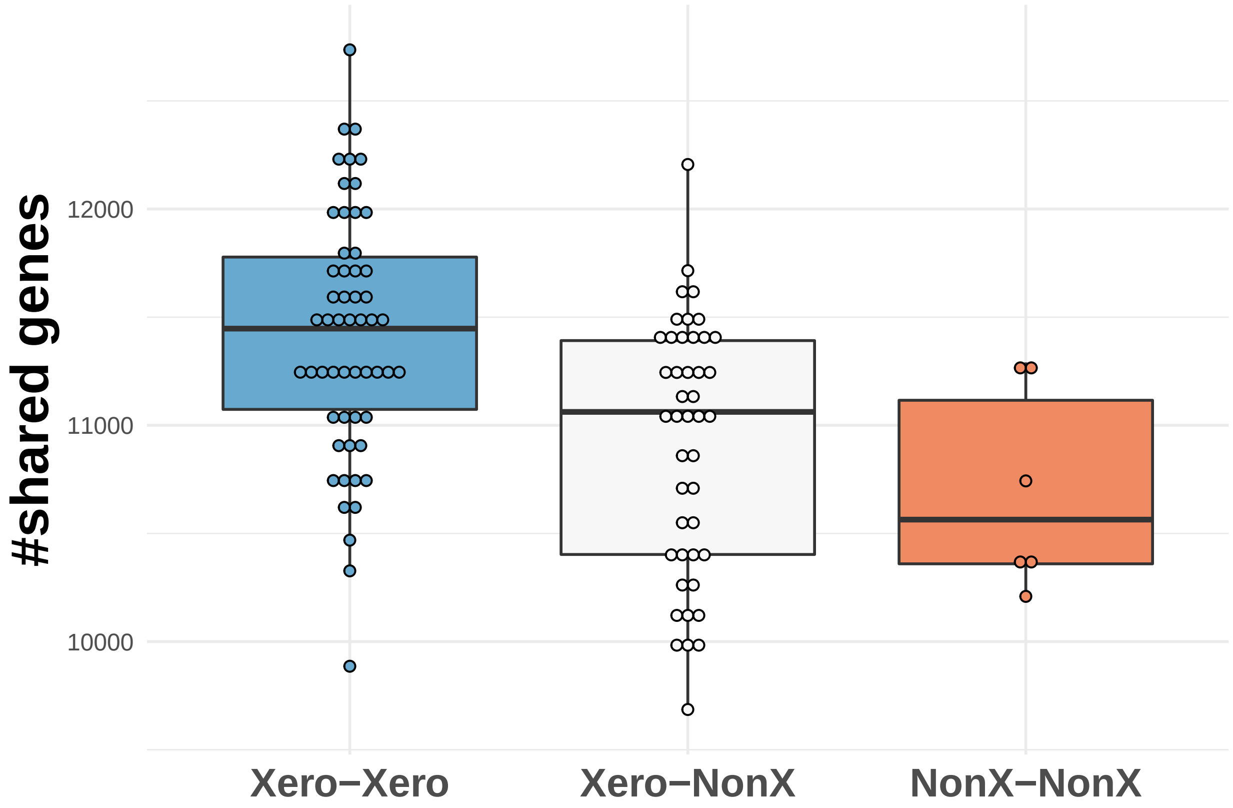


```
ggplot(obs, aes(y= sharedGenes, x=factor(group2, levels=c('CNX-CNX', 'CX-CNX', 'OX-CNX','CX-CX','OX-CX'
geom_boxplot(outlier.colour="black", outlier.shape=16,outlier.size=1, notch=FALSE,position=position_dodg
xlab('')+ylab('#shared genes')+geom_dotplot(binaxis='y', stackdir='center', dotsize=0.5,position=positio
scale_fill_brewer(palette="RdBu") + theme_minimal()+theme(legend.position = "none", axis.title.y = eleme
```

```
## Bin width defaults to 1/30 of the range of the data. Pick better value with `binwidth`.
```

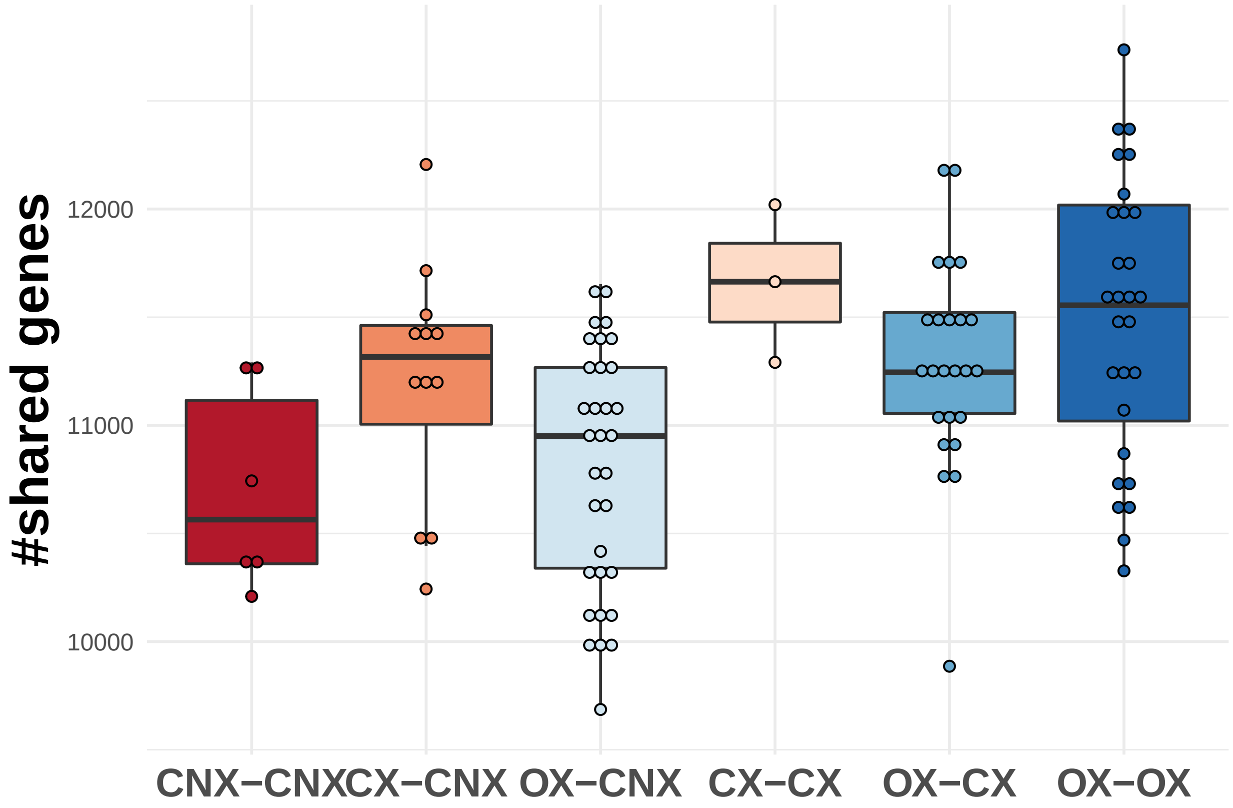


We simulated the data based on an Ornstein-Ulhenbeck process which includes a brownian process with variance sigma, i.e.a drift process with diffusion model(null model). A force moving the trait to the optimum (purifying selection) with the strength alpha and a shift of optimum between species is also allowed (adaptive selection) with the strength theta (full model). We used priors_generator.r to produce prior distributions for all parameters and run the simulations for each draw of parameter settings # e.g. Rscript –vanilla run_simulation_null.r 1 1>log1.txt 2>&1 &

```
###### read observed data

obs<-read.table("~/Desktop/ZJU/Gymnocarpos/correlation/pairspecies_overlap_genefamily_final_order.txt",
group = obs$group2
ss.obs = c(tapply(obs$sharedGenes,group,mean), tapply(obs$sharedGenes,group,sd))
names(ss.obs) = c(paste('mean_', names(ss.obs)[1:6], sep=''), paste('sd_', names(ss.obs)[7:12], sep='')
### read priors
m = 1e4 #  reduce to 1e3 to save the compiling time

## read priors
prior.null = read_prior('~/Desktop/ZJU/Gymnocarpos/correlation/abc/prior/parameters_priors_null.txt', nu
prior.neg = read_prior("~/Desktop/ZJU/Gymnocarpos/correlation/abc/prior/parameters_priors_neg.txt", num
prior.full = read_prior("~/Desktop/ZJU/Gymnocarpos/correlation/abc/prior/parameters_priors_full.txt", nu

## read simulated data
files.null=list.files('~/Desktop/ZJU/Gymnocarpos/correlation/abc/null',pattern='txt$',full=T)
files.null=files.null[c(1,12,14:20,2:11,13)]
sim.null = read_simulations(files.null, m)
names(sim.null) = obs$newOrder

files.neg = list.files('~/Desktop/ZJU/Gymnocarpos/correlation/abc/neg',pattern='txt$',full=T)
files.neg=files.neg[c(1,12,14:20,2:11,13)]
```

```
sim.neg = read_simulations(files.neg, m)
names(sim.neg) = obs$newOrder

files.full = list.files('~/Desktop/ZJU/Gymnocarpos/correlation/abc/full',pattern='txt$',full=T)
files.full = files.full[c(1,12,14:20,2:11,13)]
sim.full = read_simulations(files.full, m)
names(sim.full) = obs$newOrder

## get simulated summary statistics
ss.sim.null = cbind(data.frame(t(apply(sim.null, 1, summary_sim, group, mean))), data.frame(t(apply(sim
names(ss.sim.null) = names(ss.obs)

ss.sim.neg = cbind(data.frame(t(apply(sim.neg, 1, summary_sim, group, mean))), data.frame(t(apply(sim.ne
names(ss.sim.neg) = names(ss.obs)

ss.sim.full = cbind(data.frame(t(apply(sim.full, 1, summary_sim, group, mean))), data.frame(t(apply(sim
names(ss.sim.full) = names(ss.obs)
```

We first used cross validation to test if three models can be distinguished from each other based on simulated summary statistics

```
### model selection
models = c(rep('null', m*20), rep('neg', m*20), rep('full', m*20))
ss.sim = rbind(ss.sim.null, ss.sim.neg, ss.sim.full)
names(ss.sim)=paste('ss',1:12,sep='')
opa = par()
par(mfrow=c(3,4),mar=c(4,2,1,1))
for(i in 1:12){
    boxplot(rbind(ss.sim.null, ss.sim.neg, ss.sim.full)[,i]~models, main=names(ss.obs)[i], ylab='')
    abline(h=ss.obs[i],lty=2,col=2,lwd=2)
}
```

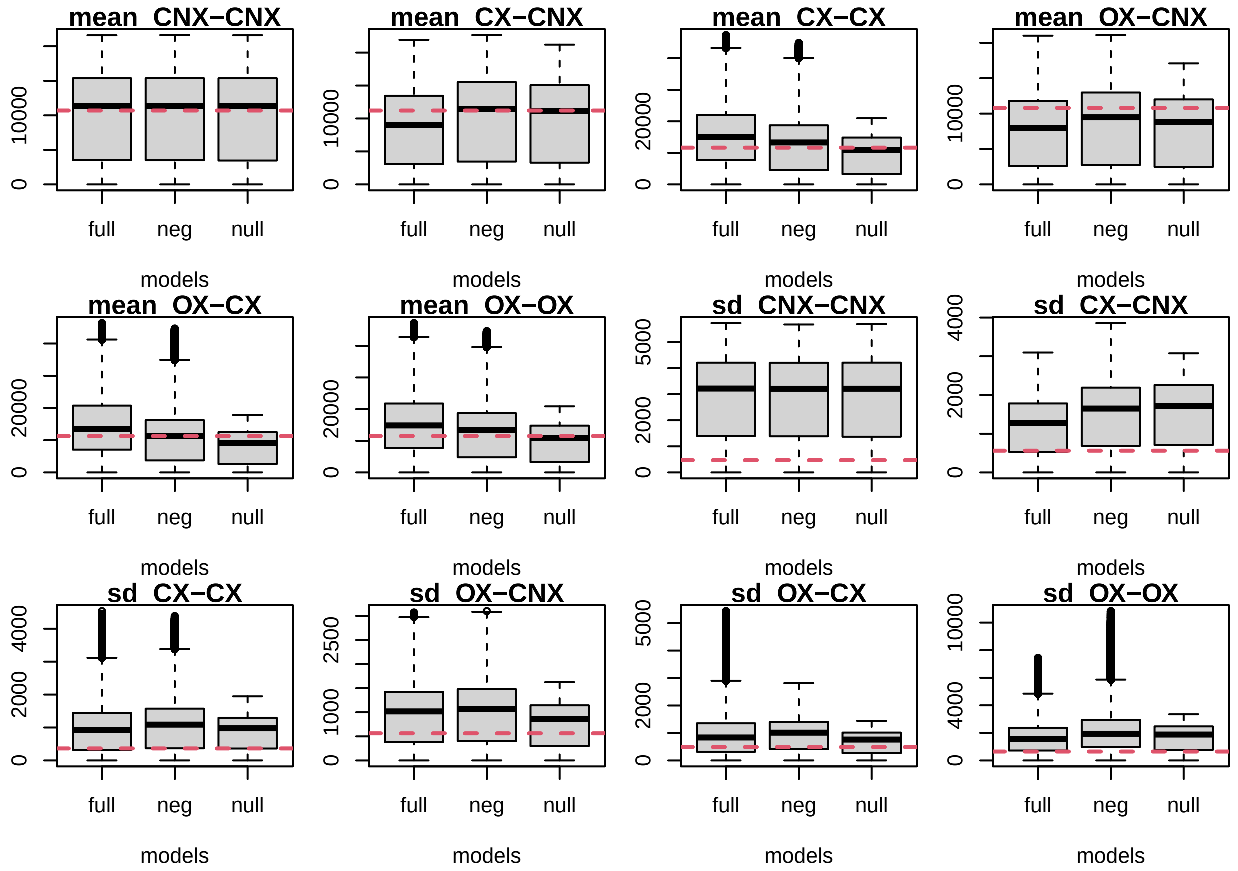


```
cv.modsel.rej <- cv4postpr(models, ss.sim, nval=10, tol=.001, method="rejection")
# cv.modsel.reg <- cv4postpr(models, ss.sim, nval=10, tol=.1, method="mnlogistic") ## we didn't use mnl
summary(cv.modsel.rej)
```

```
## Confusion matrix based on 10 samples for each model.
##
## $tol0.001
##      full neg null
## full    8   2    0
## neg     0  10    0
## null    0   0   10
##
##
## Mean model posterior probabilities (rejection)
##
## $tol0.001
##        full    neg   null
## full 0.7337 0.2638 0.0025
## neg  0.2900 0.6807 0.0293
## null 0.0293 0.0295 0.9412
```

```
par(opa)
```

```
## Warning in par(opa): graphical parameter "cin" cannot be set

## Warning in par(opa): graphical parameter "cra" cannot be set

## Warning in par(opa): graphical parameter "csi" cannot be set
```

```
## Warning in par(opa): graphical parameter "cxy" cannot be set
## Warning in par(opa): graphical parameter "din" cannot be set
## Warning in par(opa): graphical parameter "page" cannot be set
```

```
plot(cv.modsel.rej)
```

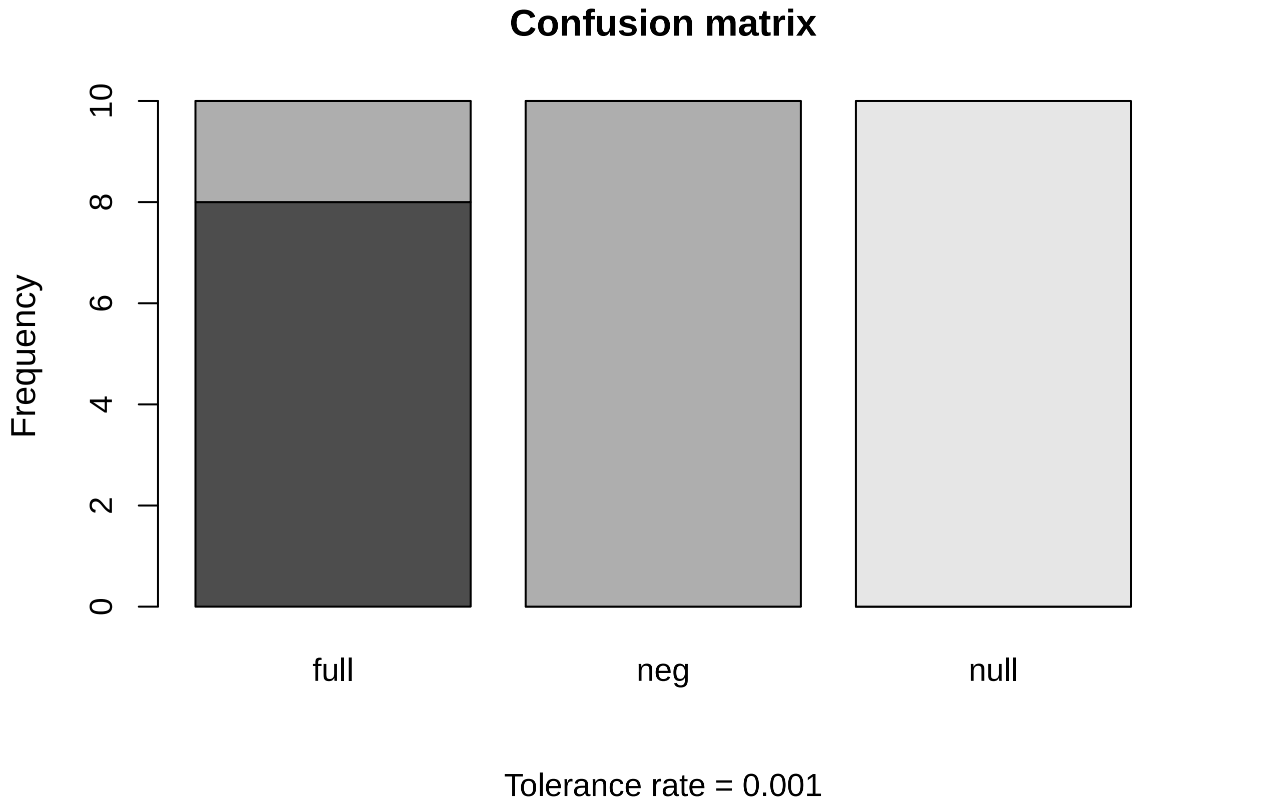


We can see that the results of the null model can be distinguished from other two models with highest probability while the results of the purifying model and the full model can be a bit confused

We then compare three models and choose the best one based on bayes factor

```
modsel.rej <- postpr(ss.obs, models, ss.sim, tol=.01, method="rejection")
modsel.reg <- postpr(ss.obs, models, ss.sim, tol=.01, method="mnlogistic")
summary(modsel.rej)
```

```
## Call:
## postpr(target = ss.obs, index = models, sumstat = ss.sim, tol = 0.01,
##     method = "rejection")
## Data:
##  postpr.out$values (6000 posterior samples)
## Models a priori:
##  full, neg, null
## Models a posteriori:
##  full, neg, null
##
## Proportion of accepted simulations (rejection):
##  full   neg  null
## 0.634 0.348 0.018
##
```

```
## Bayes factors:
##         full     neg    null
## full  1.0000  1.8218 35.2222
## neg   0.5489  1.0000 19.3333
## null  0.0284  0.0517  1.0000
```

```r
summary(modsel.reg)
```

```
## Call:
## postpr(target = ss.obs, index = models, sumstat = ss.sim, tol = 0.01,
##     method = "mnlogistic")
## Data:
##  postpr.out$values (6000 posterior samples)
## Models a priori:
##  full, neg, null
## Models a posteriori:
##  full, neg, null
##
## Proportion of accepted simulations (rejection):
##  full   neg  null
## 0.634 0.348 0.018
##
## Bayes factors:
##         full     neg    null
## full  1.0000  1.8218 35.2222
## neg   0.5489  1.0000 19.3333
## null  0.0284  0.0517  1.0000
##
##
## Posterior model probabilities (mnlogistic):
##   full    neg   null
## 0.3836 0.6164 0.0000
##
## Bayes factors:
##              full          neg         null
## full 1.000000e+00 6.223000e-01 1.442771e+47
## neg  1.606800e+00 1.000000e+00 2.318285e+47
## null 0.000000e+00 0.000000e+00 1.000000e+00
```

We can see that the full model has the largest proportion of accepted points based on both rejection and mnlogistic methods. For both methods the BF supports the full model as the best model of the three

We then used goodness-of-fit test to check the fitness of three models

```r
## GOF
res.gfit.full=gfit(target=ss.obs, sumstat=ss.sim.full, statistic=median, nb.replicate=100)
res.gfit.neg=gfit(target=ss.obs, sumstat=ss.sim.neg, statistic=median, nb.replicate=100)
res.gfit.null=gfit(target=ss.obs, sumstat=ss.sim.null, statistic=median, nb.replicate=100)
plot(res.gfit.full)
```

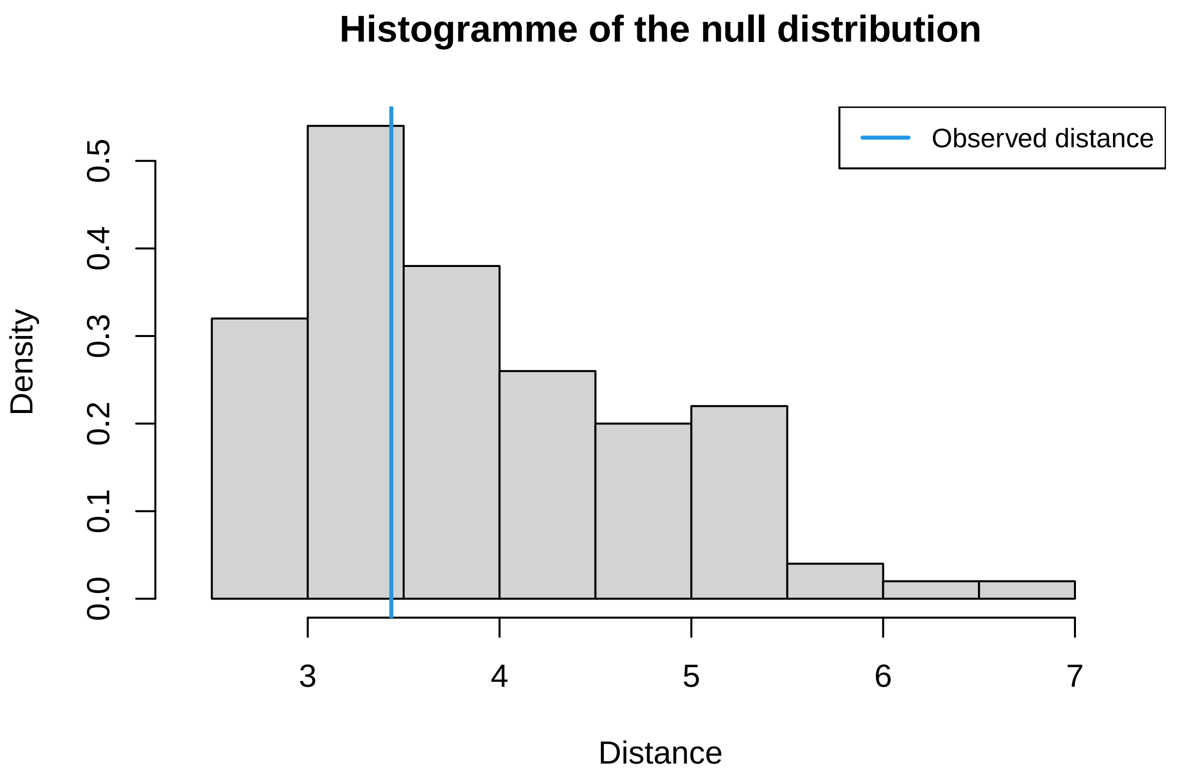


```
plot(res.gfit.neg)
```

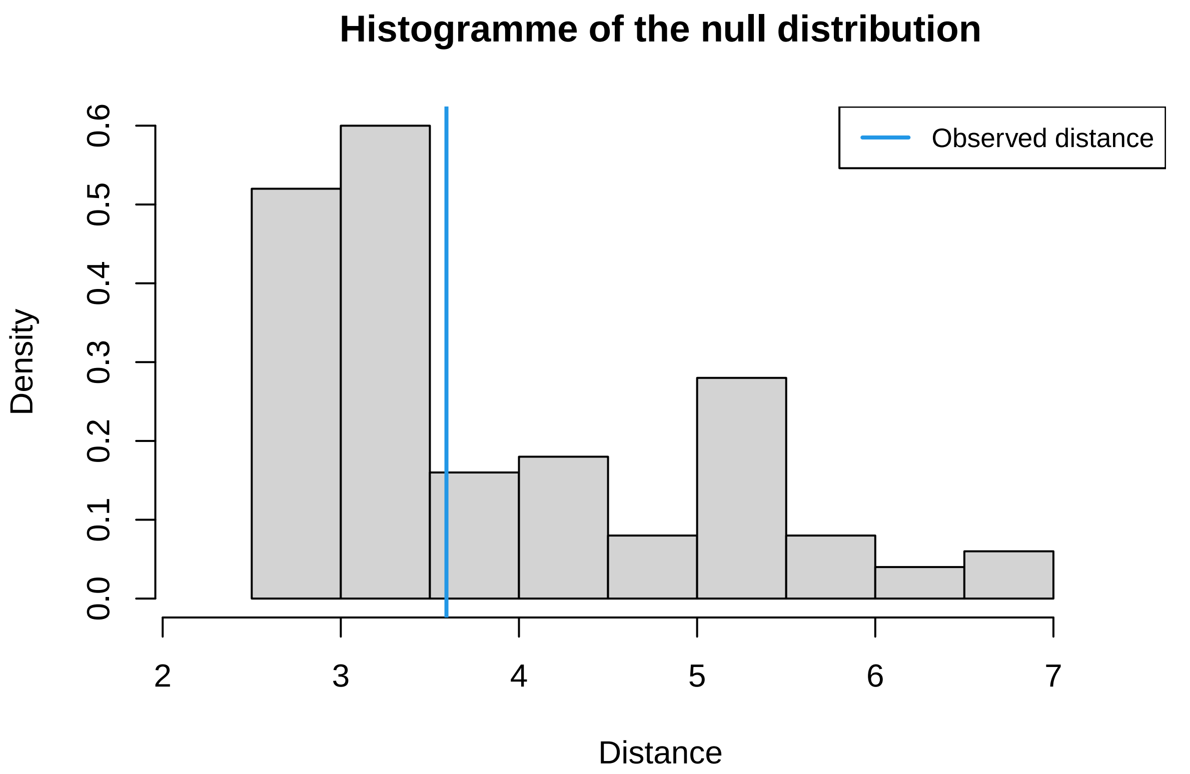


```
plot(res.gfit.null)
```

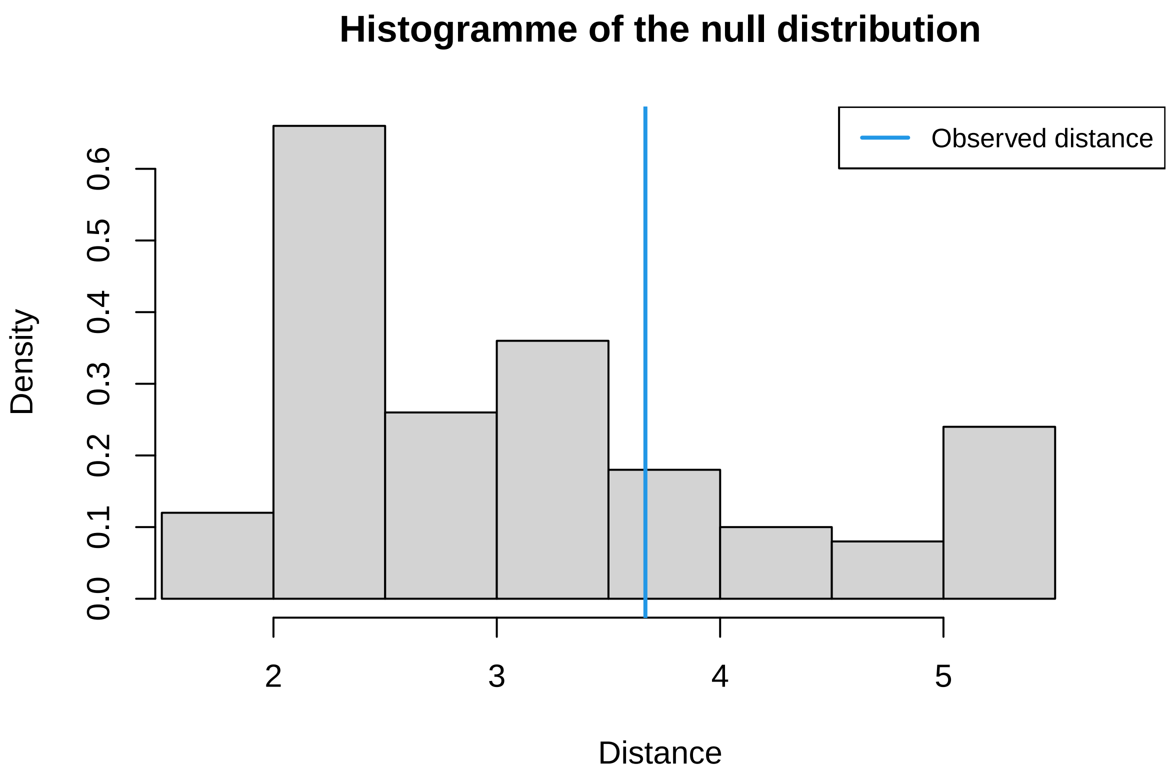


```
gfitpca(target=ss.obs, sumstat=ss.sim, index=models, cprob=.05)
```

```
## Warning in lfproc(x, y, weights = weights, cens = cens, base = base, geth =
## geth, : procv: parameters out of bounds

## Warning in lfproc(x, y, weights = weights, cens = cens, base = base, geth =
## geth, : procv: parameters out of bounds

## Warning in lfproc(x, y, weights = weights, cens = cens, base = base, geth =
## geth, : procv: parameters out of bounds

## Warning in lfproc(x, y, weights = weights, cens = cens, base = base, geth =
## geth, : procv: parameters out of bounds

## Warning in lfproc(x, y, weights = weights, cens = cens, base = base, geth =
## geth, : procv: parameters out of bounds
```

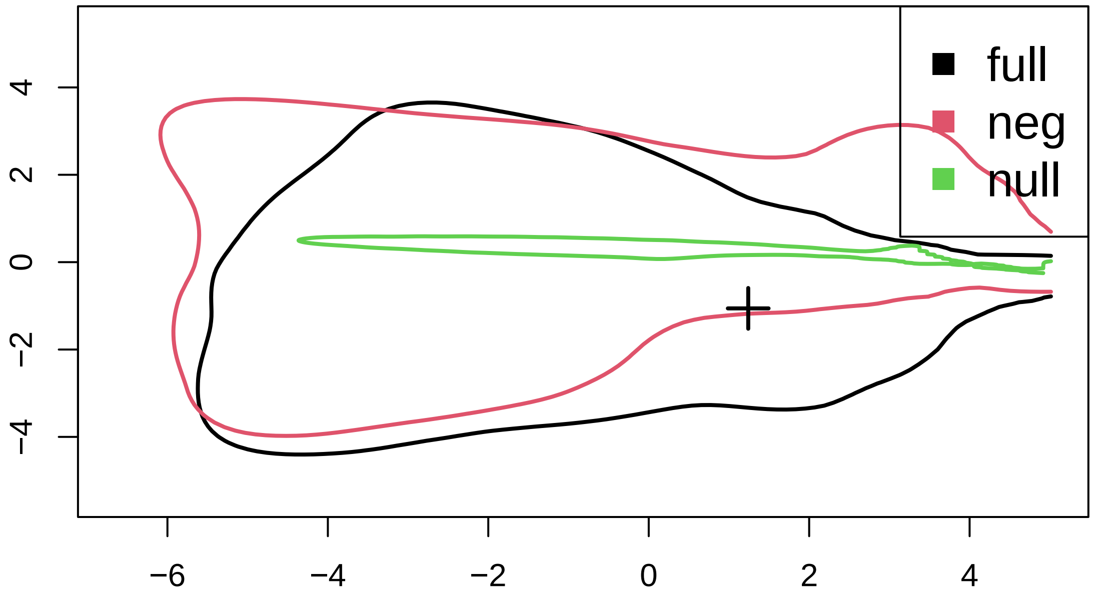


Still the simulated data of full model fits observation better than other two. The null model significantly deviated from the observed data

We then use the cross valiation to see how accurate we can get for parameter estimates at different tolerance rates. We choose the simple rejection method for parameter estimation but one can also choose 'loclinear' and 'neuralnet'.

```
## parameter estimation accuracy
cv.rej.params<- cv4abc(data.frame(num.genes=prior.full[,"num.genes"], sigma=log10(prior.full[,"sigma"])
                ss.sim.full, nval=10, tols=c(.005,.01, 0.05), method="rejection")

summary(cv.rej.params)
```

```
## Prediction error based on a cross-validation sample of 10
```

```
##        num.genes      sigma      alpha      theta
## 0.005 0.19185794 0.05735351 0.01477342 0.53367853
## 0.01  0.18166829 0.07709154 0.02416636 0.59195941
## 0.05  0.34774937 0.16074623 0.04911444 0.80104227
```

```
cv.reg.params<- cv4abc(data.frame(num.genes=prior.full[,"num.genes"], sigma=log10(prior.full[,"sigma"])
                ss.sim.full, nval=10, tols=c(.005,.01, 0.05), method="loclinear")

summary(cv.reg.params)
```

```
## Prediction error based on a cross-validation sample of 10
```

```
##         num.genes       sigma       alpha       theta
## 0.005 0.128405131 0.013127762 0.002014258 0.596903472
## 0.01  0.137437765 0.013278594 0.003645024 0.614346087
## 0.05  0.123093138 0.013419596 0.014595154 0.753521471
```

```
## we skip neuralnet here to avoid tons of screen printing
quiet(cv.neu.params<- cv4abc(data.frame(num.genes=prior.full[,"num.genes"], sigma=log10(prior.full[,"si
                ss.sim.full, nval=10, tols=c(.005,.01, 0.05), method="neuralnet",transf=c('logit','none
summary(cv.neu.params)
```

```
## Prediction error based on a cross-validation sample of 10
```

```
##         num.genes       sigma       alpha       theta
## 0.005 0.335448815 0.010904660 0.001012990 0.136433933
## 0.01  0.341675883 0.011922798 0.001439437 0.124079251
## 0.05  0.342003216 0.018777242 0.001829661 0.167540517
```

Now we do the parameter inference

```
# parameter inference
param.rejection <- abc(target=ss.obs, param=data.frame(num.genes=prior.full[,"num.genes"], sigma=log10(p
                 sumstat=ss.sim.full, tol=0.005, method="rejection")

summary(param.rejection)
```

```
## Call:
## abc(target = ss.obs, param = data.frame(num.genes = prior.full[,
##     "num.genes"], sigma = log10(prior.full[, "sigma"]), alpha = log10(prior.full[,
##     "alpha"]), theta = log10(prior.full[, "theta"])), sumstat = ss.sim.full,
##     tol = 0.005, method = "rejection")
## Data:
##  abc.out$unadj.values (1000 posterior samples)
##
##                num.genes      sigma      alpha      theta
## Min.:         30000.0000    -1.0800    -1.6900    -2.0000
## 2.5% Perc.:   30090.0000    -0.7105    -1.6200    -1.9500
## Median:       32425.0000    -0.0700    -1.3600    -0.8900
## Mean:         33427.4900    -0.1261    -0.7188    -0.8952
## Mode:         30906.2277     0.0025    -1.4292    -0.5994
## 97.5% Perc.: 41420.2500     0.1400     0.8902     0.3302
## Max.:         46580.0000     0.2000     0.9600     0.7100
```

```
par(mfrow=c(2,2))
hist(param.rejection,breaks=40)
```

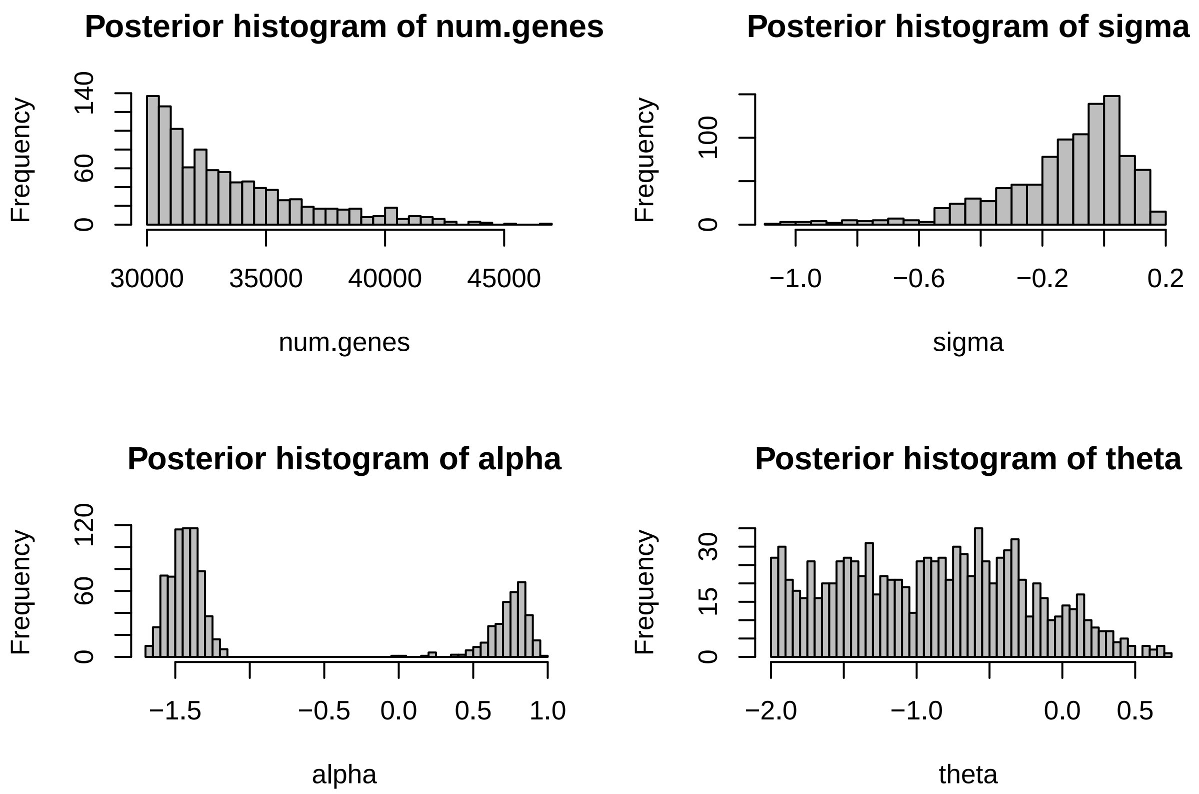


```
param.regress <- abc(target=ss.obs, param=data.frame(num.genes=prior.full[,"num.genes"], sigma=log10(pr
                     sumstat=ss.sim.full, tol=0.005, method="loclinear")
```

```
## Warning: All parameters are "none" transformed.
```

```
summary(param.regress)
```

```
## Call:
## abc(target = ss.obs, param = data.frame(num.genes = prior.full[,
##     "num.genes"], sigma = log10(prior.full[, "sigma"]), alpha = log10(prior.full[,
##     "alpha"]), theta = log10(prior.full[, "theta"])), sumstat = ss.sim.full,
##     tol = 0.005, method = "loclinear")
## Data:
##  abc.out$adj.values (1000 posterior samples)
## Weights:
##  abc.out$weights
##
##                              num.genes        sigma        alpha        theta
## Min.:                     -114202.3420     -629.4961      -0.6253    -158.8346
## Weighted 2.5 % Perc.:      -80631.6256     -217.3040      -0.5288     -47.0194
## Weighted Median:           -31888.9548       15.6246      -0.3849      -3.8179
## Weighted Mean:             -26876.0243        8.3793      -0.3787      -6.2471
## Weighted Mode:             -39069.9300       31.6644      -0.4078       5.9651
## Weighted 97.5 % Perc.:      63371.1070      187.1075      -0.1860      26.3730
## Max.:                      153627.3977      352.6976       0.0813      52.3097
```

```
par(mfrow=c(2,2))
hist(param.regress,breaks=40)
```

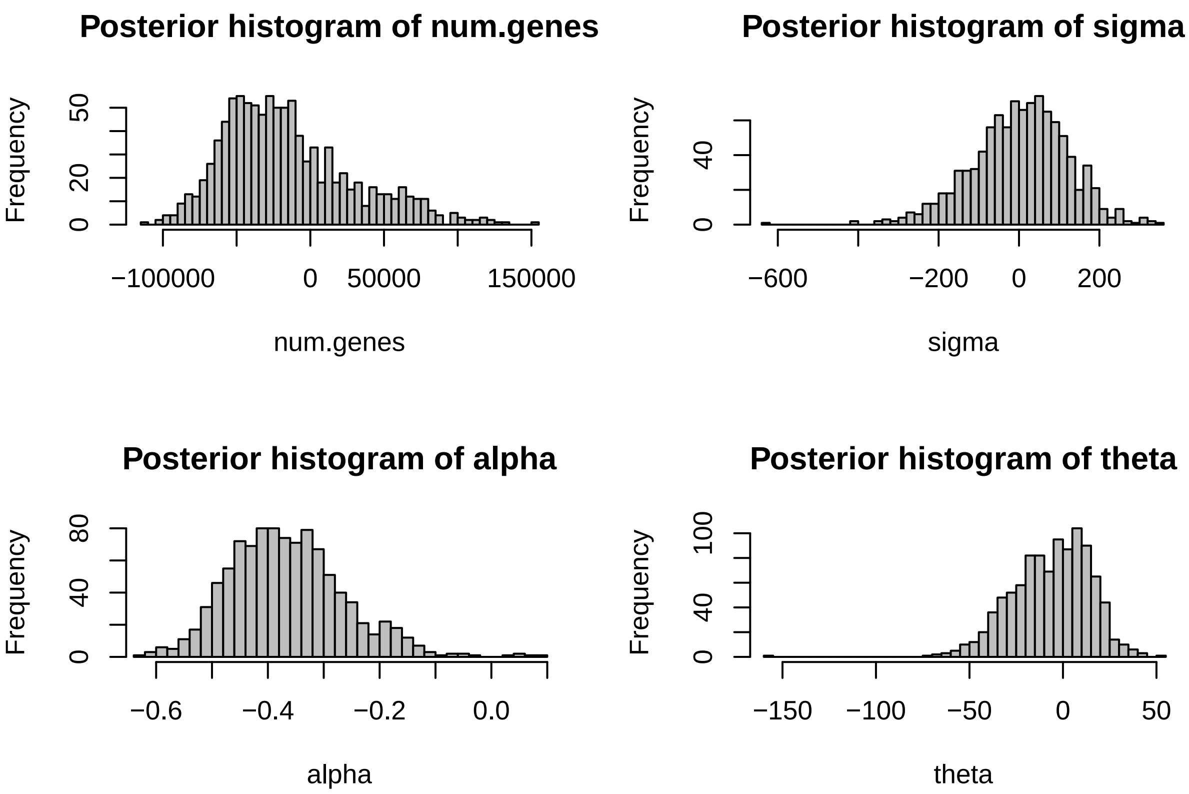


```
param.neu <- abc(target=ss.obs, param=data.frame(num.genes=prior.full[,"num.genes"], sigma=log10(prior.
                 sumstat=ss.sim.full, tol=0.005, method="neuralnet",transf=c('logit','none','none','non
```

```
## 12345678910
## 12345678910
```

```
summary(param.neu)
```

```
## Call:
## abc(target = ss.obs, param = data.frame(num.genes = prior.full[,
##     "num.genes"], sigma = log10(prior.full[, "sigma"]), alpha = log10(prior.full[,
##     "alpha"]), theta = log10(prior.full[, "theta"])), sumstat = ss.sim.full,
##     tol = 0.005, method = "neuralnet", transf = c("logit", "none",
##         "none", "none"), logit.bounds = rbind(c(20000, 60000),
##         c(NA, NA), c(NA, NA), c(NA, NA)))
## Data:
##  abc.out$adj.values (1000 posterior samples)
## Weights:
##  abc.out$weights
##
##                          num.genes     sigma     alpha     theta
## Min.:                   25390.1405   -2.4056   -0.8964   -5.7861
## Weighted 2.5 % Perc.:   25580.2693   -1.4811   -0.8698   -4.5670
## Weighted Median:        25704.1994   -0.7073   -0.8095   -3.5290
## Weighted Mean:          25707.5772   -0.7317   -0.8099   -3.5833
## Weighted Mode:          25692.6104   -0.6405   -0.8042   -3.4226
## Weighted 97.5 % Perc.: 25856.7019   -0.1114   -0.7524   -2.7374
## Max.:                   26002.1312    0.4736   -0.5464   -2.5041
```

```
par(mfrow=c(2,2))
hist(param.neu,breaks=40)
```

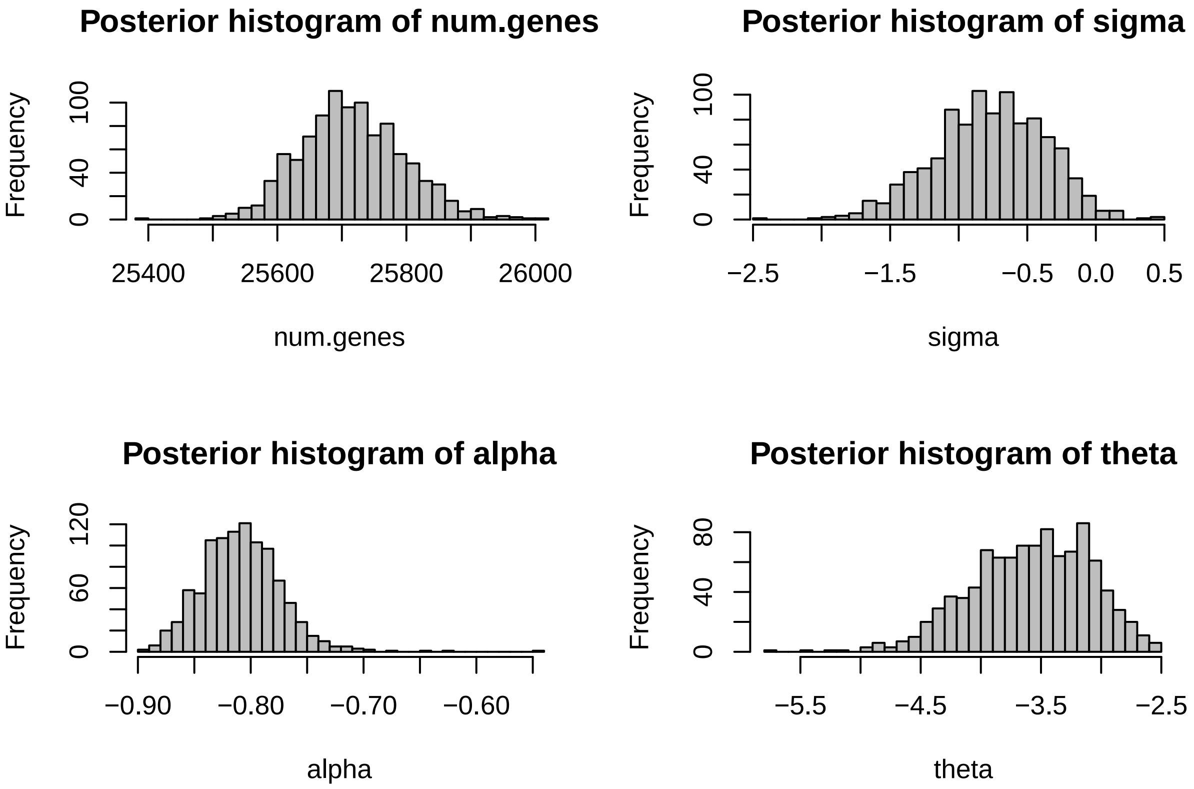


We resampled 1000 points from the accepted posterior distributions of four parameters and run the simulation, recalculate the summary statistic distributions to do the posterior predictive check. We only present the parameters estimated from the simple 'rejection' method (tol = .005) as the regression and neural network didn't give good fit.

```
### postpr check
read_postpr=function(path){
    files = list.files(path, pattern='txt$',full=T)
    n = length(files)
    postpr = NULL
    for(i in 1:n){
        dat = read.table(files[i],header=T)
        postpr = rbind(postpr, dat)
    }
    return(postpr)
}

postpr = read_postpr('~/Desktop/ZJU/Gymnocarpos/correlation/abc/posterior/rejection')
postpr.sim.ss = cbind(data.frame(t(apply(postpr, 1, summary_sim, group, mean))), data.frame(t(apply(pos
names(postpr.sim.ss) = names(ss.obs)
par(mfrow = c(3,4), mar=c(5,2,4,0))
for (i in 1:12){
 hist(postpr.sim.ss[,i],breaks=40, xlim=range(c(postpr.sim.ss[,i], ss.obs[i])), main=names(ss.obs)[i], 
 abline(v=ss.obs[i],lty=2,col=2,lwd=2)
}
```

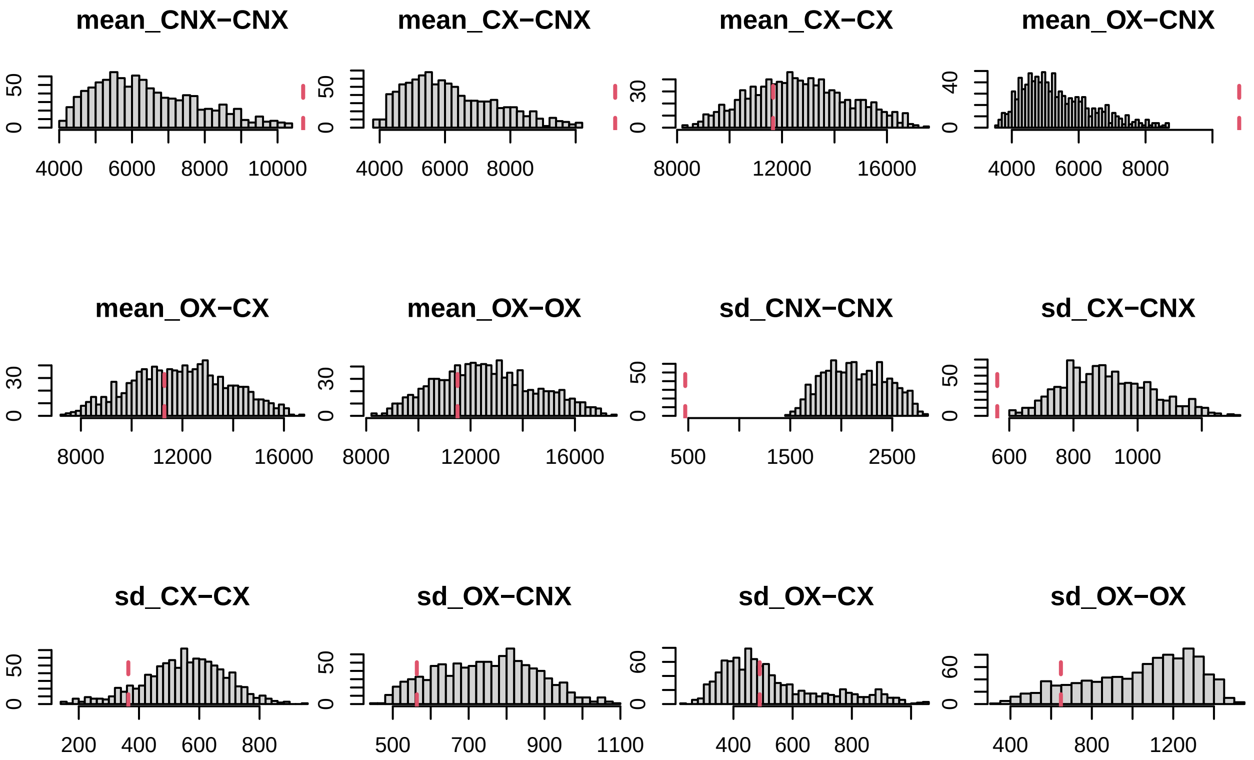


In general, we can conclude that the full model, including the drift effect and both purifying and directional selection, performed significantly better than simple drift model or drift and purifying model. However, it is difficult to gain good estimates of paramters. The could be that our model is still oversimplified with only four parameters and more complicated hidden correlation exists between groups of species other than xerophyte and non-xerophyte or the phylogenetic relationship.
